# Deciphering Spatial Cellular Organization through RNA-Anchored Multi-Omics Deconvolution

**DOI:** 10.64898/2026.10.09.757940

**Authors:** Yihao Bai, Xingbo Guan, Shucun Xiong, Xuan Guo, Xuanwu Wang, Siyu Yi, Wei Ju, Shiyang Ma, Doudou Zhou, Qishi Dong, Wei Liu

## Abstract

Spatial technologies that jointly profile RNA and additional molecular layers provide a comprehensive view of tissue organization, yet their limited resolution results in measurements that reflect mixtures of multiple cell types. While RNA-based deconvolution methods are well established, incorporating co-profiled non-transcriptomic modalities without compromising cell-type identifiability remains an open challenge, as reliable references are largely restricted to transcriptomic data. Here we present SMODER, an RNA-anchored framework for spatial multi-omics deconvolution that treats transcriptomes as the primary determinant of cell identity while incorporating complementary molecular modalities in a constrained manner. By adopting an asymmetric integration strategy, SMODER preserves RNA-driven cell-type resolution and enables robust estimation of cell-type compositions across spatial locations. Across simulations and diverse spatial multi-omics datasets spanning protein, chromatin accessibility and histone modification assays, SMODER consistently outperforms existing methods in both accuracy and spatial coherence. Beyond deconvolution, SMODER denoises multi-omic signals and enables cell-type-aware analysis of regulatory programs. Application to spatial CUT&Tag–RNA-seq and RNA–ATAC-seq data reveals spatially resolved epigenetic repression and cis-regulatory relationships linked to gene expression.

## Introduction

Spatially resolved technologies that jointly profile RNA and complementary molecular layers are rapidly expanding our ability to interrogate tissues *in situ* [1–4]. By integrating protein abundance, chromatin accessibility or histone modification measurements, these approaches provide a multi-layered view of cellular identity and regulatory state within intact tissue architecture [5, 6]. Despite these advances, these technologies remain constrained by limited spatial resolution, whereby each spatial location captures molecular signals aggregated from multiple, potentially heterogeneous cells. As a result, the observed measurements represent convoluted mixtures of cellular identities and molecular states, obscuring both cell-type composition and regulatory organization within tissues. Resolving these spatially mixed cellular and molecular programs has therefore become a central computational challenge for spatial multi-omics analysis.

A large body of computational methods has been developed to address deconvolution using transcriptomic data alone, including RCTD [7], CARD [8], IRIS [9], QR-SIDE [10], Tangram [11], cell2location [12], and STitch3D [13]. These approaches leverage the widespread availability of single-cell RNA sequencing (scRNA-seq) atlases as reference profiles for cell-type deconvolution [14–16], and have demonstrated strong performance in resolving spatial cellular composition from transcriptomic measurements [17, 18]. However, these RNA-only methods are not always sufficient for accurately resolving transcriptionally similar cell types. Many functionally distinct cells, such as naive and memory T cells, exhibit highly similar transcriptional profiles at rest, making them difficult to distinguish based solely on steady-state RNA measurements [19, 20]. Complementary modalities address this gap by providing orthogonal information. Protein abundance identifies cells through lineage-specific surface markers, while chromatin accessibility and histone modifications capture stable regulatory programs beyond transcription. Complementary modalities do more than improve cell-type discrimination; they provide direct measurements of molecular states that define cellular function.

Yet realizing spatial multi-omics deconvolution faces fundamental challenges that remain largely unresolved. While non-transcriptomic modalities, including protein abundance and epigenomic features, provide complementary biological information, they lack comprehensive and standardized reference atlases comparable to scRNA-seq [21]. Moreover, these modalities differ substantially from transcriptomic measurements in noise characteristics, sparsity and biological specificity [20, 22]. As a result, direct integration of multiple spatial omics layers can compromise cell-type identifiability and lead to unstable or confounded inference [23–25]. These observations indicate that spatial multi-omics deconvolution is fundamentally asymmetric across molecular modalities. Rather than treating all modalities equally, effective integration requires a principled framework that anchors cell-type identity in transcriptomic reference profiles while allowing complementary modalities to contribute as spatial calibrators, providing constraints on cell boundaries, technical noise reduction, and orthogonal regulatory signals, without overriding the RNA-defined identity.

To address this challenge, we introduce SMODER (Spatial Multi-Omics Deconvolution anchored by RNA), a principled framework based on an RNA-anchored integration strategy. SMODER treats transcriptomic profiles as the primary determinant of cell identity and incorporates complementary molecular layers to refine spatial inference without overriding RNA-driven signals. By combining spatial modeling with constrained cross-modal information flow, SMODER preserves cell-type identifiability while leveraging additional modalities to reduce noise, enhance spatial coherence and capture regulatory signals.

Beyond accurate estimation of cell-type composition, SMODER enables cell-type-aware analysis of spatial multi-omics data. The framework denoises multi-omic expression maps and supports the identification of regulatory programs linking gene expression with epigenomic features. This capability facilitates the dissection of spatially resolved regulatory programs, including epigenetic repression patterns and cis-regulatory interactions, directly within intact tissues. By unifying spatial deconvolution with RNA-anchored multi-omics integration, SMODER provides a general framework for reconstructing cellular composition and regulatory architecture from spatial multi-omics data.

## Results

### Overview of SMODER

SMODER is a framework for spatial multi-omics deconvolution that quantifies cell-type relative abundance from spatially resolved measurements of RNA and complementary molecular modalities (Fig. 1). In these data, each spatial location typically represents a mixture of multiple cell types due to limited spatial resolution, making the estimation of cell-type abundance a central analytical task.

**Figure 1:**
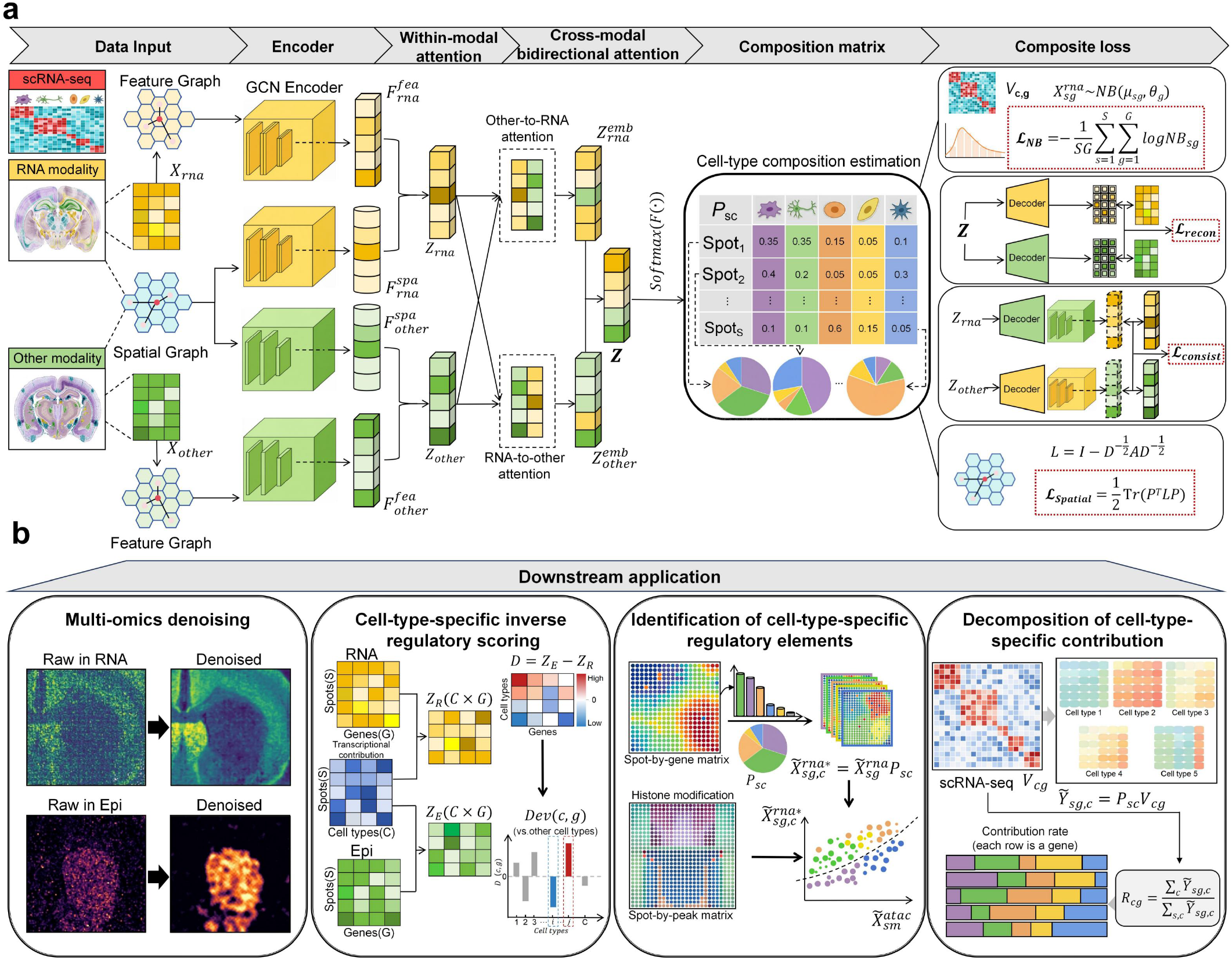
Conceptual overview of SMODER. **a**, Spatial multi-omics technologies jointly measure RNA and additional molecular layer at each spatial location, but limited spatial resolution results in mixed signals from multiple cell types. Cell-type reference information is inherently asymmetric across modalities: comprehensive cell-type atlases are available for transcriptomes, whereas comparable references are largely absent for proteins and epigenomic layers, rendering symmetric multi-modal integration prone to loss of cell-type identifiability. SMODER adopts an RNA-anchored paradigm that treats transcriptomic information as the defining source of cell identity while using non-transcriptomic modalities to refine and regularize spatial inference. SMODER implements this paradigm using a graph-based neural architecture with a constrained bidirectional cross-modal attention mechanism that integrates heterogeneous omics layers while preserving RNA-defined cell-type reference and spatial coherence. *NB*_*sg*_ represents the negative binomial probability mass function 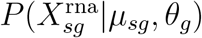. **b**, SMODER enables several downstream applications, including denoising of spatial multi-omic expression maps, scoring of cell-type-specific regulation deviation, identification of cell-type-specific regulatory programs, and decomposition of cell-type-specific expression contribution.

SMODER takes spatial multi-omics data together with a scRNA-seq reference as input (Fig. 1a). The scRNA-seq data define *C* cell types and *G* informative genes, summarized by a cell-type-specific reference basis matrix *V* ∈ ℝ^*C×G*^ that represents the mean expression level of each gene in each cell type. For the spatial multi-omics data, let *X*_rna_ ∈ ℝ^*S×G*^ denote the gene expression matrix measured across *S* spatial locations, and *X*_other_ ∈ ℝ^*S×M*^ represent measurements from other modality (e.g., proteins).

Using these inputs, SMODER anchors cell-type identity in transcriptomic signals while integrating complementary modalities through graph-based bidirectional cross-modal attention. Importantly, this asymmetric design treats RNA as the primary determinant of cell identity, with complementary modalities serving as spatial calibrators that provide constraints on cell boundaries, reduce technical noise, and reveal regulatory variation that transcriptomes alone cannot resolve, without overriding the RNA-defined reference. As a result, SMODER not only improves the quantification of cell-type relative abundance across spatial locations, but also enables downstream analyses of spatial regulatory programs (Fig. 1b).

### RNA anchoring is essential for identifiable spatial multi-omics deconvolution

A central challenge in spatial multi-omics deconvolution is accurately estimating cell-type relative abundance while integrating heterogeneous molecular modalities without distorting the RNA-defined cell identity reference. Transcriptomic measurements benefit from comprehensive single-cell references, whereas non-transcriptomic modalities such as protein abundance, chromatin accessibility and histone modifications lack comparable reference atlases and exhibit distinct noise structures. As a result, treating modalities symmetrically can obscure cell-type signals, particularly under low spatial resolution or sparse measurements.

We generated simulated spatial multi-omics datasets by aggregating cells from matched single-cell multi-omic references, including human melanoma RNA–ATAC data [26] and human central nervous system (CNS) RNA–protein datasets [27]. Each spatial location contained a controlled mixture of cells, with RNA and secondary modalities simulated under empirically calibrated noise (Methods). This design enabled systematic and unbiased comparison of symmetric multi-modal deconvolution (treating RNA and the secondary modality equally), RNA-only deconvolution (i.e., RCTD and CARD) and RNA-anchored deconvolution (SMODER) under identical ground-truth cell-type compositions. We further included non-negative least square regression (NNLS), non-negative matrix factorization (NMF), and SpatialGlue followed by Softmax (SpatialGlue-S) as representative symmetric baselines, where NNLS assumes access to paired references, while NMF and SpatialGlue-S requires post hoc alignment of latent components (Methods).

Across diverse reference-based simulated spatial multi-omics datasets, SMODER achieved accurate and stable estimation of cell-type relative abundance (Fig. 2a–b and Supplementary Figs S1–S3). In contrast, symmetric multi-modal deconvolution approaches, including reference-based NNLS, joint-latent NMF and SpatialGlue-S, produced less stable estimates and increased mixing between distinct cell types. Notably, even NNLS, which explicitly incorporates cell-type reference information, exhibited reduced identifiability when RNA and secondary modalities were treated symmetrically, highlighting the importance of RNA-anchored integration under realistic noise and mixing conditions.

**Figure 2:**
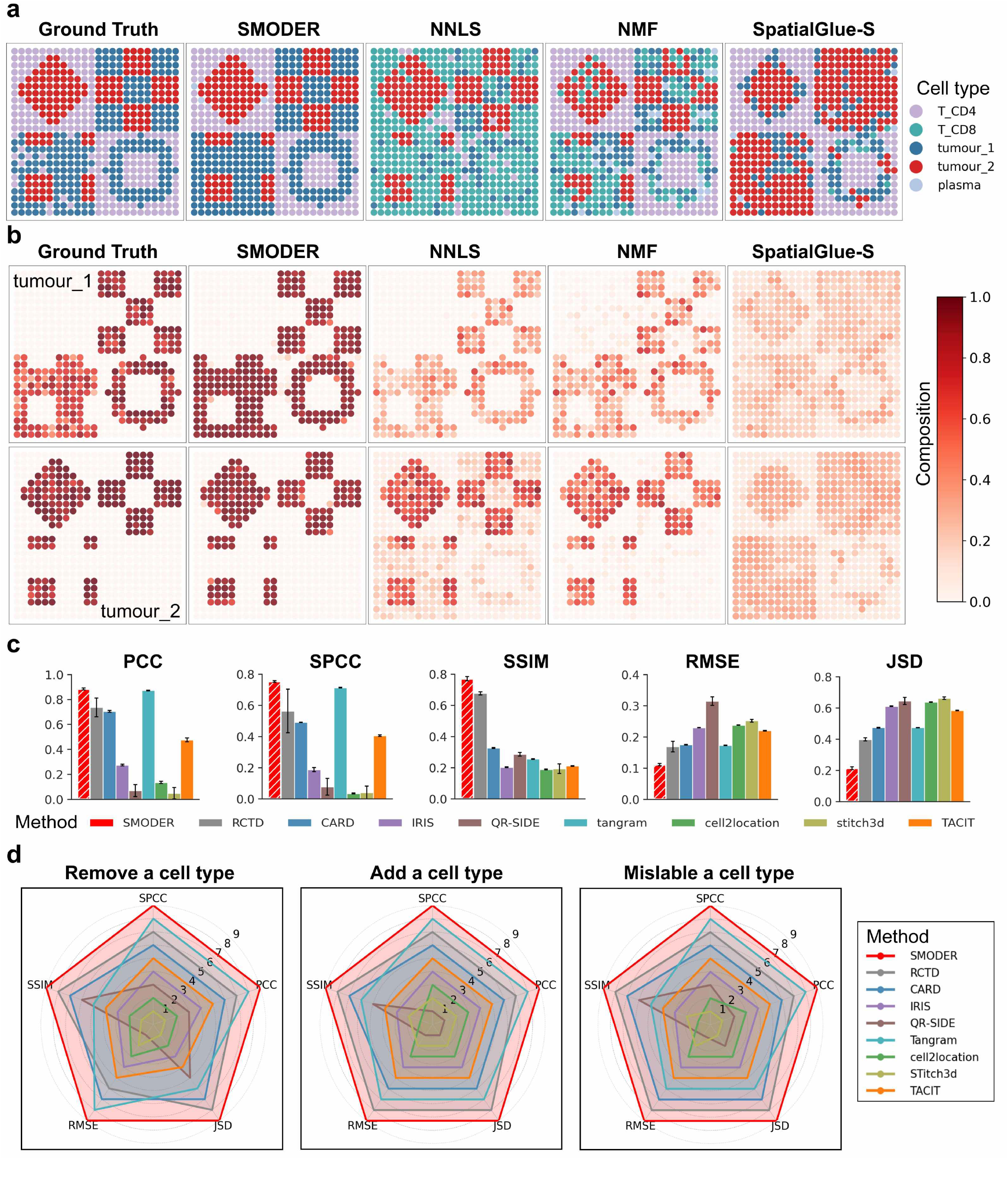
RNA anchoring preserves cell-type identifiability in spatial multi-omics deconvolution. **a**, Spatial heatmaps of dominated cell types in simulated multi-omics data with known ground truth. Compared with symmetric multi-modal approaches (NNLS, NMF and SpatialGlue-S), SMODER more accurately recovers spatial patterns and preserves distinct cell-type boundaries, closely matching the ground truth. **b**, Visualization of inferred composition for a representative cell type across spatial locations. SMODER reconstructs sharp and coherent spatial domains consistent with the ground truth. **c**, Quantitative evaluation across five complementary metrics, including Pearson correlation coefficient (PCC), Spearman correlation coefficient (SPCC), structural similarity (SSIM), root mean squared error (RMSE) and Jensen–Shannon divergence (JSD). SMODER consistently outperforms alternative approaches, achieving higher correlation and structural fidelity together with lower estimation error and distributional divergence. **d**, Robustness to reference misspecification under three scenarios: removal of a cell type, addition of an extraneous cell type and misannotation of cell identities. Radar plots summarize method rankings across evaluation metrics, with larger enclosed areas indicating better overall performance. SMODER maintains superior and stable performance across all perturbation settings, highlighting the advantage of RNA-anchored integration.

Compared with RNA-only methods, SMODER achieved consistently improved performance across multiple evaluation metrics (Fig. 2c and Supplementary Fig. S4a), including higher PCC, SPCC and SSIM, and lower RMSE and JSD. Quantitatively, it reduced RMSE by 32.62% and JSD by 45.46%, while increasing PCC by 19.08% and SSIM by 12.86% relative to the second-best method (RCTD).

Robustness analysis further showed that SMODER is insensitive to key hyperparameters, including the reduced feature dimension for each modality, the embedding dimension of graph encoders, the log_2_ fold-change threshold for selecting informative genes, and the kNN graph neighborhood size (Methods and Supplementary Notes). Stable performance was maintained across a wide range of parameter settings (Supplementary Fig. S4b).

We further evaluated the robustness of SMODER to misspecification of the scRNA-seq reference by systematically perturbing the reference annotation. Three scenarios were considered: removal of a cell type, mislabeling of cell identities, and introduction of an additional (spurious) cell type (Methods). Across these settings, SMODER was consistently compared against *M* competing methods using five evaluation metrics, with higher ranks indicating better performance. Across all perturbation regimes, SMODER consistently outperformed RNA-only baselines, demonstrating stable and accurate estimation of relative abundance under imperfect reference conditions (Fig. 2d and Supplementary Fig. S5). In the challenging setting of missing cell types, SMODER achieved substantial improvements over the second-best method, reducing RMSE by 37.85% and JSD by 44.25%, while increasing PCC and SSIM by 19.0% and 7.24%, respectively (Supplementary Fig. S6). These results highlight the robustness of RNA-anchored inference when reference annotations are incomplete or perturbed.

Collectively, these results demonstrate that spatial multi-omics deconvolution is inherently asymmetric with respect to molecular modalities. Transcriptomic information provides a stable anchor for cell identity, while additional modalities serve primarily as complementary constraints that improve inference robustness under noise and reference uncertainty.

### SMODER achieves robust and accurate deconvolution across spatial multi-omics technologies

We first evaluated SMODER across diverse spatial multi-omics technologies and tissue contexts, including spatial CUT&Tag–RNA-seq and spatial RNA–protein co-profiling (Methods), using both technology-aware simulations with known ground-truth composition and real datasets with high-confidence manual annotations (Supplementary Figs S7–S8). Across all benchmarked settings, SMODER consistently achieved superior deconvolution accuracy compared with competing approaches. In simulated RNA–histone and RNA–protein datasets, SMODER improved both point-wise accuracy (RMSE) and distributional fidelity (JSD), as well as structural and correlation consistency (SSIM, PCC/SPCC), relative to the best competing methods, including RCTD and CARD (Fig. 3a and Supplementary Fig. S9, Methods). In real RNA–histone and RNA–protein datasets, SMODER also improved most of local mixing, domain separation, spatial continuity, boundary sharpness, and global concordance with annotated spatial domains, relative to the best competing methods, including cell2location, IRIS and RCTD (Fig. 3b and Supplementary Fig. S10). An ablation study on these real datasets further demonstrated the advantages of SMODER’s loss design (Supplementary Figs. S11–S12).

**Figure 3:**
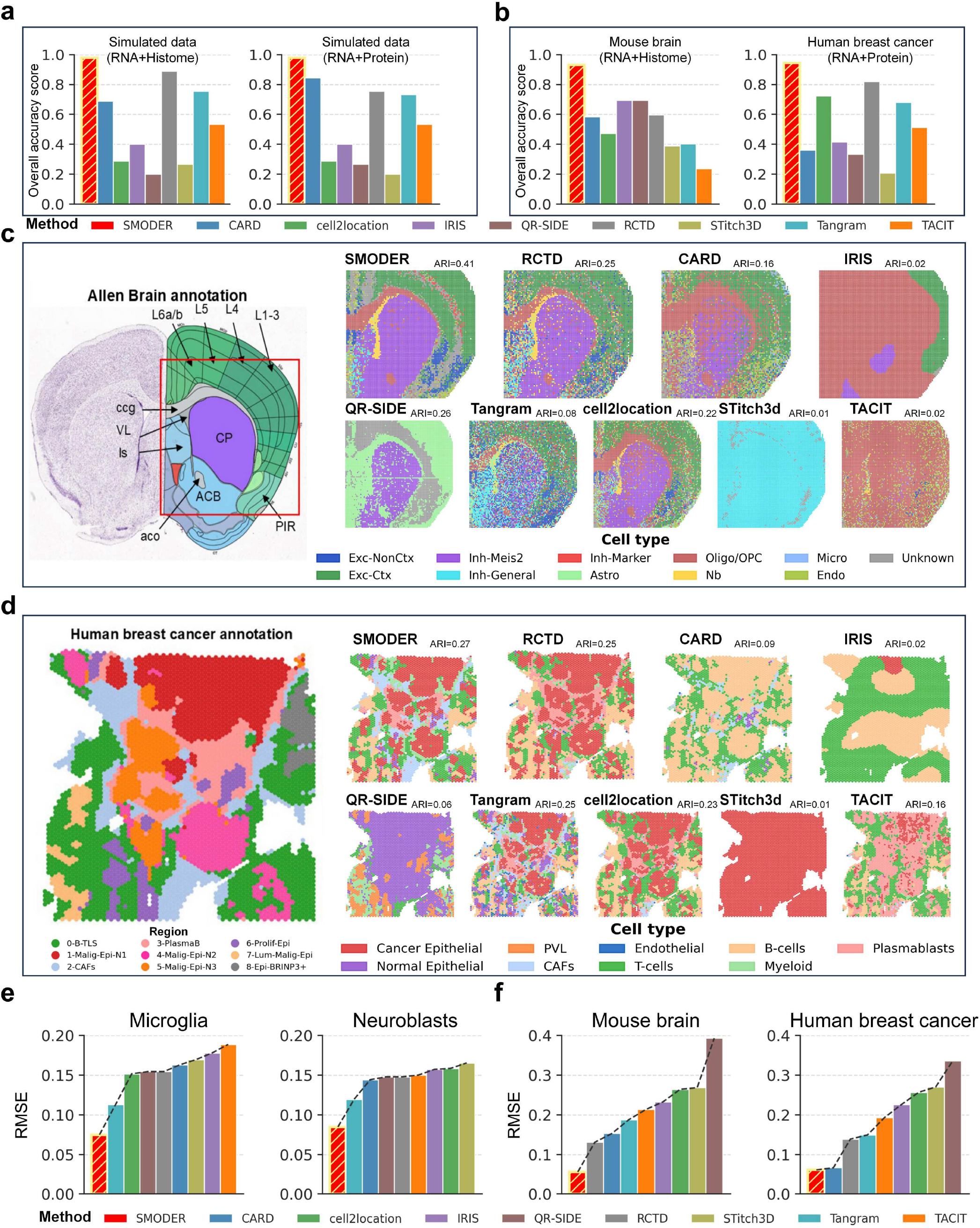
SMODER achieves robust and accurate deconvolution across spatial multiomics technologies. **a**, Overall deconvolution accuracy on technology-aware simulated spatial multi-omics datasets with known ground-truth relative abundance. Performance is summarized by an aggregated score across five complementary metrics, including RMSE, PCC, SPCC, JSD and SSIM. **b**, Performance comparison across real spatial multi-omics datasets with high-confidence manual annotations. SMODER achieves the highest overall accuracy aggregated across eight evaluation metrics, including LISI, SC, PAS, CHAOS, SMS, BEAR, ARI and NMI. **c**, Spatial coherence of inferred dominated cell-type distributions in mouse brain spatial CUT&Tag–RNA-seq data for the H3K27ac mark. SMODER reconstructs neuroanatomical structures including the genu of the corpus callosum (gcc), anterior commissure (aco), lateral ventricle (VL), caudoputamen (CP) and nucleus accumbens (ACB). **d**, Application to human breast cancer CytAssist RNA–protein co-profiling data. SMODER accurately localizes cancer epithelial, immune and CAFs populations to anatomically consistent regions, whereas existing methods fail to reliably identify cancer-enriched domains or misclassify key cell populations. **e**, Comparison of SMODER and RNA-only analyses. Incorporating complementary molecular layers improves discrimination of rare cell types (microglia, neuroblasts), while preserving RNA-driven cell-type resolution. **f**, Comparison of SMODER and RNA-only methods in deconvolution performance in regions with sparse RNA measurements.

Beyond global accuracy, SMODER produced spatially coherent cell-type composition maps that recapitulate known anatomical and pathological structures across tissues in terms of dominant cell types and their relative abundance. In mouse brain spatial CUT&Tag–RNA-seq data, SMODER reconstructed cell-type composition distributions that closely matched neuroanatomical organization, achieving higher agreement with annotated spatial domains than competing methods (ARI=0.41, Fig. 3c and Supplementary Fig. S13-S15). The inferred spatial patterns accurately resolved major brain regions, including cortical and subcortical structures, and exhibited sharper spatial boundaries with reduced cross-region mixing (Supplementary Fig. S16). Similar improvements were observed in human breast cancer RNA–protein datasets, where SMODER more faithfully recovered cancer epithelial, CAFs, immune and stromal compartments, yielding higher concordance with manual annotations (Fig. 3d and Supplementary Figs S17– S18). These results indicate that RNA-anchored inference not only improves quantitative accuracy but also enhances spatial structure recovery across heterogeneous tissue architectures.

Finally, we examined whether incorporating complementary molecular modalities improves estimation in challenging settings, including rare cell populations and spatially sparse regions. SMODER substantially improved the detection and quantification of rare cell types, such as microglia and neuroblasts, where RNA signals alone are often insufficiently informative (Fig. 3e and Supplementary Figs S19–S22). In addition, multimodal integration in SMODER reduced estimation error and stabilized cell-type composition estimation in regions with sparse transcriptomic coverage (Fig. 3f and Supplementary Fig. S23). Importantly, these gains were achieved without compromising the RNA-defined cell-type reference, indicating that additional modalities act as structured regularizers rather than competing signals.

Together, these results demonstrate that SMODER provides a unified framework for accurate, spatially coherent, and robust estimation of relative abundance across heterogeneous spatial multi-omics platforms by anchoring cell identity in transcriptomic space while leveraging complementary modalities to refine inference.

### Joint modeling enables cell-type-aware spatial epigenetic regulation

Spatial multi-omics provides an opportunity to directly connect transcriptional output with epigenetic regulation in situ. However, because each spatial measurement aggregates multiple cells, epigenomic signals are inherently confounded by cell-type composition, making it difficult to disentangle regulatory variation from mixture effects in conventional spot-level or symmetric integration analyses [23]. By anchoring cell identity in transcriptomic references while jointly modeling complementary epigenomic signals, SMODER enables decomposition of spatial multi-omics data into cell-type-resolved regulatory programs.

We first evaluated SMODER on a mouse brain spatial CUT&Tag–RNA-seq dataset profiling gene expression and H3K27me3 at 20-*µ*m resolution. SMODER accurately recovered spatially coherent cell-type composition distributions that recapitulated known neuroanatomical structures despite substantial cellular mixing and technical noise (Fig. 4a–b), while also revealing continuous transitions in transcriptional composition across anatomical boundaries (Fig. 4c and Supplementary Fig. S24). At the molecular level, SMODER reconstructed denoised spatial expression maps that recovered canonical lineage markers, including *Adora2a, Drd2, Calb1, Olig2*, and *Sox2*, across neuronal and glial populations [28–31], which are otherwise obscured in raw measurements (Supplementary Fig. S25).

**Figure 4:**
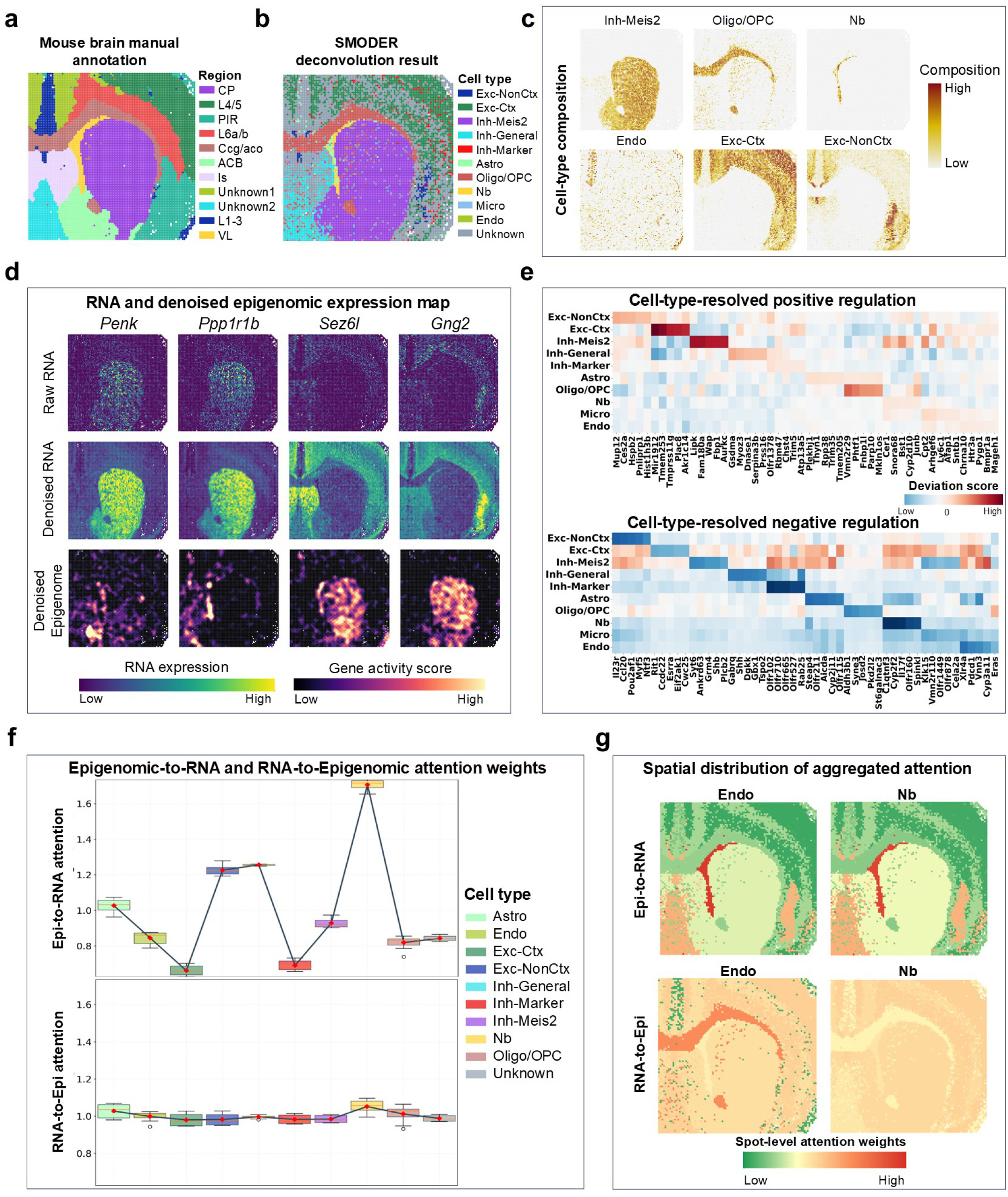
RNA-anchored joint modeling enables cell-type-aware spatial epigenomic analysis in mouse brain. Spatial multi-omics measurements aggregate multiple cell types within each spatial location, confounding regulatory variation with cell-type composition and obscuring cell-type-specific programs in spot-level analyses. By anchoring cell identity in transcriptomic references and decomposing spatial signals using inferred relative abundance, SMODER enables cell-type-resolved epigenomic analysis *in situ*. **a**, Manual annotation of neuroanatomical domains in a mouse brain spatial CUT&Tag–RNA-seq dataset profiling mRNA expression and the repressive histone modification H3K27me3 at 20-*µ*m resolution. Abbreviations: CP, caudoputamen; PIR, piriform cortex; L4/5, cortical layers 4/5; L6a/b, cortical layers 6a/6b; ccg/aco, genu of the corpus callosum and anterior commissure; ACB, nucleus accumbens; VL, lateral ventricle. **b**, Spatial heatmap of dominated cell types inferred by SMODER, showing strong agreement with known neuroanatomical structures. Cell-type abbreviations: Exc-NonCtx, non-cortical excitatory neurons; Exc-Ctx, cortical excitatory neurons; Inh-Meis2, Meis2-positive inhibitory neurons; Inh-General, general inhibitory neurons; Inh-Marker, marker-defined inhibitory neurons; Astro, astrocytes; Oligo/OPC, oligodendrocytes and oligodendrocyte precursor cells; Nb, neuroblasts; Micro, microglia; Endo, endothelial cells. **c**, Continuous cell-type proportion maps inferred by SMODER, capturing graded spatial transitions beyond discrete domain assignments for six representative cell populations, Inh-Meis2, Oligo/OPC, Nb, Endo, Exc-Ctx, and Exc-nonCtx. **d**, Cell-type-resolved relationships between H3K27me3 deposition and gene expression. Striatal inhibitory neuron markers *Penk* and *Ppp1r1b* exhibit localized expression within the caudoputamen (CP) and nucleus accumbens (ACB), accompanied by reduced H3K27me3 at the corresponding loci in inhibitory neuron populations (e.g., Inh Meis2). In contrast, surrounding regions show elevated H3K27me3 enrichment. Conversely, genes such as *Sez6l* and *Gng2* display high H3K27me3 levels and minimal transcriptional activity within inhibitory neuron populations while remaining expressed in adjacent regions dominated by other cell types. **e**, Heatmaps of cell types and the genes with regulation score. Upper panel: cell-type-specific regulation program with positive regulation scores; Lower panel: cell-type-specific regulation program with negative regulation scores. **f**, Boxplots of attention weights across cells in each dominated cell type learned by SMODER, where two directions’ weights both are visualized. Epigenomic signals exert strong and structured influence on transcriptional states, particularly in neuroblasts. **g**, Dominant cell type-specific attention weights between dual modalities for two representative cell types: Endo and Nb. Upper panel: spatial heatmaps show Epigenome-to-RNA spot-level attention scores, visualized separately for individual cell types. Lower panel: spatial heatmaps show RNA-to-Epigenome spot-level attention scores across different cell types, visualized separately for individual cell types.

At the gene level, we observed spatially coordinated coupling between transcription and H3K27me3 deposition consistent with known regulatory logic. For example, striatal inhibitory neuron markers (*Penk, Ppp1r1b*) showed localized activation in the caudoputamen and nucleus accumbens accompanied by reduced repressive chromatin in corresponding inhibitory populations (i.e., Inh-Meis2), whereas Polycomb-associated repression dominated in surrounding regions; conversely, genes such as *Sez6l* and *Gng2* exhibited inverted patterns across cell types (Fig. 4d). These observations provide direct validation of spatially resolved epigenetic–transcriptional coupling at gene resolution.

Beyond individual loci, SMODER reveals contrasting cell-type-resolved regulatory programs linking H3K27me3 activity with transcriptional output across tissue. Genes with high positive regulatory deviation scores (Methods) display cell-type-specific enrichment of H3K27me3 signals accompanied by reduced transcriptional activity, particularly within neuronal and oligodendrocyte populations (Fig. 4e, upper panel). In contrast, genes with strongly negative scores exhibit derepression-associated patterns marked by reduced H3K27me3 activity and elevated transcription in neurogenic and excitatory populations (Fig. 4e, lower panel). Importantly, most cell types participate in both regulatory regimes across distinct gene subsets, suggesting that epigenetic regulation is modular and context-dependent rather than fixed at the cell-type level.

To elucidate the mechanisms underlying these regulatory programs, we further examined the bidirectional attention weights learned by SMODER. After aggregating spot-level attention weights to the dominant cell-type level based on inferred dominant cell types, we observed a pronounced directional asymmetry: epigenomic signals exert a strong and structured influence on transcriptional states, particularly in neuroblasts, whereas the reverse direction shows substantially weaker cell-type specificity (Fig. 4f–g and Supplementary Fig. S26). This asymmetric interaction pattern provides a mechanistic explanation for the observed regulatory programs, suggesting that chromatin state acts as a primary latent driver of transcriptional variability in dynamic cell states.

We next asked whether this RNA-anchored regulatory principle generalizes to developmental chromatin regulation. Applying SMODER to a mouse embryo spatial RNA–ATAC-seq dataset at embryonic day 13, we recovered coherent tissue architectures across developing organs and observed strong concordance between RNA expression and ATAC-derived gene activity scores, indicating consistent spatial regulatory signals across modalities (Fig. 5a–b and Supplementary Figs S27–S28). Focusing on the neurodevelopmental regulator *Pax6* [32, 33], we observed that its spatial expression is decomposed into neural-dominant cell-type contributions (Fig. 5c). Furthermore, we found that distal regulatory elements linked to *Pax6* exhibit predominantly neuron-specific accessibility, with a subset showing significant cell-type-dependent peak–gene coupling (Fig. 5d), while spatial comparison of Pax6-high and Pax6-low regions further reveals compartmentalized enhancer activity defining transcriptionally active developmental domains (Fig. 5e). Consistent results were observed in the cell-type-specific decomposition of gene expression for additional neuronal marker genes, including *Snap25, Rbfox3*, and *Tubb3*, as well as in their associated distal regulatory elements and spatial expression patterns (Supplementary Figs S29—S31), further reinforcing this regulatory organization.

**Figure 5:**
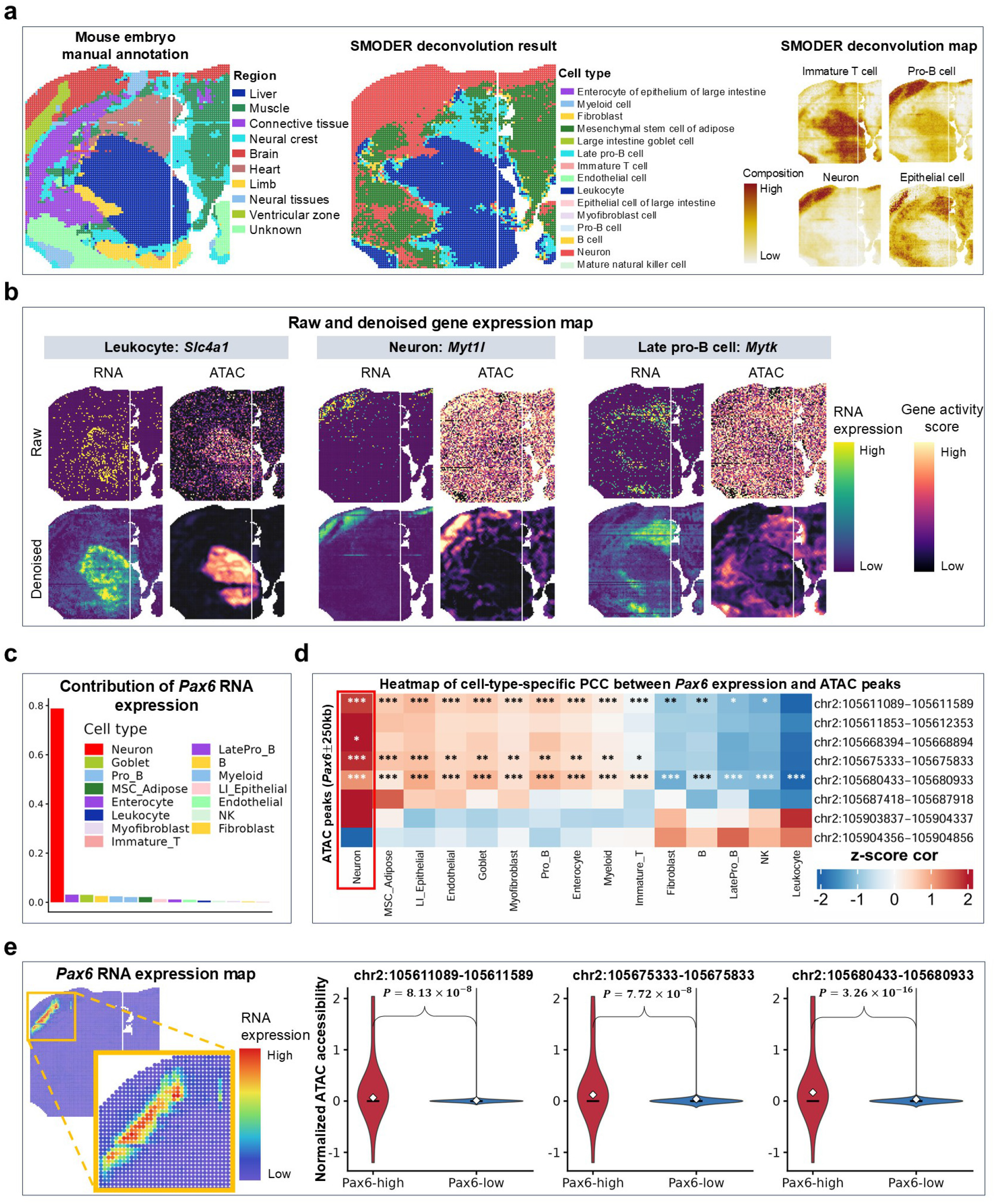
Cell-type-aware integration of spatial RNA–ATAC-seq data reveals regulatory mechanisms underlying E13 embryonic brain development. **a**, Left panel: spatial heatmap of manual annotated regions; middle panel: spatial heatmap of dominated cell types inferred by SMODER, illustrating the cellular architecture of developing organs, including brain, liver and heart; right panel: spatial heatmaps of proportions inferred by SMODER for four representative cell types. **b**, Denoised spatial maps of chromatin accessibility and RNA expression generated by SMODER, highlighting coherent spatial patterns that are obscured in the raw measurements. Representative marker genes for three cell types are shown: leukocytes (*Slc4a1* ), neurons (*Myt1l* ), and late pro-B cells (*Mytk* ). **c**, Decomposition of spatial *Pax6* expression into cell-type-specific contributions using SMODER-inferred relative abundance, showing dominant expression in neural populations. **d**, Proportion-based association analysis linking chromatin accessibility at distal regulatory elements to *Pax6* expression at cell-type resolution. Each row represents an ATAC peak within ±250kb of the *Pax6* transcription start site, and each column corresponds to a major cell type. Colors indicate z-scored Spearman correlation coefficients between peak accessibility and cell-type-specific *Pax6* expression (red, positive; blue, negative). Statistical significance is indicated by * (0.01 *< p* ≤ 0.05), ** (0.001 *< p* ≤ 0.01) or *** (*p* ≤ 0.001). **e**, Comparison of chromatin accessibility at *Pax6* - associated distal regulatory elements between *Pax6* -high and *Pax6* -low spatial regions, revealing selective enhancer accessibility in transcriptionally active domains.

Collectively, these results demonstrate a unified principle of RNA-anchored spatial epigenetic regulation: transcriptomic identity defines a stable reference axis, while epigenomic signals act as structured modulators of cell-type-specific regulatory programs. SMODER operationalizes this principle to enable mechanistic decomposition of spatial multi-omics data across both adult and developmental systems.

### SMODER generalizes across references, resolutions and noise regimes

To assess robustness and generalizability under realistic experimental variability, we evaluated SMODER across four settings capturing key sources of uncertainty in spatial multi-omics deconvolution, including measurement noise and reference misspecification, as well as changes in spatial resolution and tissue context.

We first assessed robustness to technical noise by introducing increasing levels of modality-specific dropout. SMODER consistently outperformed competing methods across all noise regimes (Fig. 6a–b and Supplementary Fig. S32), while preserving spatial coherence and accurate estimation of relative abundance for both dominant and rare cell types (Supplementary Fig. S33). Even under extreme dropout (80–90%), performance degraded smoothly rather than abruptly, indicating that multimodal integration stabilizes inference beyond RNA-only signals.

**Figure 6:**
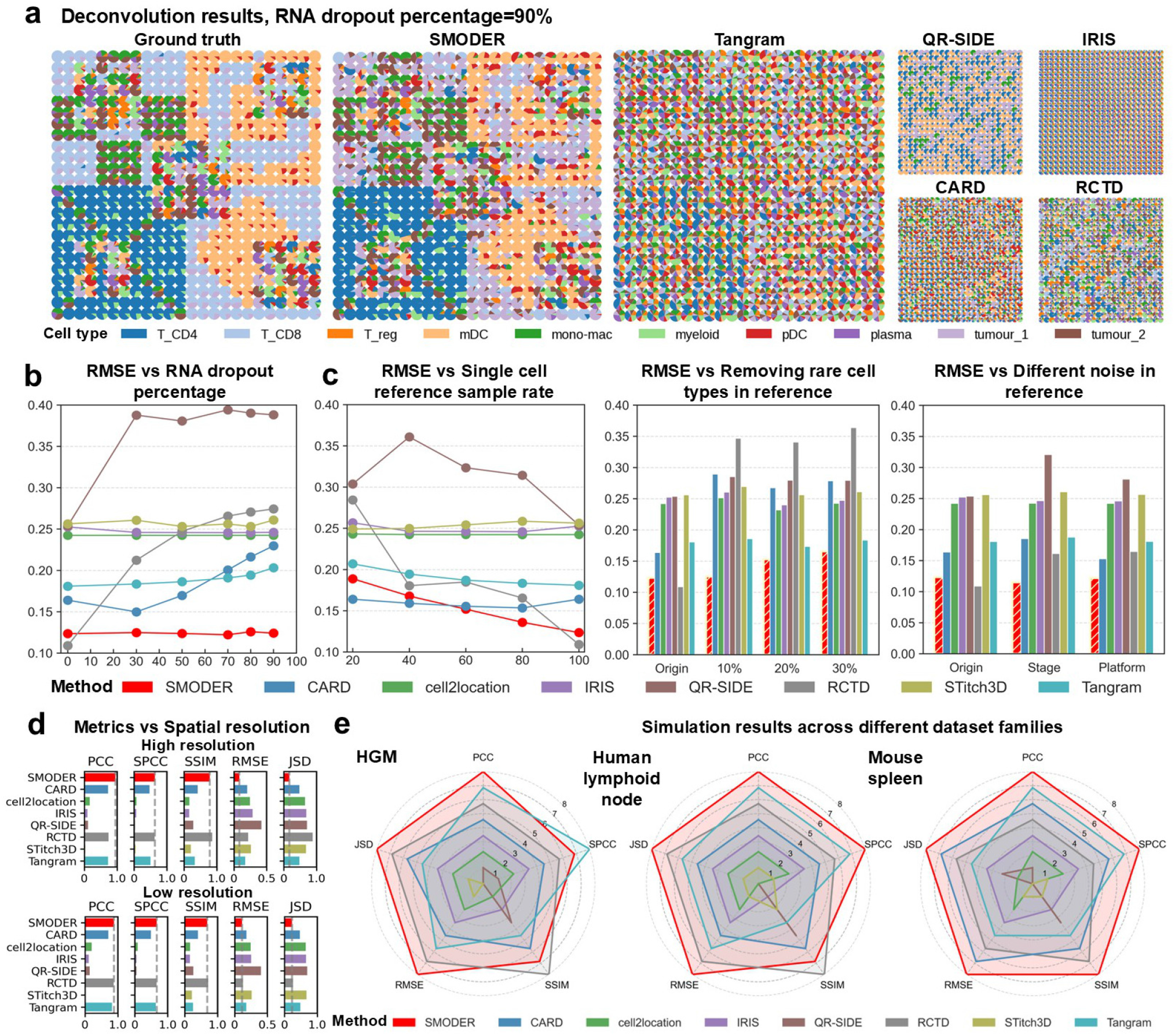
SMODER exhibits robust performance under noise, reference perturbation, spatial resolution changes, and diverse data conditions. **a**, Qualitative results on simulated spatial multi-omics data under extreme RNA dropout (90%). SMODER preserves spatial cell-type structure, whereas competing methods (Tangram, QR-SIDE, IRIS, CARD, RCTD) show degraded and noisy reconstructions. **b**, Robustness to RNA dropout. RMSE across increasing dropout levels, showing stable performance of SMODER under progressively sparser data. **c**, Sensitivity to scRNA-seq reference perturbation. Left: RMSE under varying reference sampling rates, assessing sensitivity to reference size. Middle: RMSE under progressive removal of rare cell types, evaluating robustness to incomplete references. Right: RMSE under reference noise, including stage-specific and platform-specific perturbations, measuring robustness to distributional shifts in the reference atlas. **d**, Performance across high-resolution and low-resolution settings, showing stable accuracy and spatial structure preservation of SMODER. **e**, Cross-dataset generalization. Radar plots summarizing performance across three simulated datasets constructed from different reference sources (HGM, lymph node, and spleen), demonstrating consistently strong performance of SMODER across diverse biological contexts.

We next evaluated sensitivity to reference misspecification by perturbing the scRNA-seq atlas, including downsampling, cell-type removal, and cross-platform or cross-stage substitution. Across all perturbations, SMODER showed only modest performance degradation relative to matched-reference settings (Fig. 6c and Supplementary Figs S34–S36), whereas RNA-only approaches were substantially more sensitive to reference quality. This highlights the benefit of RNA-anchored modeling with complementary modalities in maintaining stable inference under imperfect references.

We then examined robustness to spatial resolution by applying SMODER to datasets with different spot sizes. SMODER maintained accurate cell-type composition estimates and coherent spatial structure even under highly aggregated measurements, where each spot contains mixtures of many cell types (Fig. 6d). This robustness reflects the synergy between spatial neighborhood modeling and RNA-anchored integration, which together mitigate loss of transcriptomic resolution at lower spatial scales.

Finally, we evaluated cross-dataset generalization across tissues and species, including human lymphoid tissue, mouse spleen, and human glioblastoma profiled with distinct experimental platforms. Without dataset-specific tuning, SMODER consistently achieved strong performance across all settings (Fig. 6e and Supplementary Fig. S37), demonstrating broad applicability to heterogeneous spatial multi-omics data.

In summary, these results establish SMODER as a robust and generalizable framework for spatial multi-omics deconvolution, maintaining stable performance across diverse experimental conditions and data regimes.

## DISCUSSION

Spatial multi-omics technologies offer unprecedented opportunities to interrogate gene regulation and cellular organization directly within intact tissues. However, disentangling cell-type-specific signals from spatially aggregated measurements remains a fundamental challenge that limits mechanistic interpretation. In this study, we present SMODER, an RNA-anchored framework for spatial multi-omics deconvolution that enables accurate estimation of cell-type relative abundance while preserving and leveraging complementary regulatory information from additional molecular layers.

A central insight of this work is that spatial multi-omics integration is inherently asymmetric. Rather than treating modalities equally [24, 34], anchoring cell-type identity in transcriptomic information provides a stable reference axis that preserves interpretability, while additional modalities act as structured modulators of cellular states. Our results demonstrate that this RNA-anchored strategy consistently improves robustness and accuracy across diverse experimental conditions, suggesting that asymmetric integration is not merely a design choice but a principled solution for multi-modal spatial inference.

Beyond improving the estimation of relative abundance, this framework expands the scope of spatial multi-omics analysis from descriptive mapping to mechanistic inference. By jointly modeling transcriptomic and epigenomic layers, SMODER enables cell-type-resolved characterization of regulatory programs, linking chromatin state to transcriptional output within spatial context [35, 36]. This capability allows systematic dissection of regulatory architecture, such as repression–activation programs and enhancer–gene coupling, that are inaccessible to RNA-only analyses or post hoc integration approaches.

SMODER is designed to be broadly applicable and robust to practical challenges, including reference mismatch, variable spatial resolution and technical noise. Nevertheless, several limitations remain. As a reference-based approach, SMODER depends on the availability and completeness of scRNA-seq atlases, and performance may degrade when relevant cell types are missing. In addition, the resolution and interpretability of inferred regulatory interactions are ultimately constrained by the quality of epigenomic measurements.

In summary, SMODER establishes an RNA-anchored paradigm for spatial multi-omics analysis that reconciles accurate estimation of relative abundance with integrative regulatory modeling. By enabling cell-type-specific interrogation of gene regulation directly within tissue context, this framework provides a foundation for extracting mechanistic insight from increasingly complex spatial multi-omics data.

## Methods

### SMODER model

We describe the technical formulation of SMODER here, with additional implementation details provided in the Supplementary Notes. 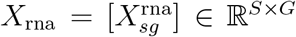 denote the RNA count matrix measured across *S* spatial locations for *G* informative genes, and let *X*_other_ ∈ ℝ^*S×M*^ denote the expression matrix for *M* features from a complementary modality (e.g., protein). A scRNA-seq reference defines *C* cell types and provides the cell-type-specific reference basis matrix *V* = [*V*_*cg*_] ∈ ℝ^*C×G*^, where *V*_*cg*_ represents the mean expression level of gene *g* in cell type *c*. SMODER consists of modality-specific graph encoders, a cross-modal attention fusion layer and a generative module that estimates relative abundance and reconstructs RNA counts.

#### Within-modality graph encoders

After dimensionality reduction for normalized data of RNA and complementary modalities, we obtain *F*_rna_ ∈ ℝ^*S×D*^ and *F*_other_ ∈ ℝ^*S×D*^, respectively. For each modality, two graph convolutional encoders are constructed based on feature similarity and spatial adjacency, respectively. Let 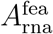 and 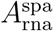 denote the adjacency matrices of the feature and spatial graphs. Graph convolutional encoding yields

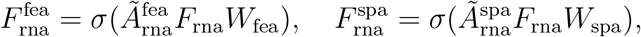

where *Ã* is the normalized adjacency matrix, *W* are learnable parameters and *σ*(·) denotes a nonlinear activation.

To combine feature-derived and spatial representations, we introduce an attention mechanism

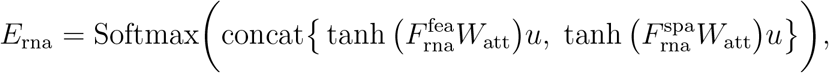

where *u* ∈ ℝ^*D*^ is a learnable attention vector and the Softmax function is applied row-wise, yielding *E*_rna_ ∈ ℝ^*S×*2^. The modality embedding is then

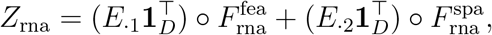

where *E*_.*j*_ denotes the *j*-th column of *E*_rna_, **1**_*D*_ is a *D*-dimensional vector of ones, and ○ denotes element-wise multiplication. The embedding *Z*_other_ is obtained analogously.

#### Cross-modal bidirectional attention

To integrate multimodal information, we employ a bidirectional cross-attention mechanism [37]. Through bidirectional interactions, each modality attends to informative signals from the other, enabling complementary information sharing. Given embeddings *Z*_rna_ and *Z*_other_, attention weights are computed as 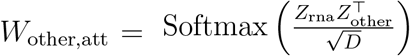, 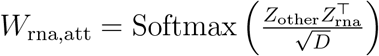. The resulting embeddings are

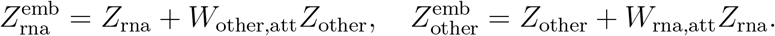

The concatenated representation 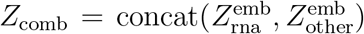 is passed through a fully connected layer

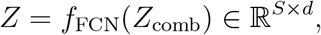

which serves as the final latent embedding.

#### Cell-type composition estimation

Based on the shared embedding *Z*, SMODER estimates relative abundance of each cell type in every spatial spot. Let *P*_*s,c*_ denote the relative abundance of cell type *c* in spot *s*, and define *P*_*s*_ = (*P*_*s*,1_, …, *P*_*s,C*_ )^*⊤*^ ∈ ℝ^*C*^, where *P*_*s,c*_ ≥ 0 and 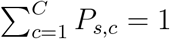. Let *Z*_*s*_ ∈ ℝ^*d*^ denote the latent embedding for spot *s*. We model the relative abundance as

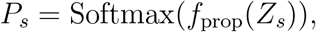

where *f*_prop_(·) is a two-layer full connected neural network. Since *Z*_*s*_ integrates RNA, complementary modality, and spatial information, the estimated relative abundance inherit cross-modal and spatial context. Critically, the biological interpretation of *P*_*s*_ as cell-type relative abundance is not imposed by the Softmax transformation alone, but is grounded in the negative binomial generative model (described below).

#### Negative binomial generative model

Given the reference basis matrix *V* and the relative abundance matrix *P*, SMODER reconstructs the observed RNA counts under a negative binomial (NB) generative model:

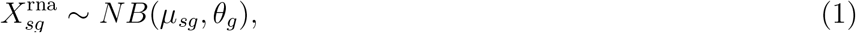

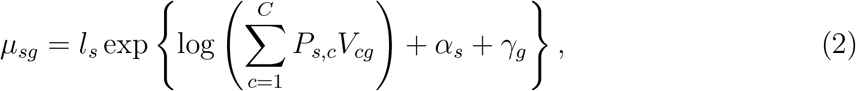

where *θ*_*g*_ denotes a gene-specified dispersion parameter, *l*_*s*_ denotes the total transcript count in spot *s*, and *α*_*s*_ and *γ*_*g*_ represent spot-specific and gene-specific effects, respectively, learned during training.

### Model training

To incorporate multiple aspects of the model, we construct a composite objective function. Specifically, four loss components are used to train the deconvolution network: the negative binomial log-likelihood loss, the reconstruction loss, the cross-modal consistency loss, and the spatial smoothing loss.

The negative binomial loss models RNA counts under the generative framework described in Eqns (1) and (2), ensuring accurate estimation of relative abundance while accounting for spot- and gene-specific effects. It is defined as

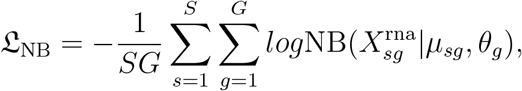

where 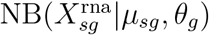 is the probability mass function of 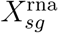, with specific form in Supplementary Notes.

The reconstruction loss preserves modality-specific structural information by constraining the latent embeddings to reconstruct the original low-dimensional features. To this end, two modality-specific decoders are introduced: 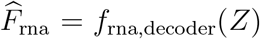 and 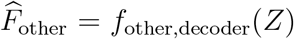. The mean squared error is used to measure the discrepancy between reconstructed and original features:

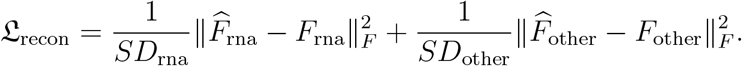

The cross-modal consistency loss enforces semantic alignment between modalities, promoting a unified cross-modal latent space. Specifically, the embedding of one modality is first decoded using the decoder of the other modality and then re-encoded by its encoder: 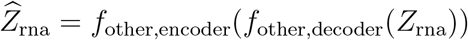 and 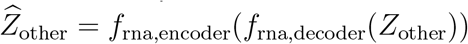. The resulting encoded output is then compared with the input to calculate discrepancies, given by

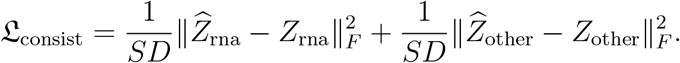

The spatial smoothing loss regularizes relative abundance across neighboring spots, enhancing spatial coherence and robustness. Based on the adjacency matrix *A*_*spa*_, we calculate the normalized Laplacian matrix 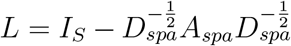, where *I*_S_ is an *S* × *S* identity matrix and *D*_*spa*_ is the diagonal degree matrix. We define the spatial smoothing loss as

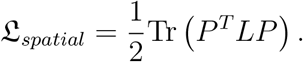

Building upon the four loss functions, SMODER incorporates adjustable parameters to formulate its loss function.

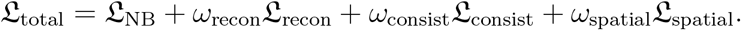

This framework trains parameters across the entire deconvolution network and attention layer.

### Implementation details

#### Preprocessing

Low-quality cells and genes were filtered from scRNA-seq data using Scanpy. Genes shared between scRNA-seq and spatial transcriptomics datasets were retained. Differentially expressed genes were selected either by choosing a fixed number per cell type or by applying a log_2_ fold-change threshold. The average expression of each gene within each cell type was computed to form the reference matrix *V* . For additional modalities, protein measurements were normalized using centered log-ratio transformation, whereas peak count matrices were normalized using TF–IDF. A detailed description of preprocessing is provided in the Supplementary Notes.

#### Dimensionality reduction

Principal component analysis (PCA) was applied to log-normalized RNA expression to obtain a 256-dimensional representation *F*_rna_. For the complementary modality (e.g., protein or epigenomic data), the resulting feature matrix is uniformly denoted by *F*_other_. Specifically, for protein data, PCA was applied when the number of proteins exceeded 256; otherwise, the centered log-ratio (CLR) normalized matrix was used directly. For epigenomic peak matrices, latent semantic indexing (LSI) was used to derive a 256-dimensional representation.

#### Graph construction

Feature graphs were constructed for each modality using the dimensionally reduced feature matrices. A spatial graph was defined using spot coordinates. All graphs were generated using a *k*-nearest-neighbor strategy with *k* = 8 neighbors per node.

### Downstream analyses

#### Denoising of multi-omic expression maps

Due to the substantially higher sparsity of epigenomic data compared with RNA and protein modalities, we adopt modality-specific strategies for denoising.

#### Denoising RNA/protein expression maps

To capture high-frequency spatial variation in expression, we augment spatial coordinates using Fourier feature mappings. Given spatial coordinates 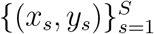, we construct transformed features as

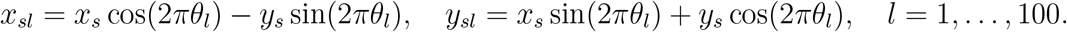

These Fourier features are concatenated with the normalized modality-specific encoder output to form the input feature matrix 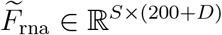. The normalized RNA or protein expression matrix is used as the reconstruction target. A three-layer fully connected network is trained using the Adam optimizer to reconstruct the spatial expression profiles.

#### Denoising ATAC/Histone mark expression maps

Epigenomic gene activity scores exhibit sparse and locally clustered patterns. To account for this structure, we employ a spatial graph convolutional network (GCN) for denoising. Raw gene activity scores are first filtered and log-normalized, and then used as the reconstruction target.

The modality-specific embedding *Z*_other_ and the spatial adjacency matrix 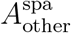 are used as inputs to the GCN. The network consists of two hidden layers, each with 256 dimensions, and is trained using the Adam optimizer with mean squared error loss. To mitigate overfitting due to sparsity, dropout (rate 0.2) and *ℓ*_2_ weight decay are applied. The reconstructed epigenomic expression profiles are obtained as the model output.

#### Cell-type-specific regulation deviation score

Let 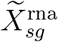 and 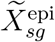 denote log-normalized spatial transcriptomic and epigenomic signals with matched genes/loci, and *P* ∈ ℝ^*S×C*^ the cell-type relative abundance matrix. Cell-type-specific signals were aggregated as: 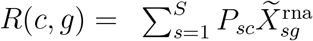 and 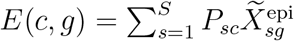. We standardized signals across cell types: 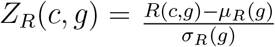 and 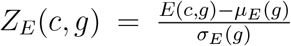, where *µ*_*·*_(*g*) and *σ*_*·*_(*g*) denote mean and standard deviation across cell types. The cross-modality difference is defined as:

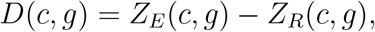

which quantifies the relative imbalance between H3K27me3-associated chromatin repression and transcriptional activity for gene *g* in cell type *c*. We further define the cell-type-specific regulation deviation score:

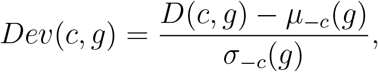

where *µ*_*™c*_(*g*) and *σ*_*™c*_(*g*) are computed across all cell types except *c*. A high positive deviation score indicates relatively stronger epigenomic repression (or accessibility imbalance) coupled with reduced transcriptional output in a specific cell type compared with other cell types, whereas negative values indicate relatively enhanced transcriptional activity.

#### Identification of cell-type-specific regulatory elements

To identify cell-type-specific cisregulatory elements, we examined composition-weighted correlations between gene expression and chromatin accessibility across spatial locations. Recall that 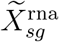 denote the normalized expression of gene *g* at spatial spot *s, P*_*sc*_ the inferred composition of cell type *c* at that spot, and 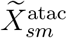 the accessibility of ATAC peak *m*. To approximate cell-type-specific expression, we constructed a contribution-weighted expression 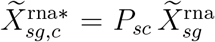. For each peak *m* and cell type *c*, regulatory association was quantified by the Pearson correlation

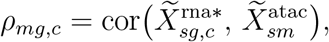

computed across spatial spots. Corresponding *P* values were obtained using a two-sided correlation test.

#### Decomposition of cell-type-specific expression contribution

To quantify cell-type-specific contributions to spatial gene expression, we modeled the observed expression at each spot as a linear combination of cell-type relative abundance and cell-type-specific expression profiles. For gene *g*, the expression attributable to cell type *c* at spot *s* was defined as 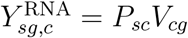, and the overall contribution of cell type *c* to gene *g* across all spots was calculated as

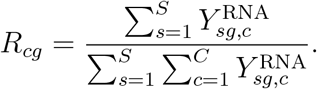

This decomposition provides a concise mapping from cell-type composition to spatial gene expression, enabling the identification of cell-type-dominated expression patterns across the tissue.

### Simulation studies

#### Simulation framework

To systematically evaluate deconvolution performance under controlled conditions, we generated simulated spatial multi-omics datasets by aggregating single cells from matched single-cell multi-omic references. Specifically, single-cell datasets with paired RNA and secondary modalities (e.g., chromatin accessibility or protein abundance) were used as the ground truth. Spatial spots were constructed by randomly sampling cells according to predefined cell-type compositions and aggregating their molecular profiles, thereby mimicking the mixture structure commonly observed in spatial multi-omics experiments.

Unless otherwise specified, simulated datasets were generated on a 24 × 24 spatial grid with an average of ten cells per spatial location. relative abundance were assigned according to spatial regions to reproduce structured tissue organization, and molecular counts were aggregated across sampled cells to form spatial multi-omic measurements. Modality-specific noise and sparsity were subsequently introduced to emulate realistic experimental characteristics. Each simulation experiment was repeated five times to ensure robust evaluation. Detailed descriptions of all simulation settings are provided in the Supplementary Notes.

#### Concept validation simulation

To validate the proposed RNA-anchored modeling strategy, we designed simulations that explicitly contrast it with symmetric multi-modal deconvolution approaches under controlled conditions. Specifically, we generated spatial multiomics data where RNA and the secondary modality (ATAC or protein) exhibit different levels of noise, sparsity, and cell-type specificity. Cell-type compositions were first sampled according to predefined spatial domains, and modality-specific measurements were then generated from the same underlying cell mixtures. For RNA, counts were simulated to preserve strong cell-type signals, whereas for the secondary modality, additional noise and sparsity were introduced to reflect realistic measurement characteristics.

Under this setting, we compared SMODER with symmetric multi-modal baselines, including joint-feature NNLS and joint-latent NMF, all applied to the same simulated datasets with identical ground-truth relative abundance. This design enables a direct assessment of how different modeling strategies, anchoring cell identity in RNA versus treating modalities symmetrically, affect deconvolution accuracy. These simulations are intended as a proof-of-concept to demonstrate the impact of modality-specific modeling assumptions, rather than as a comprehensive benchmark of multi-modal deconvolution methods.

#### Reference-matched and perturbation experiments

In this experiment, we first generated reference-matched simulated data. We then evaluated robustness to imperfect single-cell references by introducing three types of perturbations: (i) removal of a cell type from the reference, (ii) misannotation of cell identities, and (iii) addition of an extraneous cell type. These scenarios mimic common challenges encountered in practical applications, including incomplete cell atlases, annotation errors, and the presence of unrelated cell populations.

#### Technology-aware simulations

To assess performance across spatial multi-omics technologies, simulations were constructed using real single-cell datasets representing distinct modality combinations, including RNA–epigenomic and RNA–protein measurements. These datasets provided realistic molecular distributions and cell-type heterogeneity. Simulated spatial profiles were generated by aggregating sampled cells while preserving modality-specific signal structures.

#### Robustness experiments

To evaluate the generalizability of SMODER under challenging conditions, additional simulations were performed by varying dropout rates, reference datasets, spatial resolutions and biological contexts. These experiments assess the stability of the method under increased measurement sparsity, reference mismatch and different cell densities across spatial locations.

### Benchmark methods

#### Symmetric multi-modal deconvolution baselines

To assess the effect of treating molecular modalities symmetrically during spatial deconvolution, we implemented two types of representative symmetric baselines.

The first baseline performs joint-feature deconvolution by enforcing a single set of relative abundance to simultaneously explain spatial RNA measurements and a secondary molecular modality. Specifically, RNA features and non-transcriptomic features are concatenated after appropriate scaling, and relative abundance are estimated by non-negative least square (NNLS) regression against concatenated reference signatures (Supplementary Algorithm 1). This approach treats RNA and the secondary modality on equal footing during inference.

The second baseline comprises two methods, both of which follow a joint-latent strategy commonly used in multimodal integration. In the first method, spatial RNA and secondary-modality measurements are jointly factorized using nonnegative matrix factorization (NMF) to learn a shared latent representation. After normalization, the resulting latent components are interpreted post hoc as cell-type mixture proportions (Supplementary Algorithm 2). In the second method, SpatialGlue [23] is used to learn a shared latent representation. A softmax transformation is then applied to convert the latent components into probabilities, which are subsequently interpreted as cell-type proportions. In both methods, the latent dimensions are not explicitly linked to biological cell types during model fitting.

Both baselines are symmetric with respect to molecular modalities. However, the first baseline represents an idealized setting, as paired reference data are often difficult to obtain in practice. In contrast, the joint-latent approach remains agnostic to cell-type identity during model estimation and therefore requires post hoc alignment of latent components to cell types.

#### Spatial RNA-only baseline methods

SMODER was compared with seven established spatial deconvolution methods: RCTD [7], CARD [8], IRIS [9], QR-SIDE [10], Tangram [11],cell2location [12], STitch3D [13] and TACIT-based baseline [38], with detailed descriptions provided in the Supplementary Notes.

### Evaluation metrics

#### Simulation metrics

In simulation studies, where the ground-truth cell-type proportion matrix is available, we evaluated deconvolution accuracy using five complementary metrics that capture pointwise error, linear correlation, rank concordance, distributional similarity, and structural fidelity. Root mean square error (RMSE) measures the average magnitude of estimation errors in the inferred relative abundance, with smaller values indicating higher accuracy. Pearson correlation coefficient (PCC) quantifies the linear association between estimated and true proportions. Spearman correlation coefficient (SPCC) assesses the monotonic agreement between estimated and true proportions, reflecting robustness to scale differences. To quantify similarity in the overall distribution of relative abundance, we employed Jensen– Shannon divergence (JSD), a symmetric and bounded measure of divergence between the estimated and true proportion distributions [39]. Lower JSD values indicate closer agreement. In addition, we used the structural similarity index (SSIM) to evaluate the preservation of spatial structure in the proportion maps by jointly comparing local means, variances, and spatial correlations [40]. Higher SSIM values indicate better recovery of spatial patterns present in the ground-truth proportions.

To facilitate comparison across a large number of metrics, we further summarized method performance using an overall accuracy score (OAS) based on rank aggregation [13]. Specifically, for each metric, the *M* competing methods were ranked from 1 to *M*, with larger rank values indicating better performance. Five sets of ranks were obtained corresponding to RMSE, PCC, SPCC, JSD, and SSIM. The overall accuracy score (OAS) was computed as

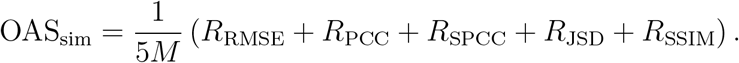

Here, OAS ranges from 1*/M* to 1, with larger values indicating better overall performance across metrics.

#### Real data metrics

In real data applications, where ground-truth relative abundance are unavailable, we assessed deconvolution performance using eight complementary metrics that characterize local mixing, domain separation, spatial continuity, boundary sharpness, and global concordance with annotated spatial domains. Local inverse Simpson’s index (LISI) [41] and silhouette coefficient (SC) evaluate local domain purity and separation in the inferred proportion space. Percentage of abnormal spots (PAS) and CHAOS score [42] quantify spatial incoherence by measuring neighborhood-level disagreement and within-domain spatial fragmentation, respectively. We define spatial map similarity (SMS) and balanced edge agreement ratio (BEAR) to directly compare inferred spatial patterns with annotated domains, assessing large-scale spatial alignment and local boundary consistency while correcting for spatial and class imbalance (Supplementary Notes). In addition, adjusted Rand index (ARI) and normalized mutual information (NMI) computed between dominated cell-type assignments and manual annotations assess the global consistency of inferred spatial structure.

The detailed definitions of all evaluation metrics are provided in Supplementary Notes. To comprehensively assess the performance, eight sets of ranks were obtained corresponding to LISI, SC, PAS, CHAOS, SMS, BEAR, ARI, and NMI, and the overall accuracy score was computed as

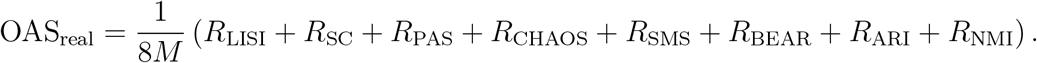

### Analyzed datasets

We evaluated our method on four publicly available spatial multi-omics datasets.

#### Mouse brain spatial CUT&Tag–RNA-seq datasets

We analyzed two mouse brain spatial CUT&Tag–RNA-seq datasets: one profiling RNA and H3K27ac (MB-ac) and the other profiling RNA and H3K27me3 (MB-me3). MB-ac contains 23,442 genes and 79,159 peaks across 9,363 spatial locations, whereas MB-me3 contains 25,740 genes and 59,331 peaks across 9,692 spatial locations. After preprocessing (see Supplementary Notes), MB-ac retained 8,928 genes and 20,000 peaks across 9,363 locations, and MB-me3 retained 9,331 genes and 20,000 peaks across 9,692 locations. Spatial domains annotated based on the Allen Brain Atlas and SpatialGlue were used as ground truth for deconvolution evaluation. The mouse brain snRNA-seq reference [12] comprises 40,532 cells and 31,017 genes across 11 major cell types (Supplementary Notes).

#### Human breast cancer FFPE CytAssist RNA–protein data

The human breast cancer formalin-fixed paraffin-embedded (FFPE) CytAssist dataset comprises paired spatial transcriptomic and protein measurements profiled using the 10x Genomics CytAssist platform. The raw dataset contains 17,957 genes and 35 antibody-derived protein markers measured across 4,169 spatial capture locations. After data preprocessing (Supplementary Notes), 1,774 genes and 35 proteins across 4,168 spatial locations were retained for downstream analysis. The corresponding human breast cancer scRNA-seq reference dataset [43] comprises 100,064 cells and 29,733 genes across 9 annotated cell types (Supplementary Notes).

#### Mouse embryo spatial RNA–ATAC data

The mouse embryo spatial RNA–ATAC dataset profiles paired gene expression and chromatin accessibility measurements across 8,886 spatial capture locations. The raw data include 21,380 genes and 30,566 accessible chromatin peaks representing the two molecular modalities. Following quality control and preprocessing (Supplementary Notes), including filtering of low-quality spatial spots and low-confidence features, 7,910 genes and 20,000 chromatin peaks were retained for downstream analysis. Peak selection was restricted to high-confidence accessible regions, while gene features were harmonized across modalities to ensure compatibility with the downstream integrative modeling framework. The corresponding mouse embryonic single-cell RNA-seq reference dataset [44] consists of 2,688 cells profiled at embryonic day 13 (E13), spanning 35,100 genes and 15 annotated cell types (Supplementary Notes).

## Acknowledgment

This work was supported by National Natural Science Foundation of China (12401361), Natural Science Foundation of Sichuan Province (2025ZNSFSC0809), Sichuan Provincial Natural Science Foundation Project (Innovative Research Group, 2026NSFSCZY0056), and the Fundamental Research Funds for the Central Universities of China.

## Author contributions

W.L. contributed to Conceptualization and Methodology. W.L. and Q.D. contributed to Project Administration. Y.B. contributed to Methodology and Software by developing the model and implementing the computational framework. S.X. and X.G. contributed to Software development of the associated toolkit. Y.B. and X.B.G. contributed to Investigation by conducting simulation studies and benchmark evaluations. W.L. drafted the initial manuscript (Writing – Original Draft), while Y.B., X.B.G., S.X., X.G., X.W., W.J., S.Y., Q.D., and W.L. collectively revised and edited the manuscript (Writing – Review & Editing).

## Data availability

All datasets used in this study are publicly available from the corresponding repositories listed below.

### Spatial multi-omics datasets

We analyzed two mouse brain spatial CUT&Tag–RNA-seq datasets (GSE205055), a mouse embryo spatial RNA–ATAC dataset (GSM6204637; GSM6204624), and a human breast cancer CytAssist RNA–protein dataset (10x Genomics: Gene and Protein Expression Library of Human Breast Cancer, FFPE, CytAssist platform).

### Single-cell and single-nucleus reference datasets

Reference datasets include a human melanoma joint RNA–ATAC single-cell dataset (SCP2171), a human central nervous system innate immune RNA–protein dataset (GSE245311), a mouse brain snRNA-seq dataset (E-MTAB-11115), a human breast cancer scRNA-seq dataset (GSE176078), a mouse embryo scRNA-seq dataset at embryonic day 13 (E-ENAD-15), a human lymphoid scRNA-seq dataset (Cell2location dataset portal), a mouse spleen scRNA-seq dataset (SRA653146), and a human glioblastoma scRNA-seq dataset (GSE182109).

### Anatomical reference

The anatomical reference of the P56 mouse brain was obtained from the Allen Brain Atlas (https://atlas.brain-map.org/).

## Code availability

We provide an open-source Python implementation of the SMODER framework at https://github.com/HokazeJunko/SMODER. Comprehensive documentation and tutorials are available at https://smoder.readthedocs.io/en/latest/. All analysis pipelines and Python scripts used to reproduce the results in this study, are publicly available at https://github.com/HokazeJunko/SMODER/tree/main/analysis_scripts.

