## Supplementary Material for "Deciphering Spatial Cellular Organization through RNA-Anchored Multi-Omics Deconvolution"

### Supplementary information for “Deciphering spatial cellular organization through RNA-anchored multi-omics deconvolution”

**6 Details of sensitivity analysis and ablation study** **51**

### Supplementary Figures

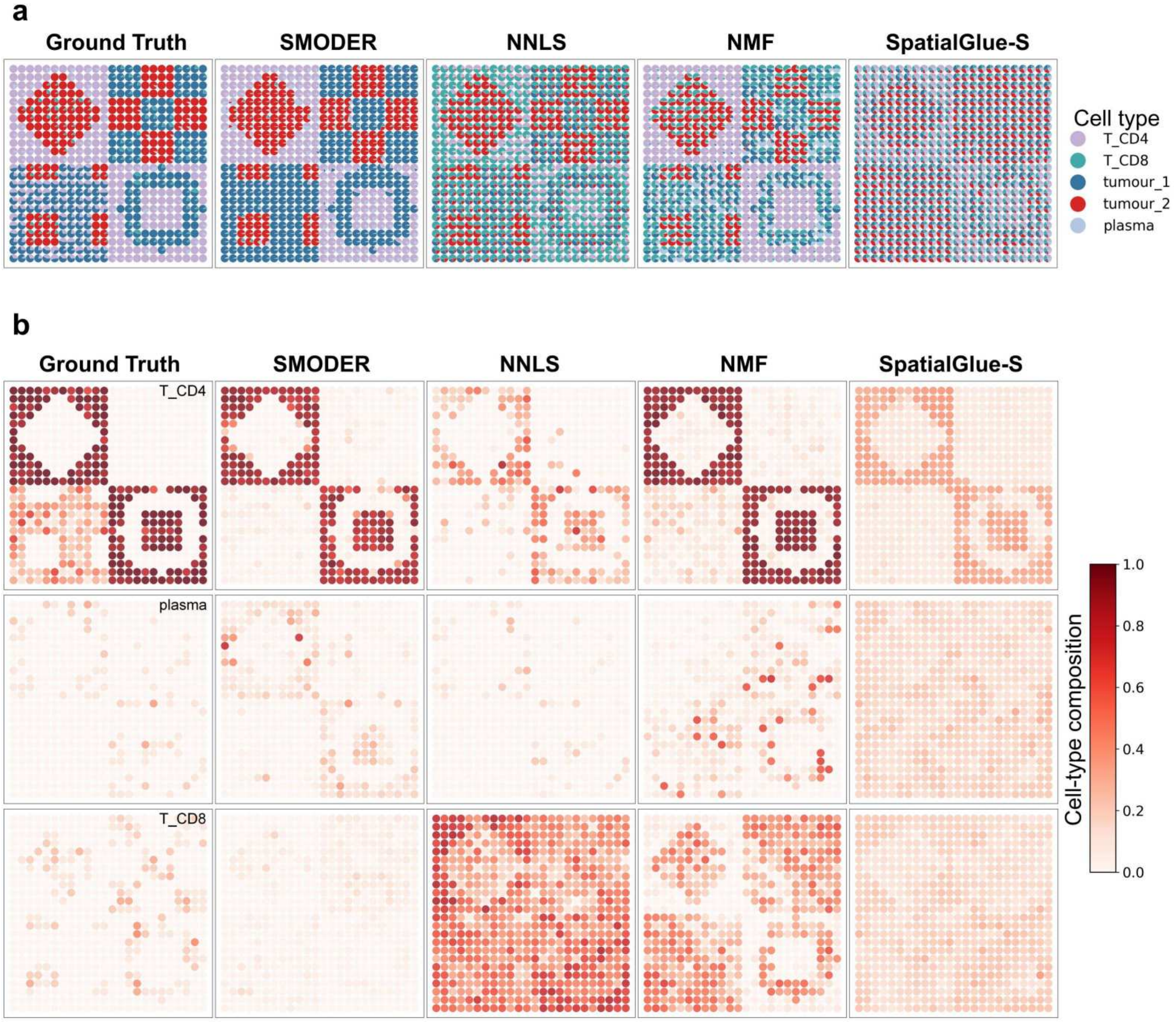

Supplementary Figure S1: **Proof-of-concept evaluation of SMODER for spatial multi-omic deconvolution in simulated human melanoma data with joint RNA and ATAC-seq profiles.** **a**, Comparison of cell-type composition estimates across spatial locations between ground truth and methods including SMODER, NNLS, NMF and SpatialGlue-S. **b**, Spatial heatmaps of inferred cell-type composition for three representative cell types (T_CD4, plasma, and T_CD8) compared with ground truth.

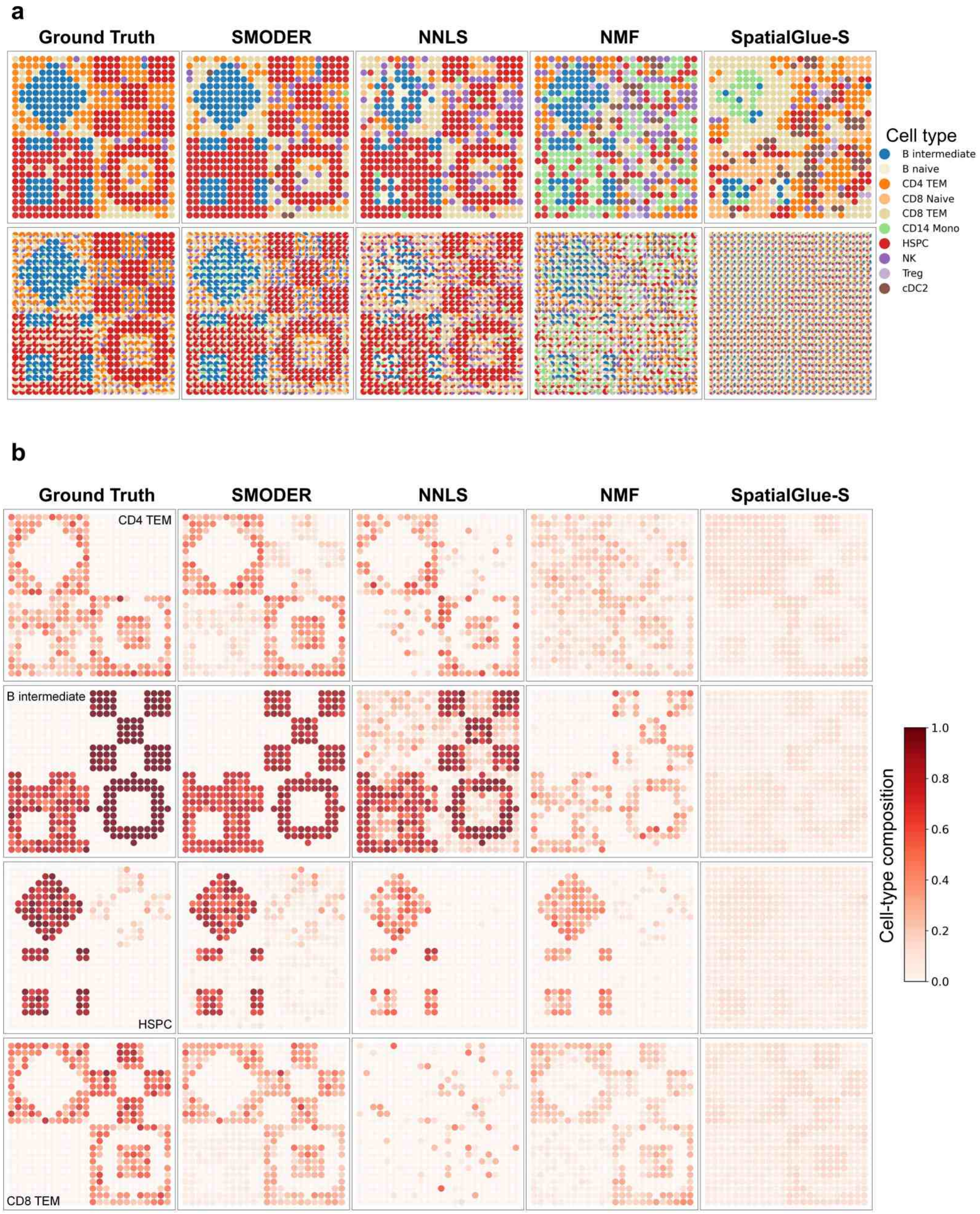

Supplementary Figure S2: **Proof-of-concept evaluation of SMODER for spatial multi-omic deconvolution in simulated human central nervous system (CNS) innate immune datasets with paired RNA and protein measurements.** **a**, Comparison of cell-type composition estimates across spatial locations between ground truth and methods including SMODER, NNLS, NMF and SpatialGlue-S. Upper panel: Dominated cell types; bottom panel: cell type composition. **b**, Spatial heatmaps of inferred cell-type composition for four representative cell types compared with ground truth.

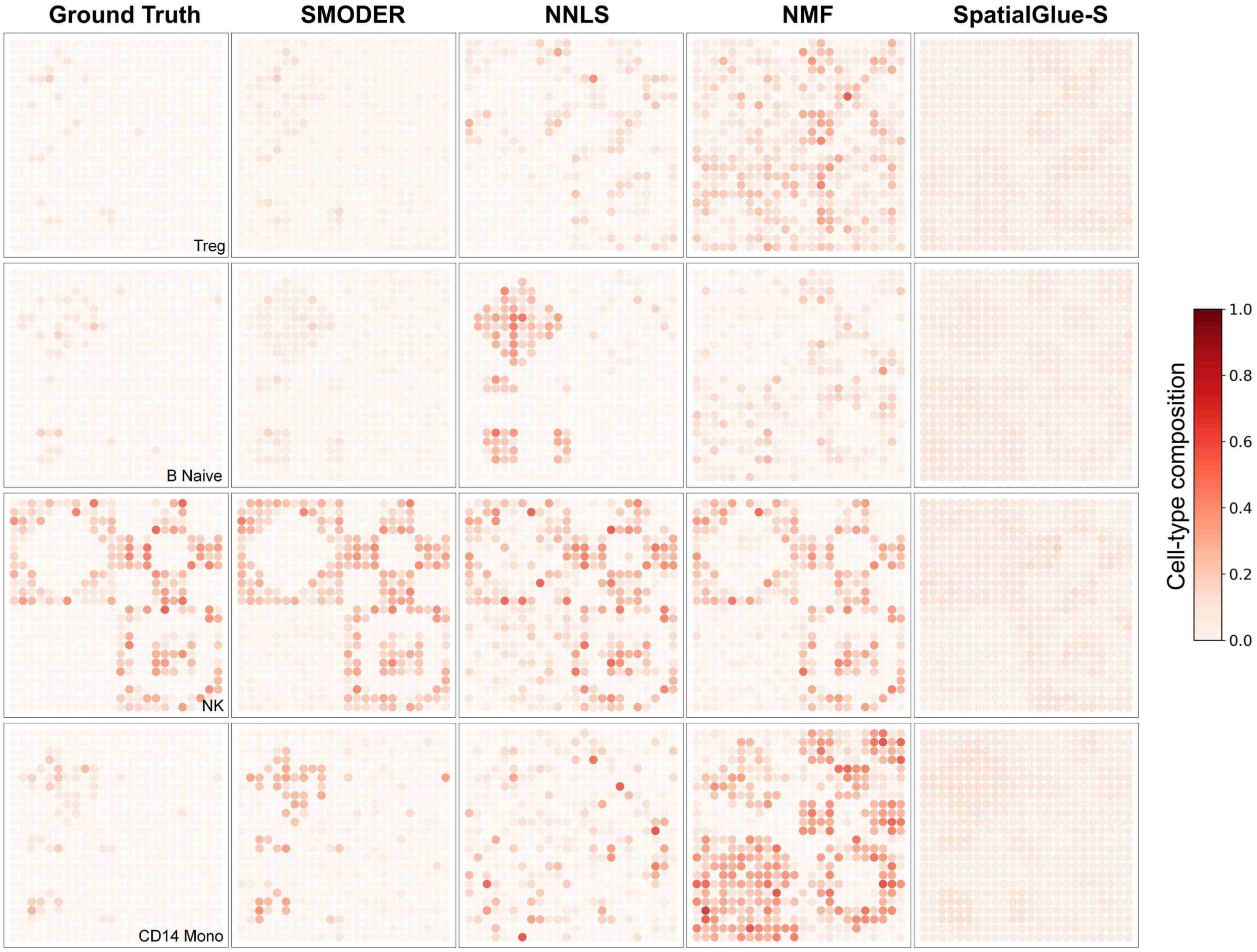

Supplementary Figure S3: **Proof-of-concept evaluation of SMODER for spatial multi-omic deconvolution in simulated human central nervous system (CNS) innate immune datasets with paired RNA and protein measurements.** Spatial heatmaps of cell-type composition for four cell types inferred by SMODER, NNLS, NMF and SpatialGlue-S, compared with ground truth.

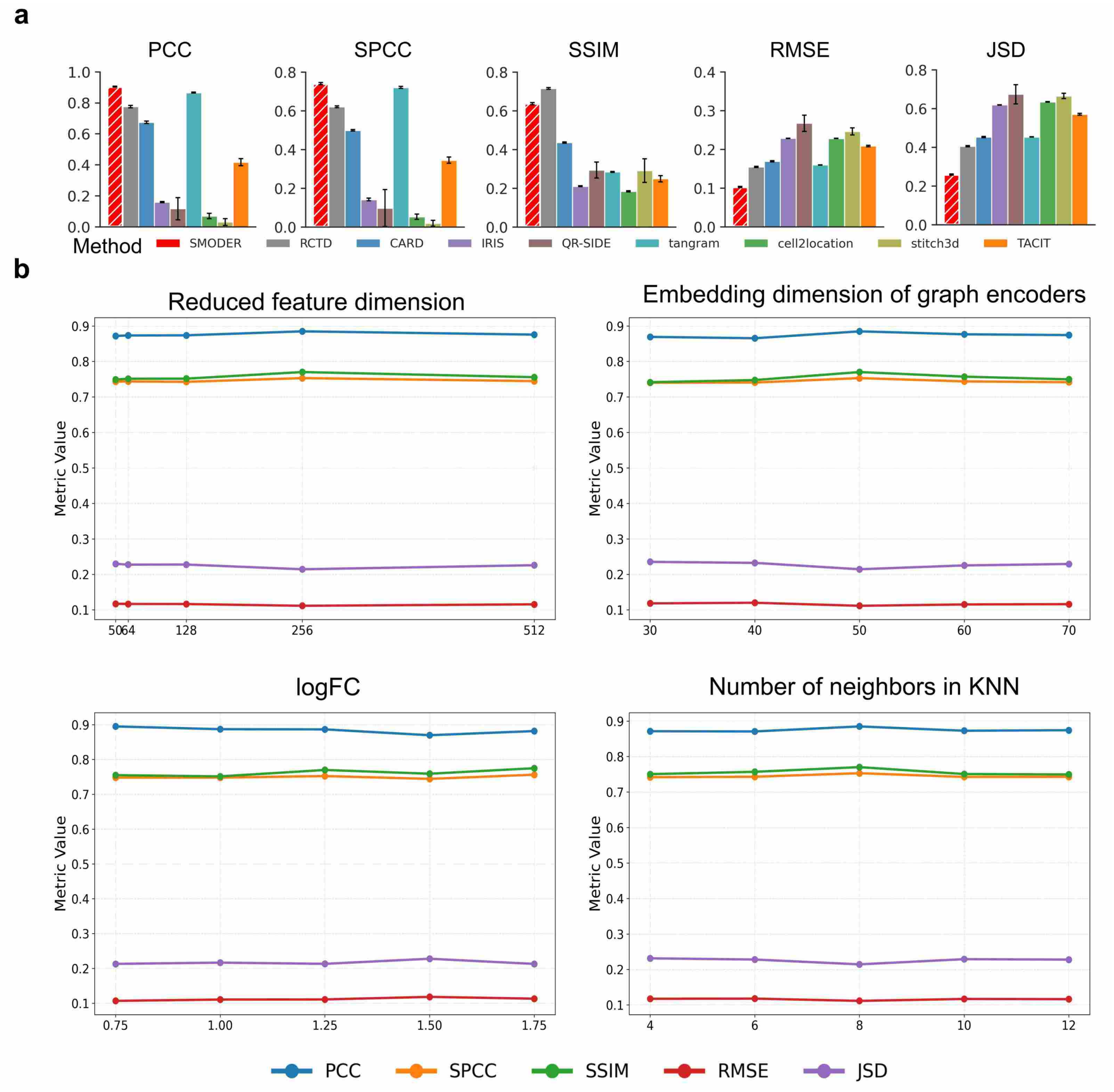

Supplementary Figure S4: **Quantitative comparative analysis and robustness analysis in simulated data. a**, Quantitative evaluation in simulated human CNS innate immune dataset (RNA+Protein) across five complementary metrics over five data generation repeats, including PCC, SPCC, SSIM, RMSE, and JSD. **b**, Sensitivity analysis in simulated human melanoma data (RNA+ATAC). Line plots of average PCC, SPCC, SSIM, RMSE, and JSD values across different values of parameter in SMODER over five data generation repeats, including embedding dimension of graph encoder, the reduced feature dimension of each modality, the $\log_2$ fold-change threshold for selecting informative genes, and the graph neighborhood size.

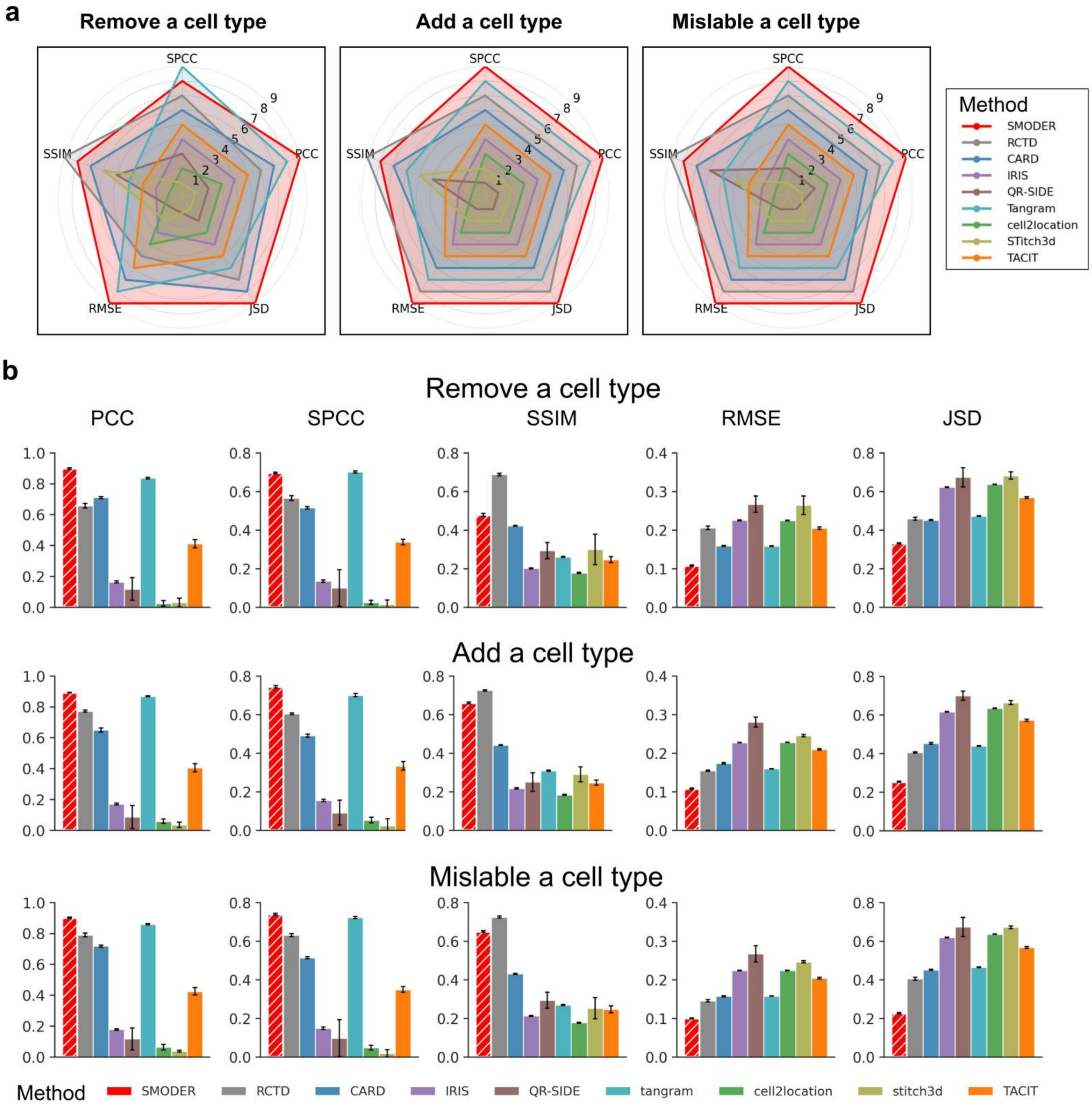

Supplementary Figure S5: **Robustness to reference misspecification under three scenarios in a simulated human CNS innate immune dataset (RNA+protein). a**, Radar plots summarizing method rankings across evaluation metrics, where larger enclosed areas indicate better overall performance. SMODER consistently achieves superior and stable performance across all perturbation settings. **b**, Quantitative evaluation across five complementary metrics over five independent data-generation repeats, including PCC, SPCC, SSIM, RMSE, and JSD.

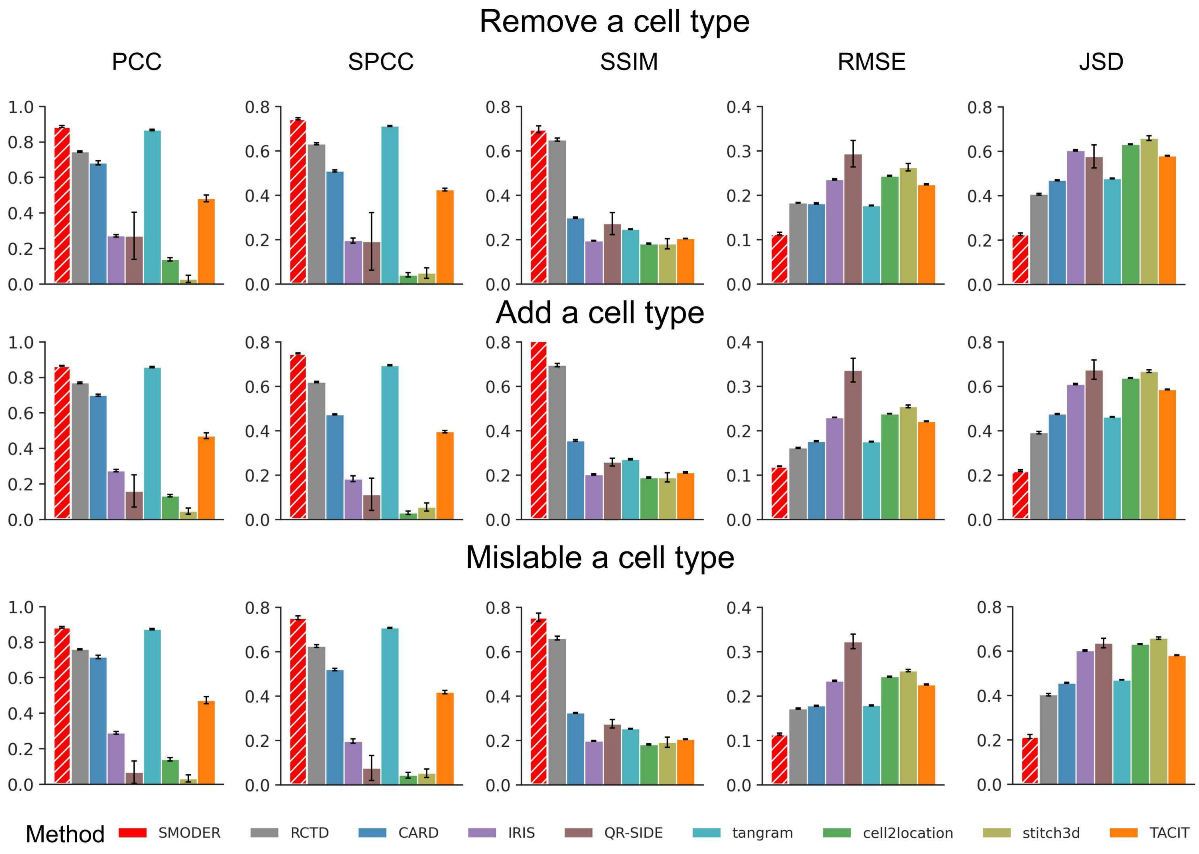

Supplementary Figure S6: **Robustness to reference misspecification under three scenarios in simulated human melanoma dataset (RNA+ATAC).** Quantitative evaluation across five complementary metrics over five data generation repeats, including PCC, SPCC, SSIM, RMSE, and JSD.

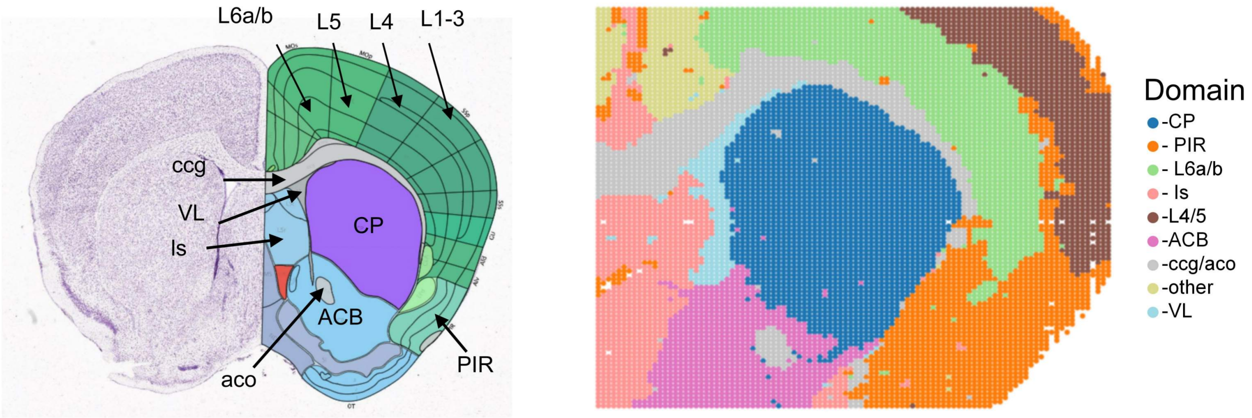

Supplementary Figure S7: **Manual annotations of mouse brain spatial CUT&Tag—RNA-seq data for the H3K27ac mark.** Left, reference structural annotations of a coronal section of the P56 mouse brain from the Allen Brain Atlas. Right, spatial heatmaps of nine manual-annotated domains.

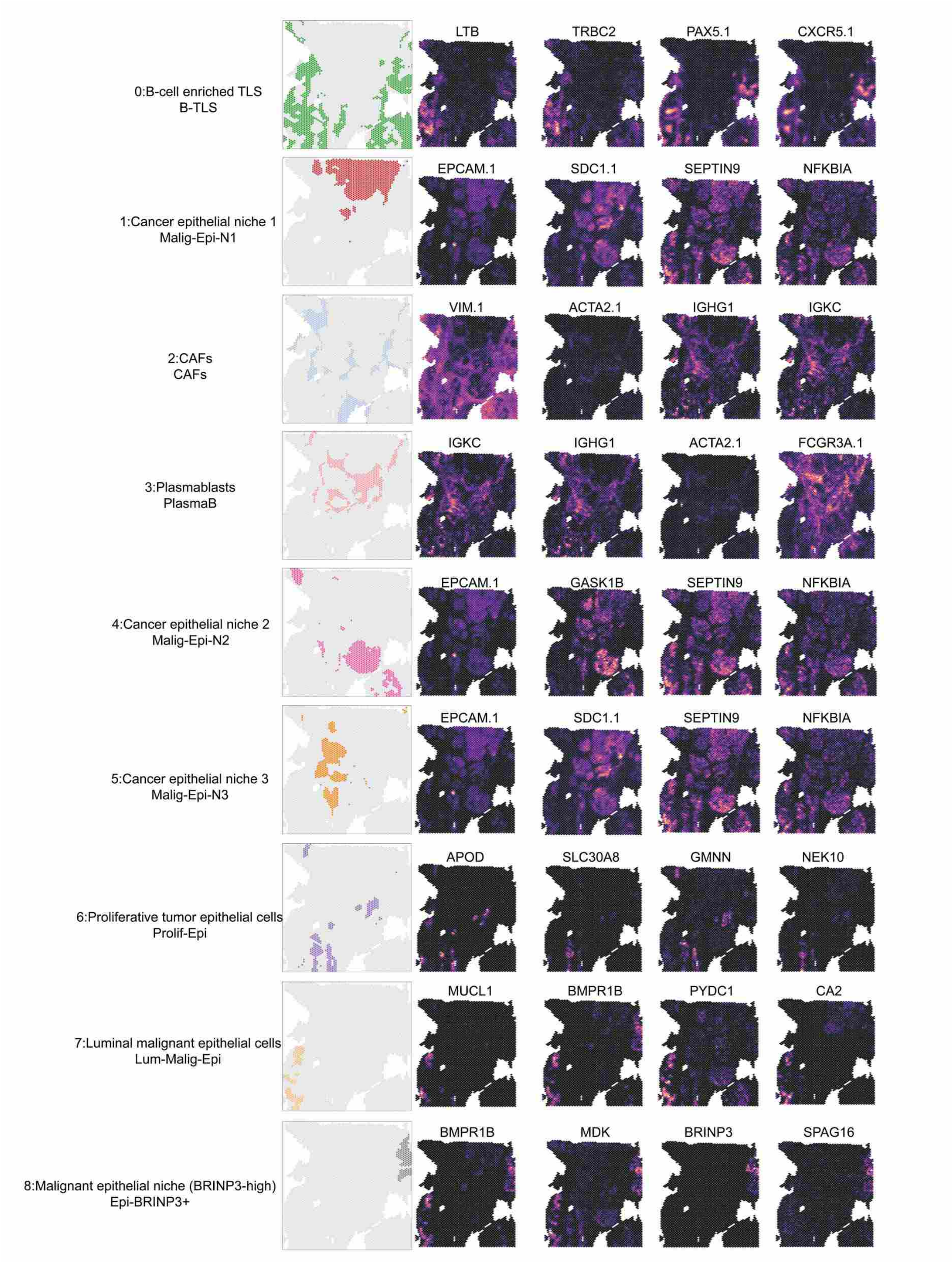

Supplementary Figure S8: **Manual annotations of human breast cancer CytAssist RNA–protein co-profiling data.** Spatial heatmaps showing each of nine manually annotated spatial domains alongside the expression patterns of their corresponding marker genes.

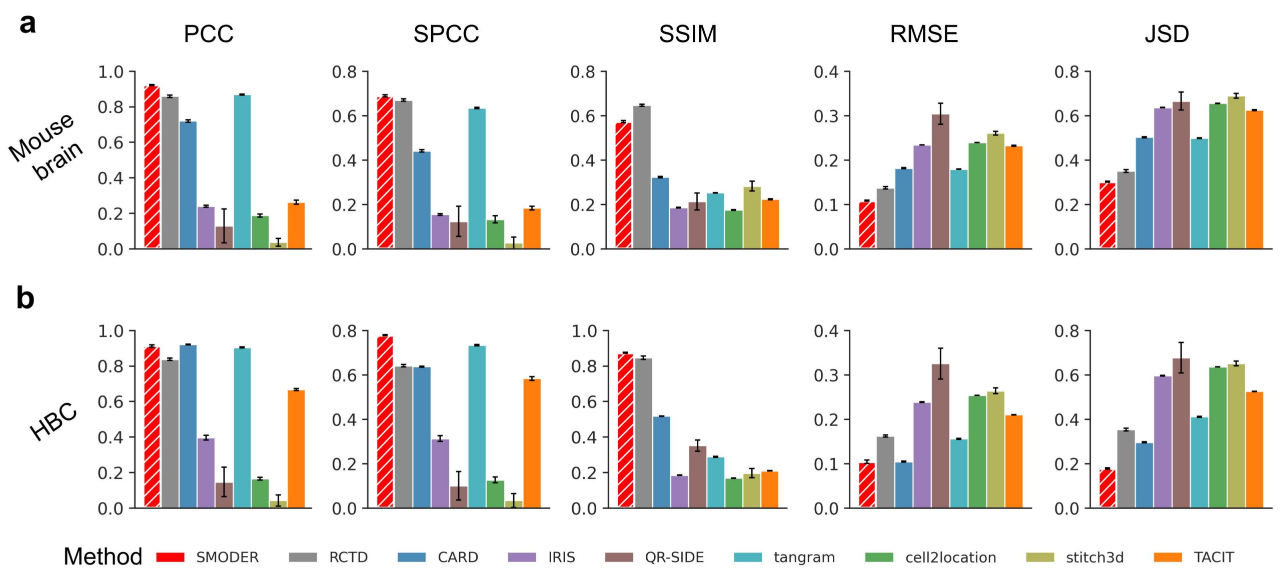

Supplementary Figure S9: **Quantitative evaluation in simulated mouse brain (RNA+Histone) and human breast cancer (RNA+protein) across five complementary metrics over five data generation repeats.** Barplots of average metric values obtained by SMODER and compared methods, including PCC, SPCC, SSIM, RMSE, and JSD. **a**, mouse brain data. **b**, human breast cancer data.

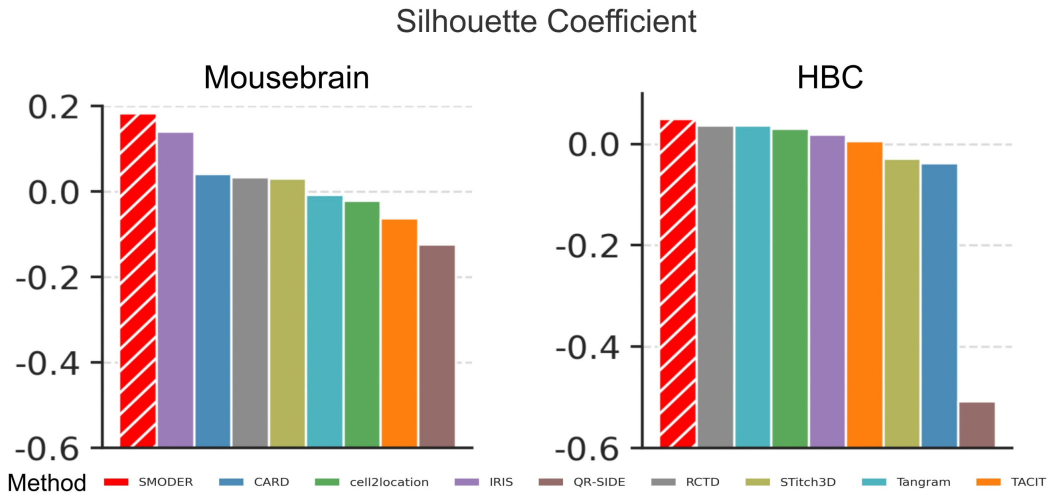

Supplementary Figure S10: **Comparative analysis of deconvolution for SMODER and competing methods in two real datasets.** Barplots of average silhouette coefficients obtained by SMODER and compared methods. **Left**, mouse brain data; **Right**, human breast cancer data.

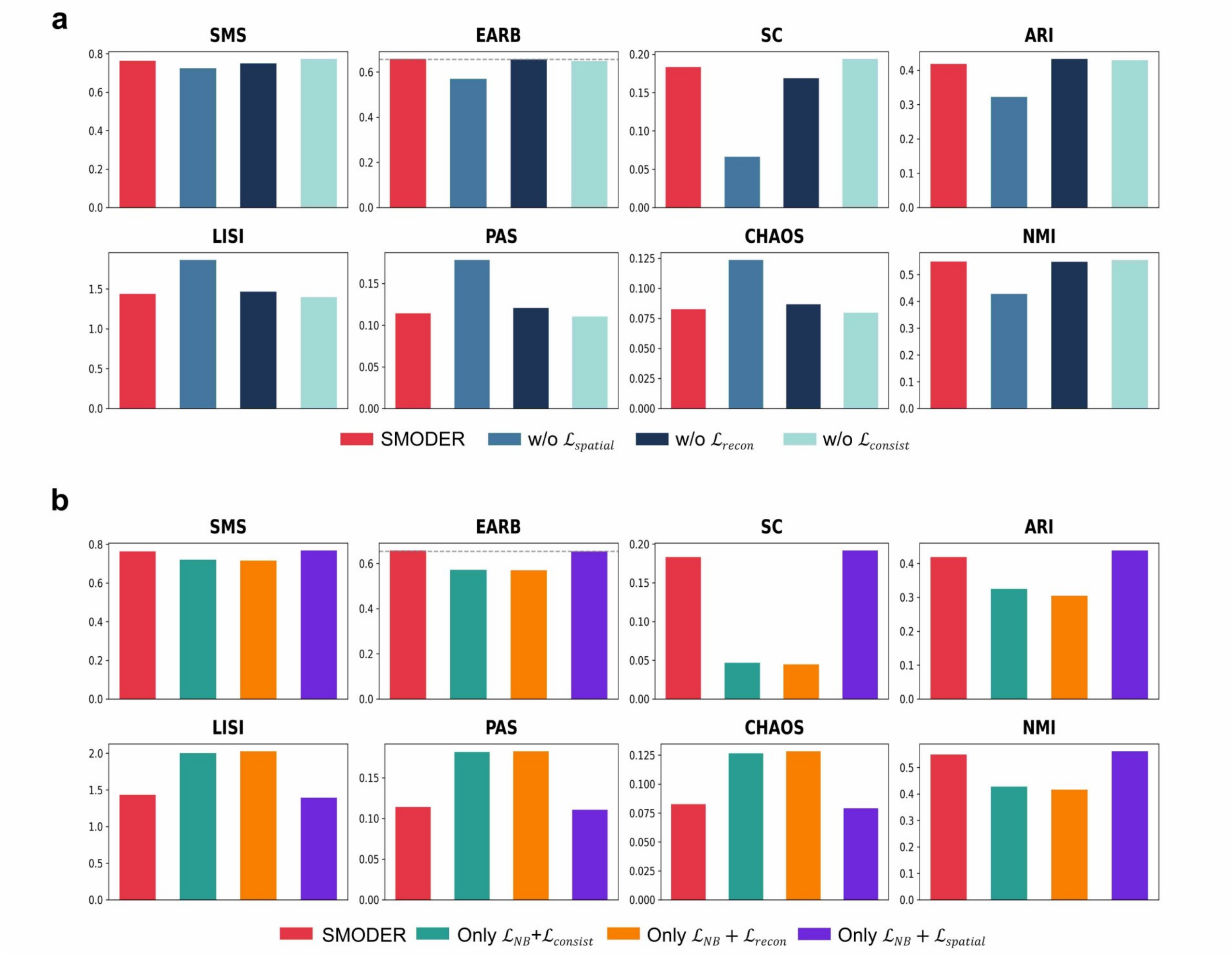

Supplementary Figure S11: **Ablation experiments of SMODER on the H3K27ac mouse brain spatial CUT&Tag—RNA-seq data.** Quantitative evaluation of SMODER and its variants using SMS, BEAR, silhouette coefficient (SC), ARI, and NMI metrics, where higher values indicate better performance, as well as LISI, PAS, and CHAOS metrics, where lower values indicate better performance. **a**, Ablation analysis of individual loss components in SMODER. Barplots from left to right: SMODER with complete four loss components, w/o $\mathfrak{L}_{spatial}$, w/o $\mathfrak{L}_{recon}$, w/o $\mathfrak{L}_{consist}$. **b**, Analysis of combination effects among loss components in SMODER with complete four loss components. Barplots from left to right: SMODER, $\mathfrak{L}_{NB}$ + only $\mathfrak{L}_{consist}$, only $\mathfrak{L}_{recon}$, and only $\mathfrak{L}_{spatial}$, respectively.

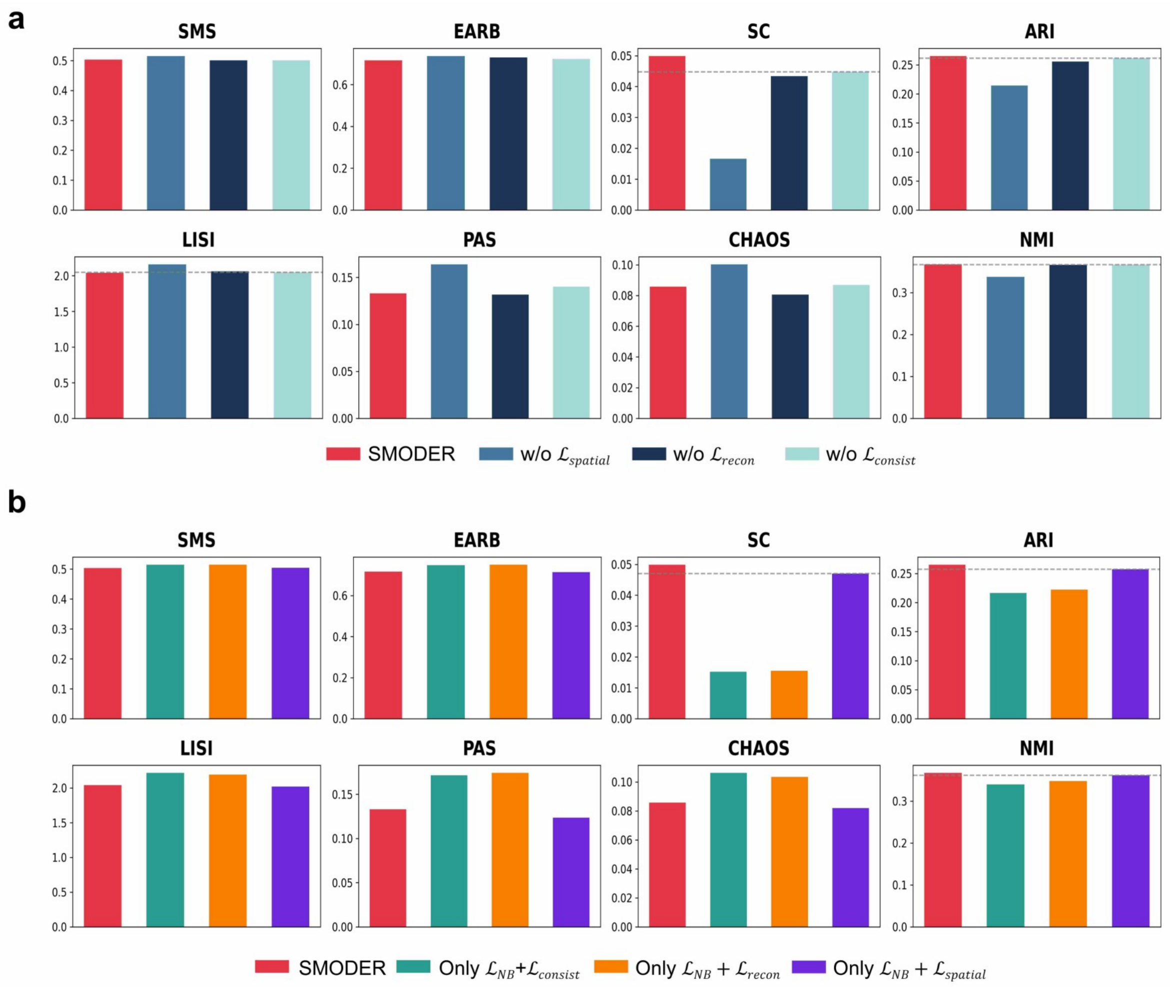

Supplementary Figure S12: **Ablation experiments of SMODER on the human breast cancer CytAssist RNA—protein co-profiling data.** Quantitative evaluation of SMODER and its variants using SMS, BEAR, silhouette coefficient (SC), ARI, and NMI metrics, where higher values indicate better performance, as well as LISI, PAS, and CHAOS metrics, where lower values indicate better performance. **a**, Ablation analysis of individual loss components in SMODER. Barplots from left to right: SMODER with complete four loss components, w/o $\mathfrak{L}_{spatial}$, w/o $\mathfrak{L}_{recon}$, and w/o $\mathfrak{L}_{consist}$. **b**, Analysis of combination effects among loss components in SMODER. Barplots from left to right: baseline SMODER, $\mathfrak{L}_{NB}$ + only $\mathfrak{L}_{consist}$, only $\mathfrak{L}_{recon}$, and only $\mathfrak{L}_{spatial}$, respectively.

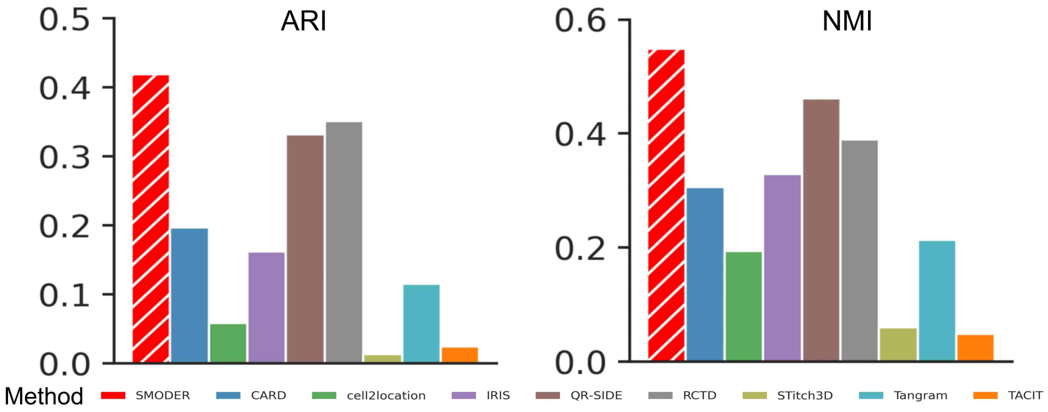

Supplementary Figure S13: **Quantitative comparison between annotated spatial domains and inferred dominant cell types in mouse brain spatial CUT&Tag–RNA-seq data.** Bar plots of ARI and NMI values computed from dominant cell types inferred by SMODER and competing methods.

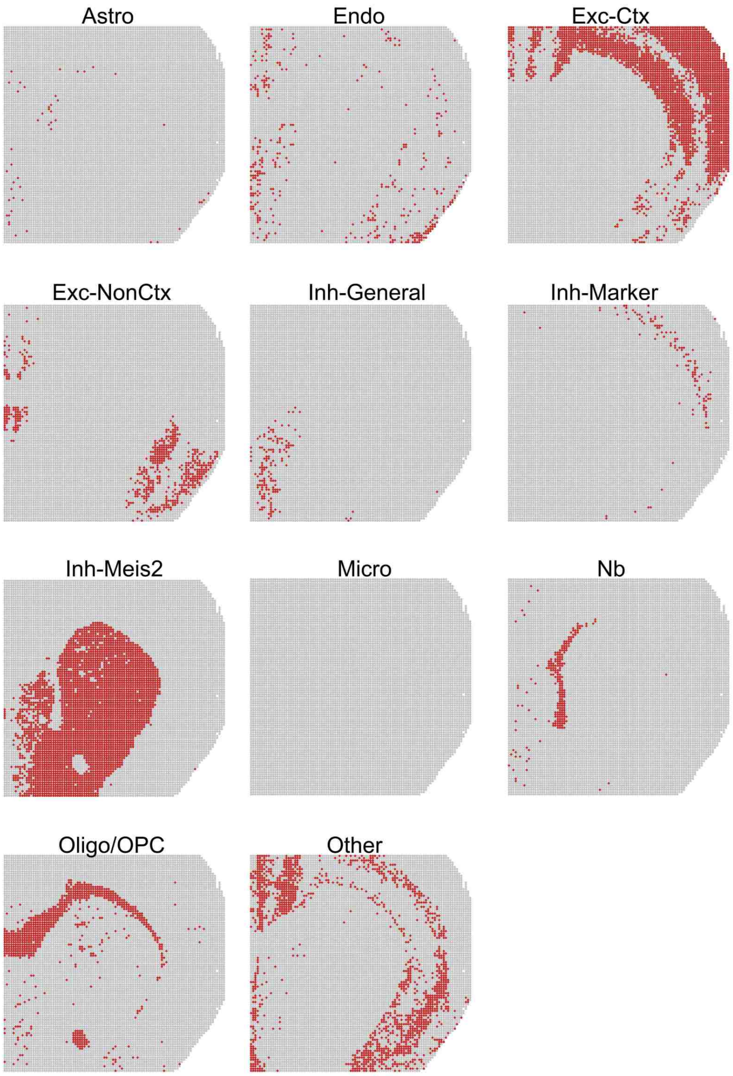

Supplementary Figure S14: **Spatial distribution of dominant cell types inferred by SMODER in mouse brain spatial CUT&Tag–RNA-seq data.** Spatial heatmap for each dominant cell type obtained from SMODER deconvolution.

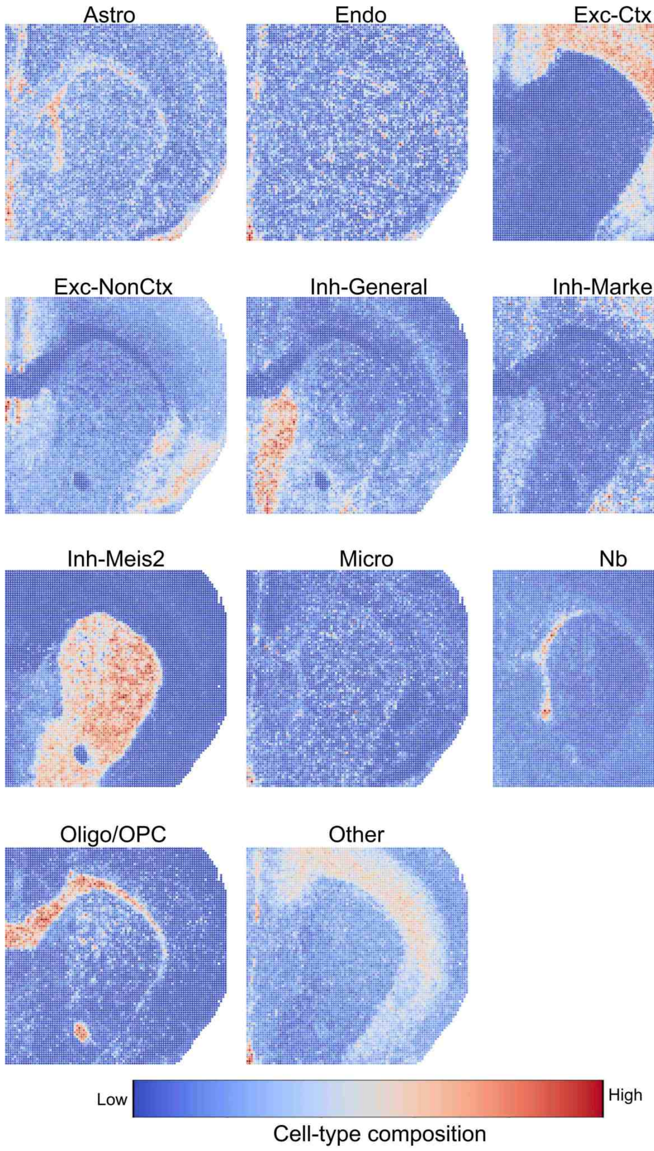

Supplementary Figure S15: **Spatial distributions of cell-type composition inferred by SMODER in mouse brain spatial CUT&Tag–RNA-seq data.** Spatial heatmaps show the inferred composition of each cell type obtained from SMODER deconvolution, visualized separately for individual cell types.

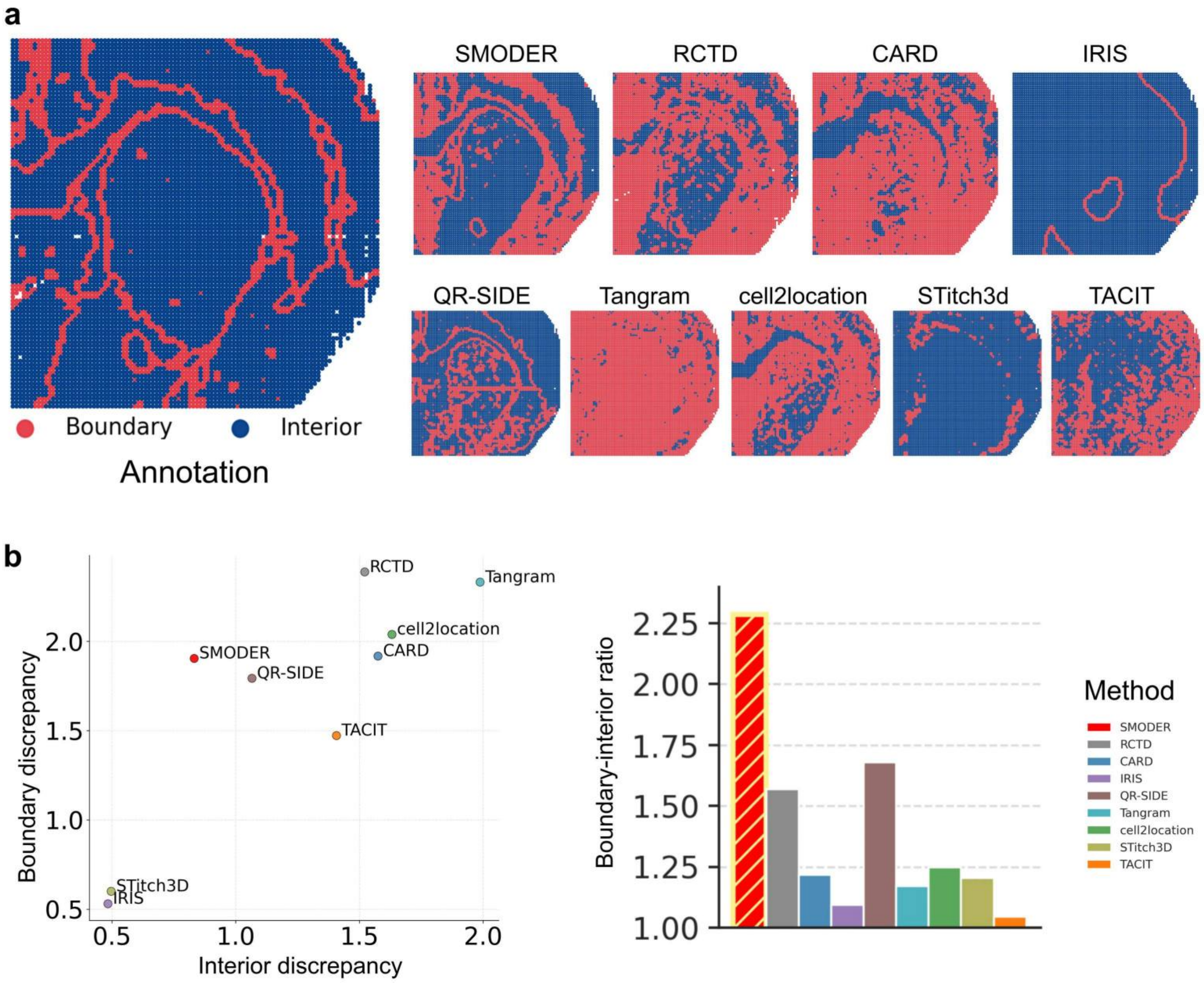

Supplementary Figure S16: **Spatial distribution and quantitative analysis of boundaries in deconvolution results.** Spots are divided into boundary and interior units. The boundary distinguishability of deconvolution outputs from SMODER and competing methods is compared, with further quantitative analysis performed. **a**, Spatial heatmaps show the boundary distribution of manual annotations, as well as the spatial distribution of boundaries from deconvolution outputs generated by SMODER and other competing methods, stratified by dominant cell types. **b**, The left scatter plot quantifies the degree of dominant cell type discrepancy at boundary and interior spots between SMODER and competing methods, while the right bar plot presents the boundary-interior ratio scores. Higher values indicate better performance.

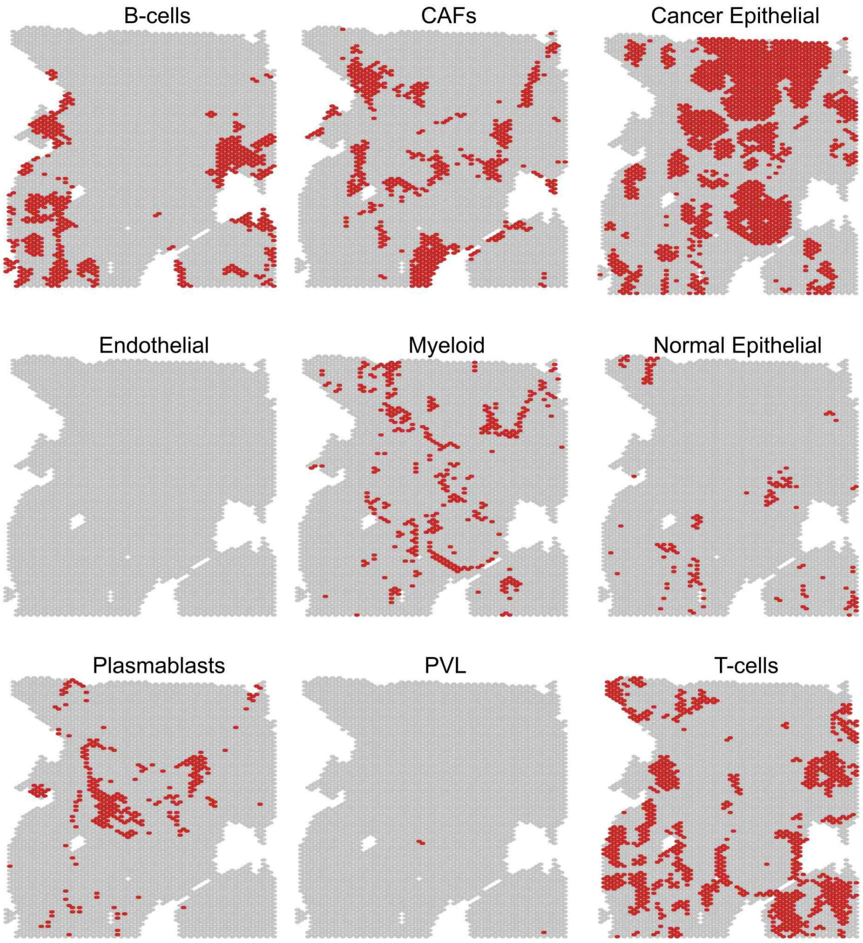

Supplementary Figure S17: **Spatial distribution of dominant cell types inferred by SMODER in human breast cancer CytAssist RNA–protein co-profiling data.** Spatial heatmaps show the dominant cell type assignments obtained from SMODER deconvolution, visualized separately for each cell type.

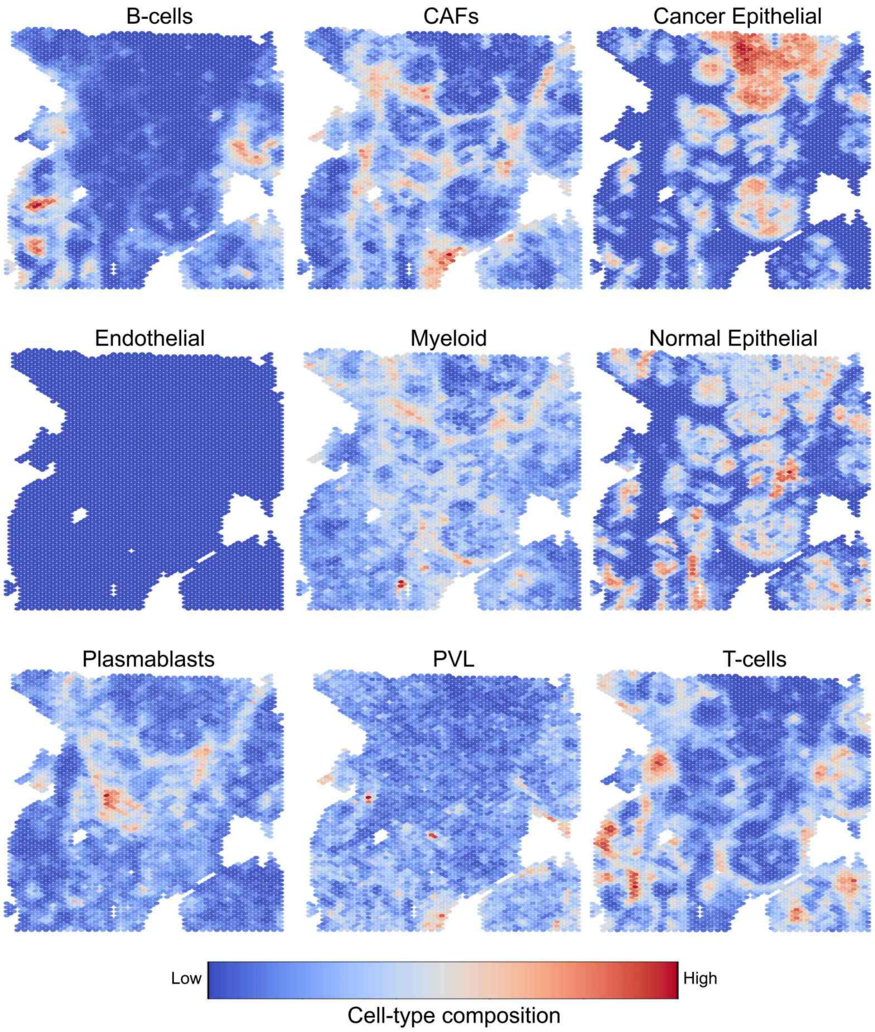

Supplementary Figure S18: **Spatial distributions of cell-type composition inferred by SMODER in human breast cancer CytAssist RNA–protein co-profiling data.** Spatial heatmaps show the inferred composition of each cell type obtained from SMODER deconvolution, visualized separately for individual cell types.

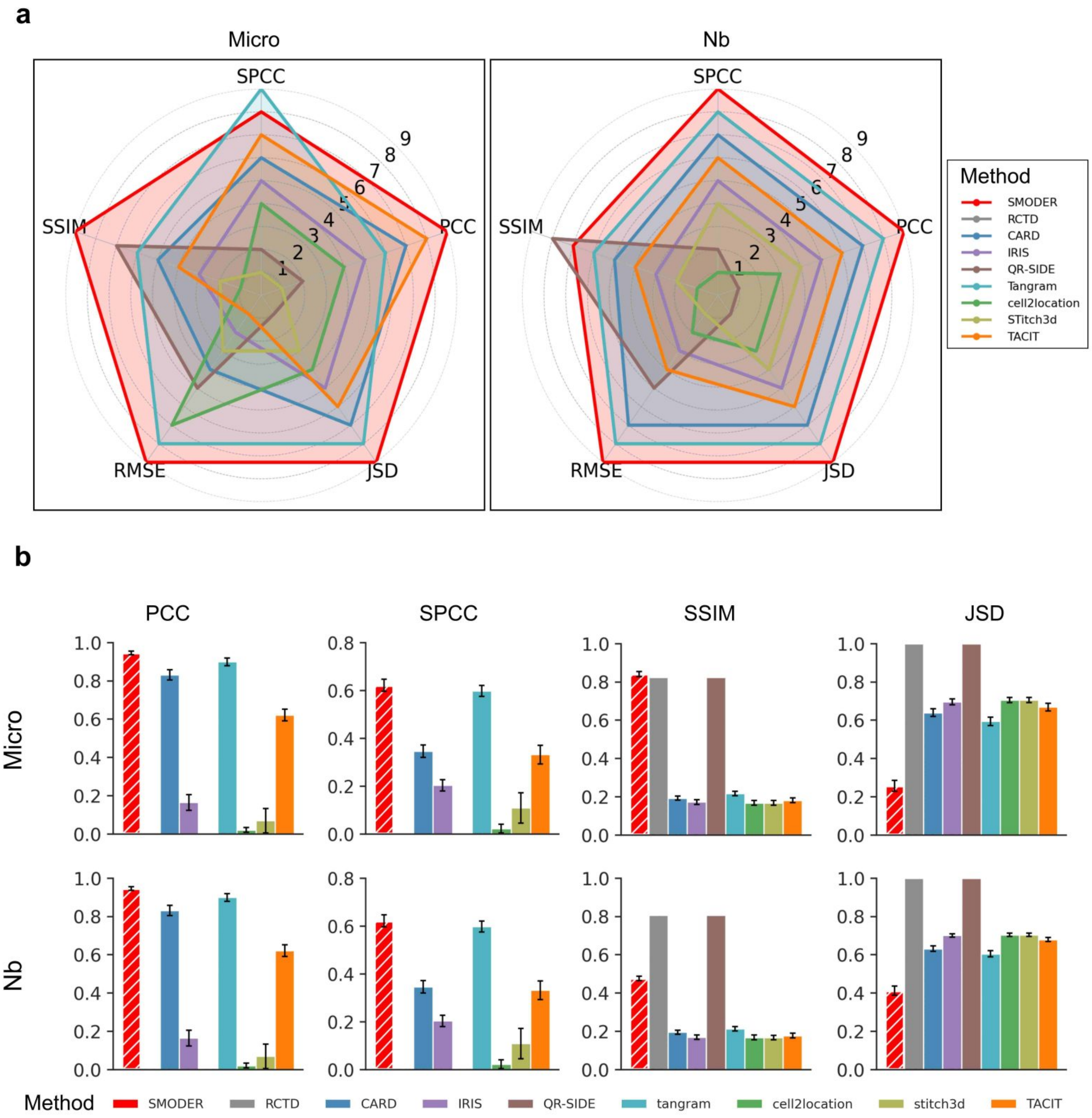

Supplementary Figure S19: **Quantitative analysis of deconvolution performance for rare cell types in mouse brain spatial CUT&Tag–RNA-seq data for the H3K27ac mark.** **a**, Radar plots illustrating method rankings across evaluation metrics for two rare cell types (microglia (Micro) and neuroblasts (Nb)). Larger enclosed areas indicate better overall performance. SMODER achieves superior and stable performance for both cell types. **b**, Bar plots showing quantitative performance across five independent data-generation replicates, evaluated using four metrics: PCC, SPCC, SSIM, and JSD.

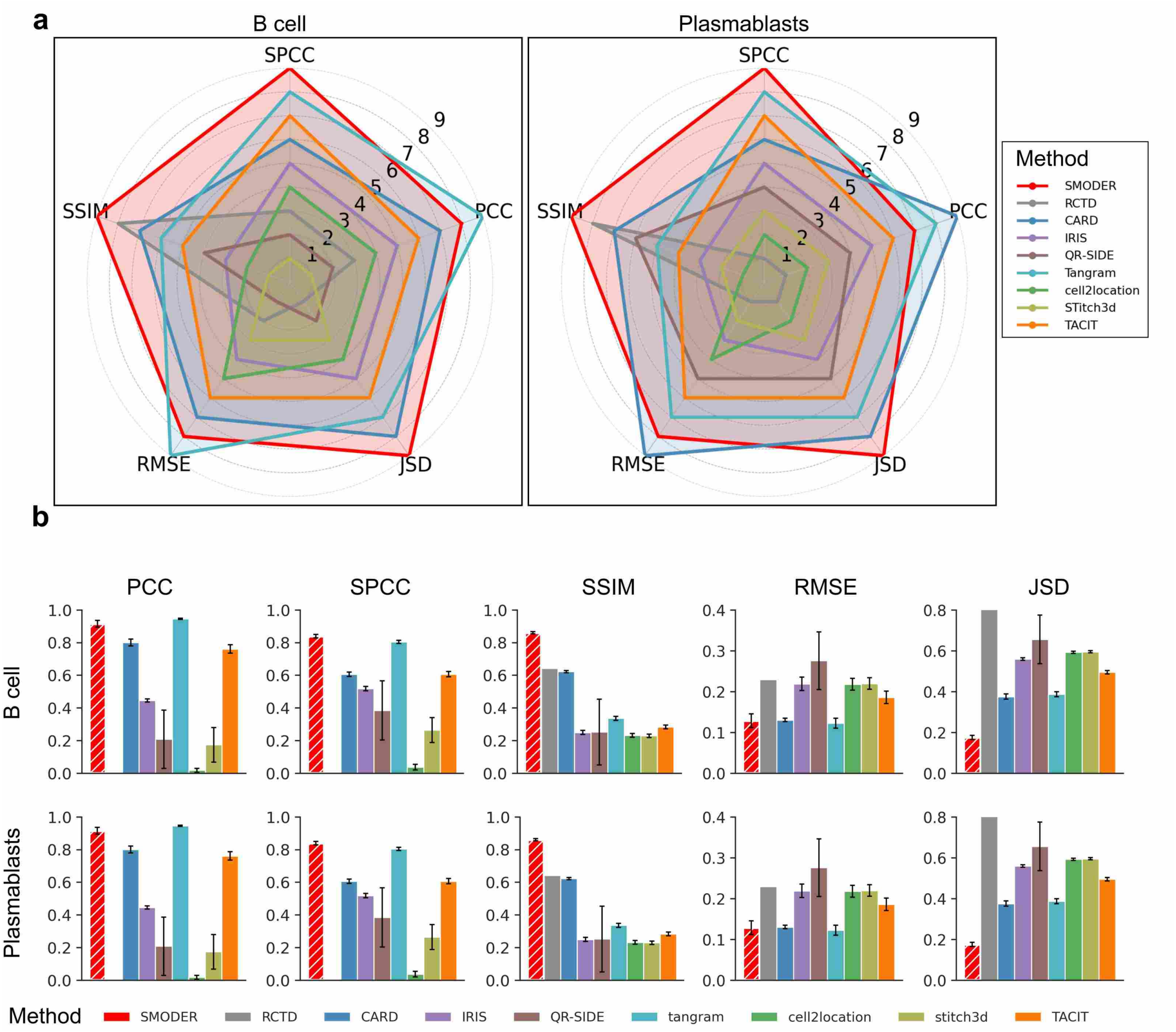

Supplementary Figure S20: **Quantitative analysis of deconvolution performance for rare cell types in human breast cancer CytAssist RNA–protein co-profiling data.** **a**, Radar plots illustrating method rankings across evaluation metrics for two rare cell types (B cells and plasmablasts). Larger enclosed areas indicate better overall performance. **b**, Bar plots showing quantitative performance across five independent data-generation replicates, evaluated using five metrics: PCC, SPCC, SSIM, RMSE, and JSD.

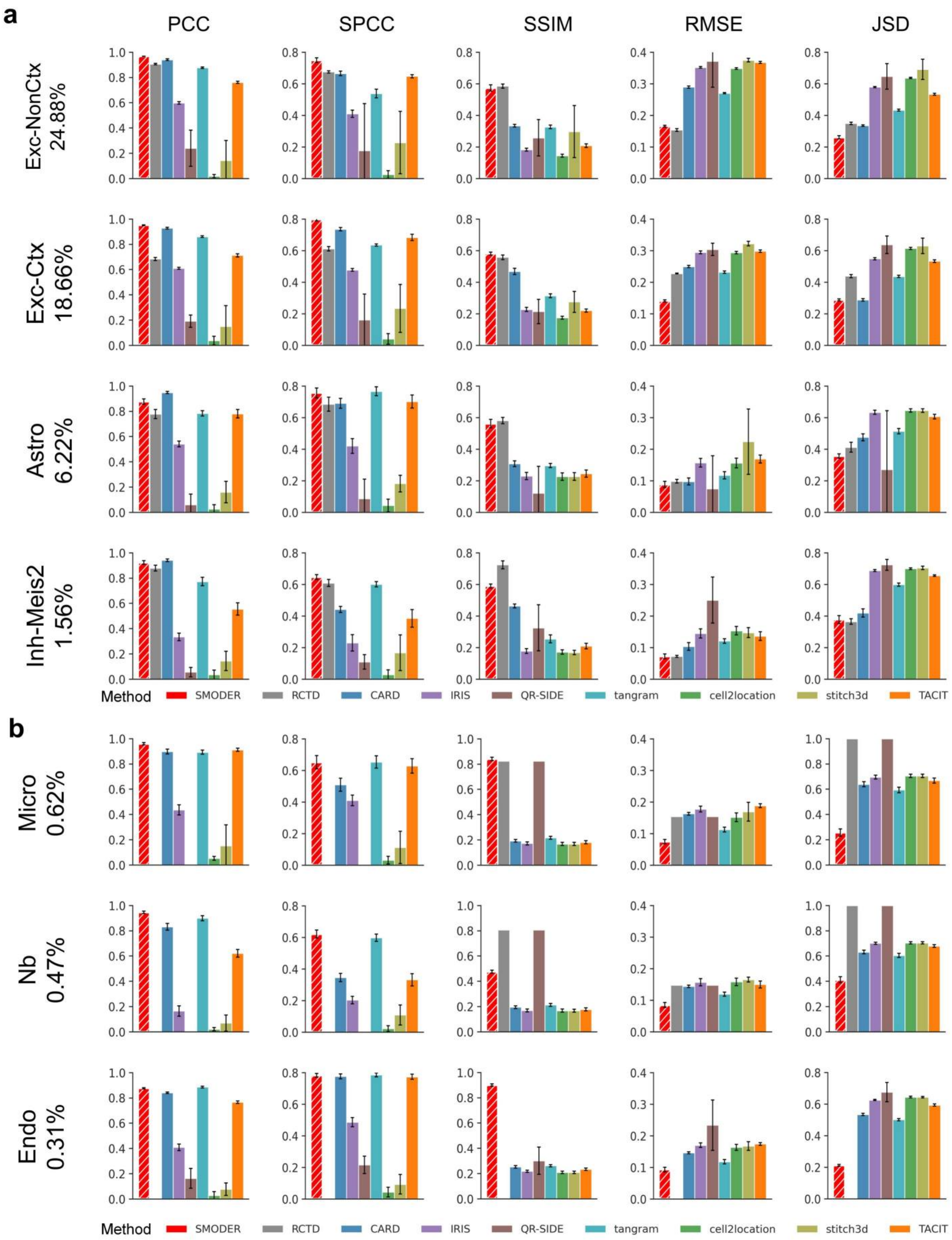

Supplementary Figure S21: **Quantitative evaluation of deconvolution performance across cell types with different abundance levels in mouse brain spatial CUT&Tag–RNA-seq data (H3K27ac mark).** **a**, Bar plots showing performance across five metrics (PCC, SPCC, SSIM, RMSE, and JSD) for high-abundance cell types in the single-cell reference data. SMODER exhibits stable and consistently strong performance. **b**, Bar plots showing performance across the same five metrics for low-abundance cell types. SMODER demonstrates markedly improved overall performance for low-abundance cell types.

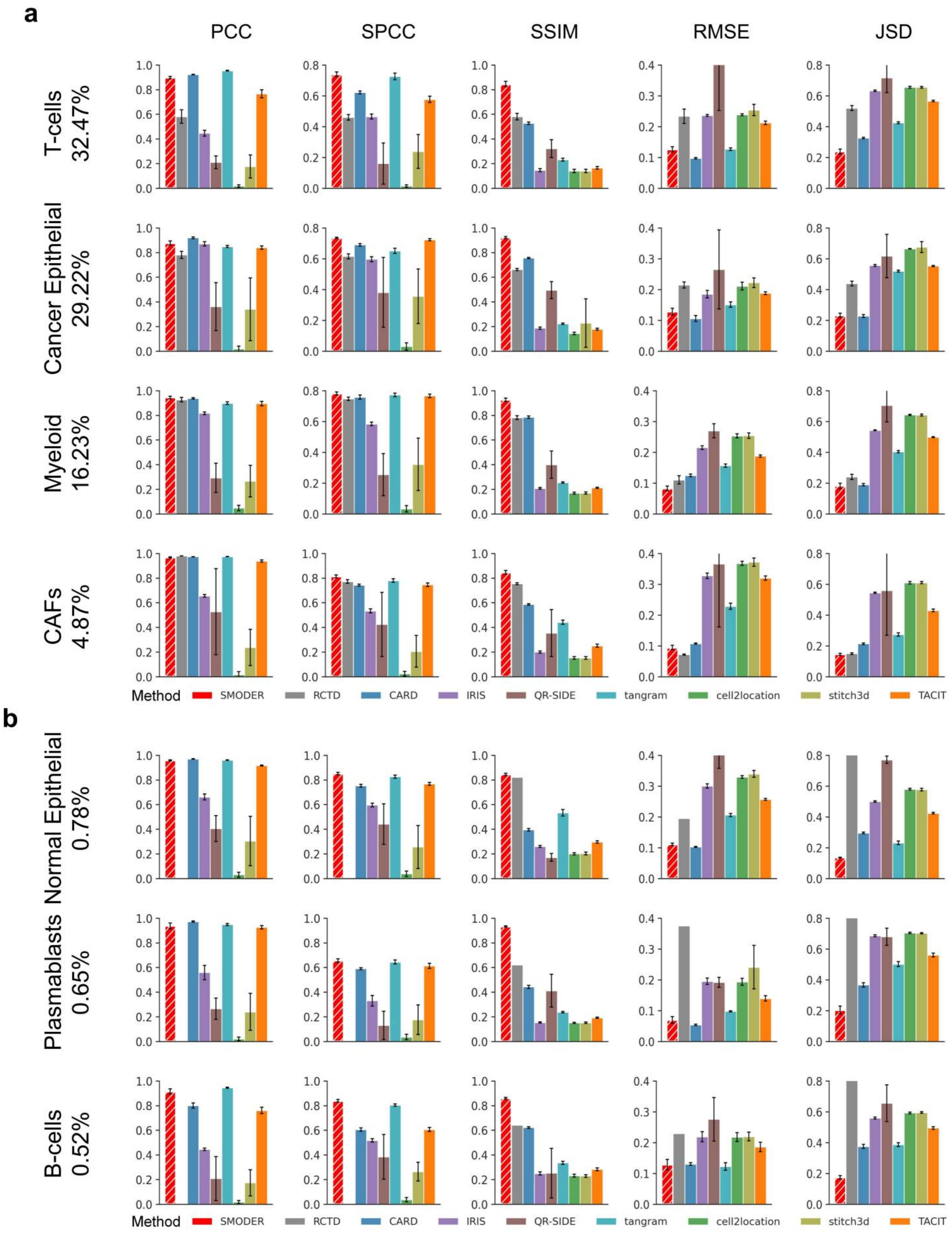

Supplementary Figure S22: **Quantitative evaluation of deconvolution across cell types with different abundance levels in human breast cancer CytAssist RNA–protein co-profiling data.** **a**, Bar plots showing performance across five metrics (PCC, SPCC, SSIM, RMSE, and JSD) for high-abundance cell types in the single-cell reference data. SMODER exhibits stable and consistently strong performance. **b**, Bar plots showing performance across the same five metrics for low-abundance cell types. SMODER demonstrates markedly improved overall performance.

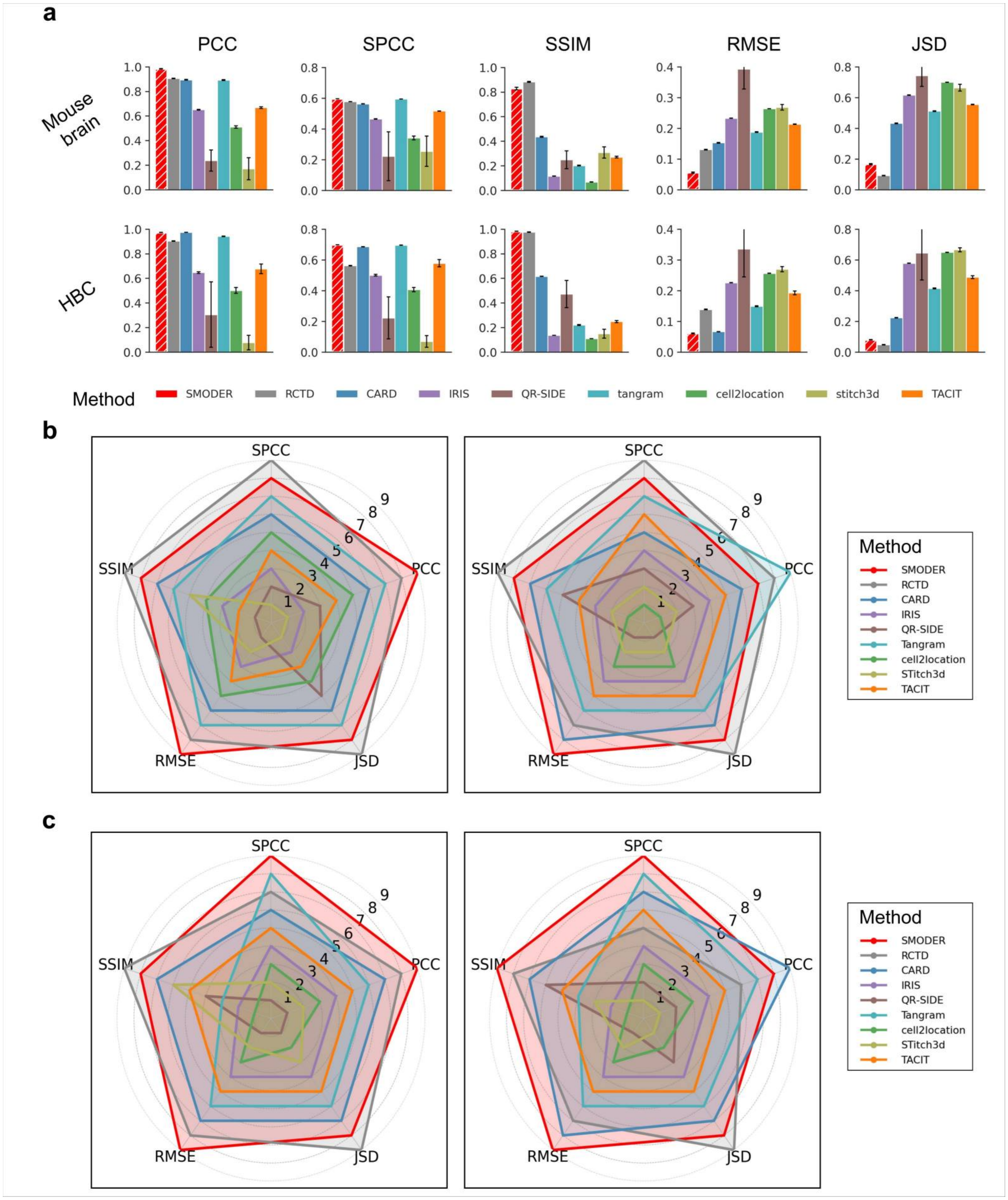

a
PCC
SPCC
SSIM
RMSE
JSD
Mouse brain
HBC
Method
SMODER
RCTD
CARD
IRIS
QR-SIDE
tangram
cell2location
stitch3d
TACIT
b
c
Tangram
STitch3d

Supplementary Figure S23: **Quantitative analysis of deconvolution performance in regions with sparse and dense expression levels. a**, Bar plots showing deconvolution performance on the top 10% expression-sparse spots across five independent replicates, evaluated using five metrics (PCC, SPCC, SSIM, RMSE, and JSD) for mouse brain (upper) and human breast cancer (lower) datasets. **b**, Radar plots showing method rankings on the top 10% expression-dense spots across five replicates. SMODER achieves performance comparable to RCTD and other competing methods on both mouse brain (left) and human breast cancer (right) datasets. **c**, Radar plots illustrating method rankings on the top 10% expression-sparse spots across five replicates. Larger enclosed areas indicate better performance. SMODER achieves superior performance on both mouse brain (left) and human breast cancer (right) datasets.

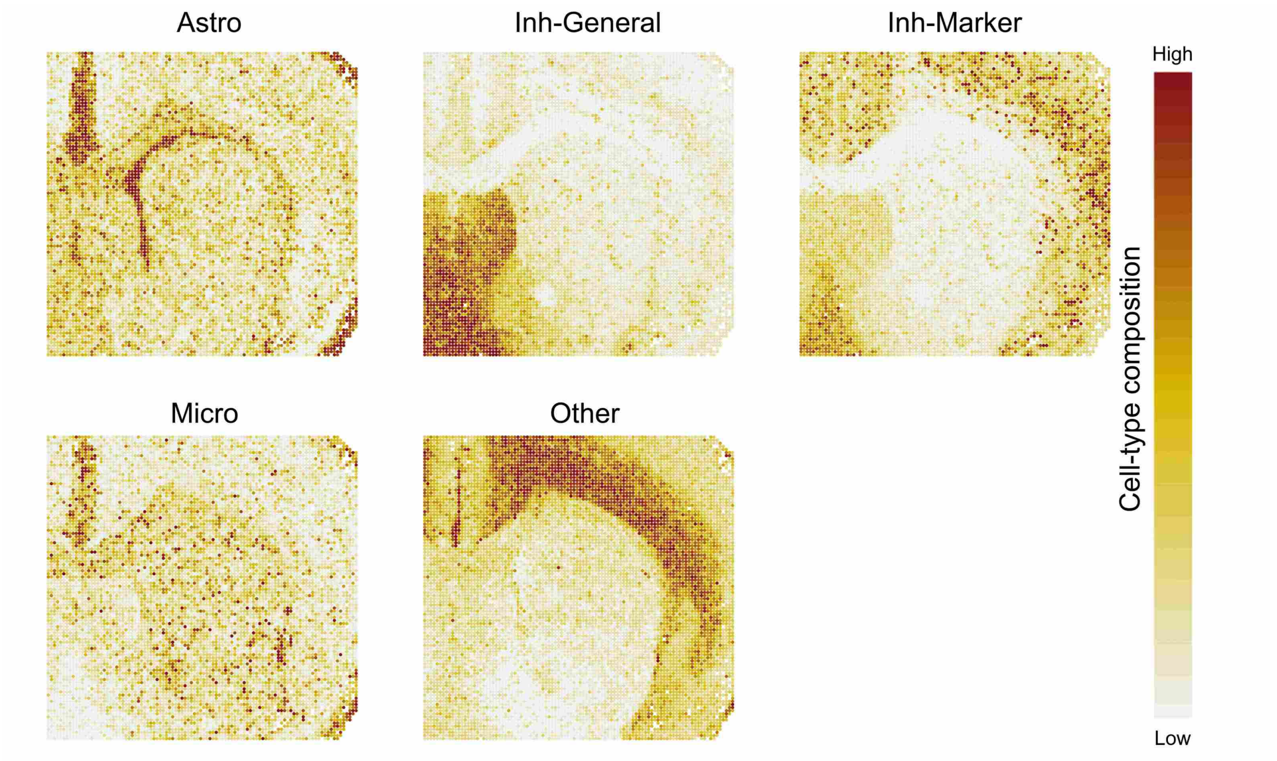

Supplementary Figure S24: **Spatial distributions of cell-type composition inferred by SMODER in mouse brain spatial CUT&Tag–RNA-seq data for the H3K27me3 mark.** Spatial heatmaps show the inferred composition of each cell type obtained from SMODER deconvolution, visualized separately for individual cell types not illustrated in the main text.

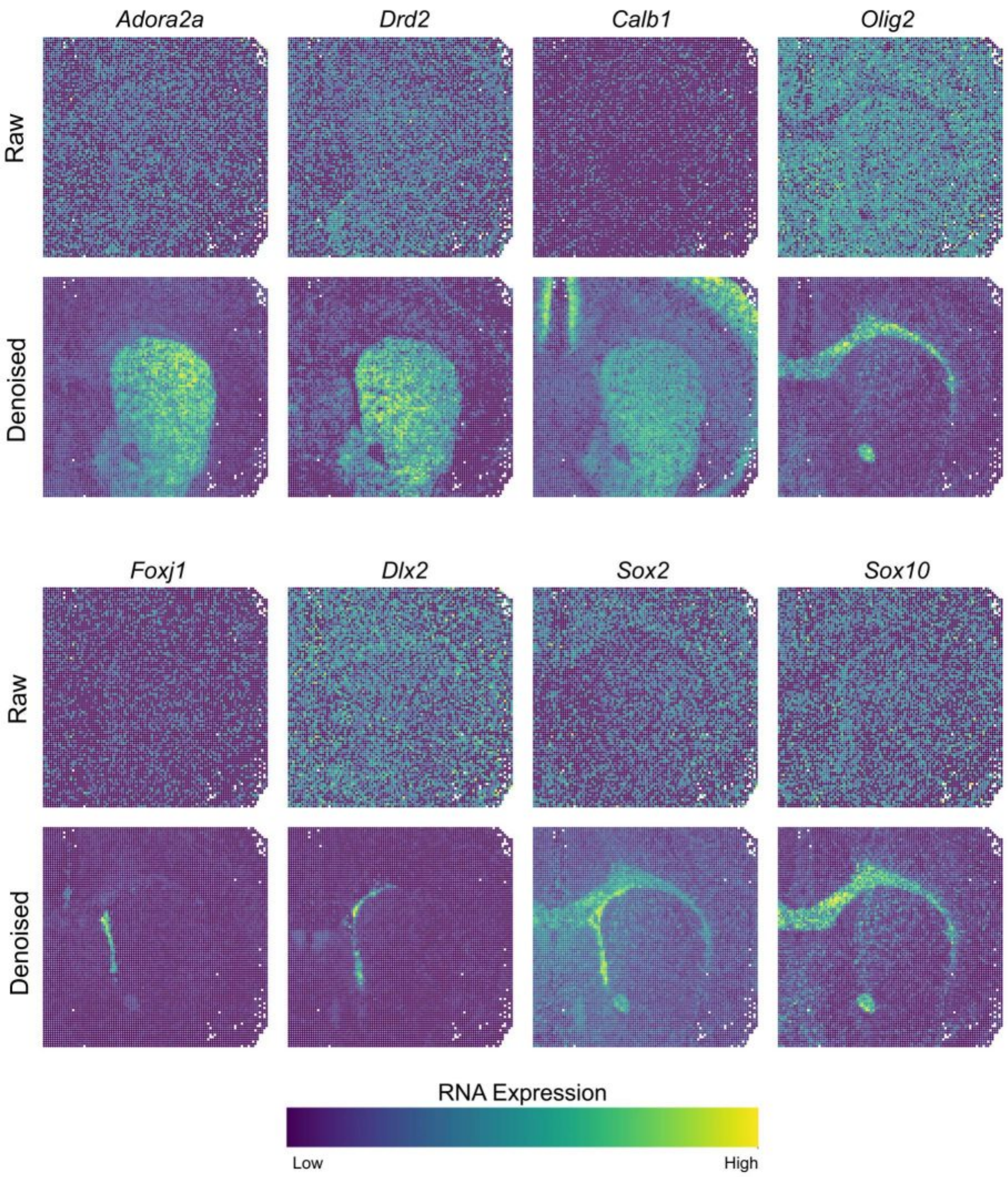

Supplementary Figure S25: **Spatial distribution of reconstructed spatial transcriptomic results in mouse brain spatial CUT&Tag–RNA-seq data for the H3K27me3 mark.** Spatial heatmaps show the reconstruction results of partial genes obtained by SMODER, with raw expression levels displayed above and reconstructed profiles below.

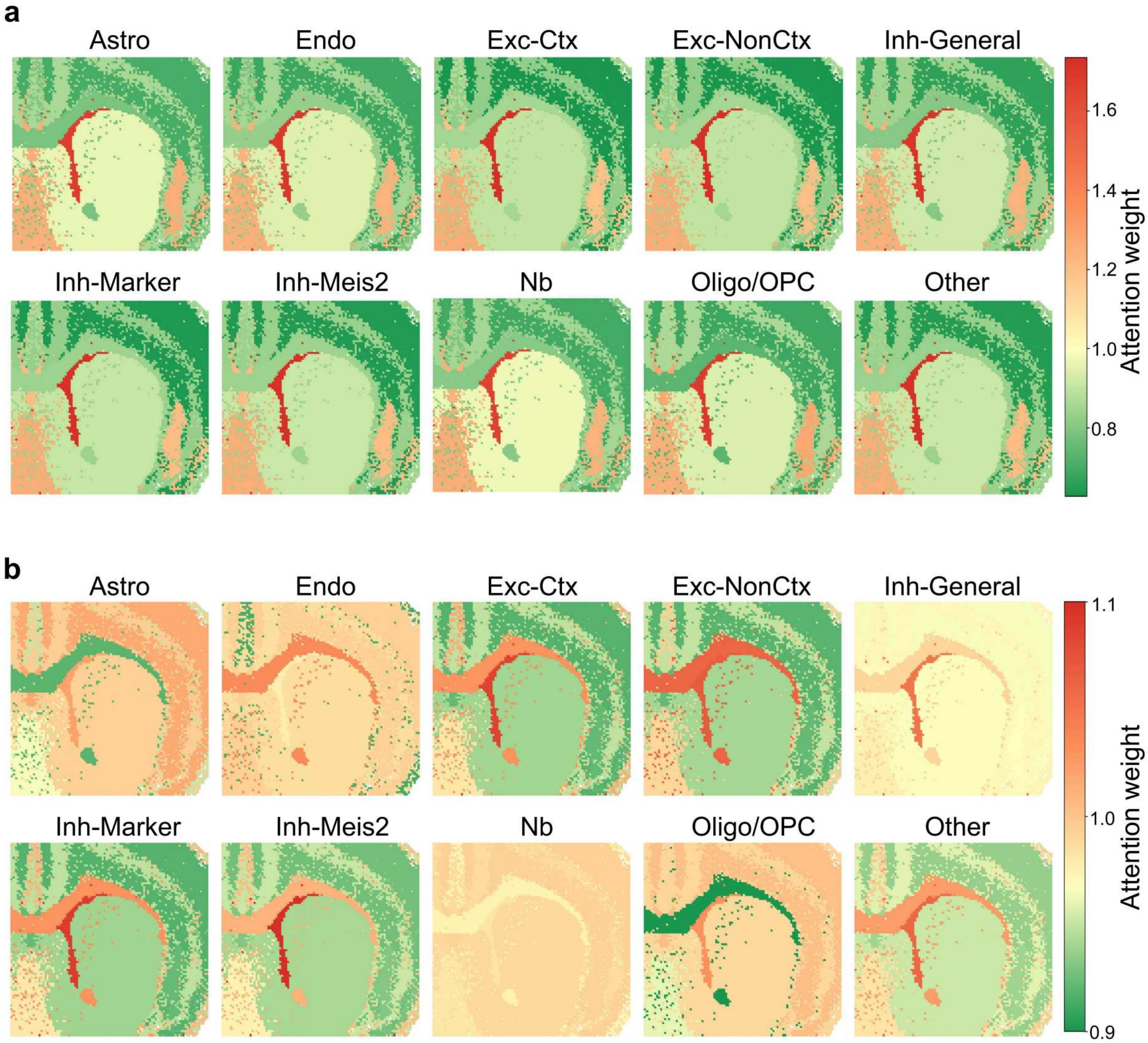

Supplementary Figure S26: **Dominant cell type-specific attention scores between dual modalities.** **a**, Spatial heatmaps show epigenome-to-RNA attention scores across different cell types, visualized separately for individual cell types. **b**, Spatial heatmaps show RNA-to-epigenome attention scores across different cell types, visualized separately for individual cell types.

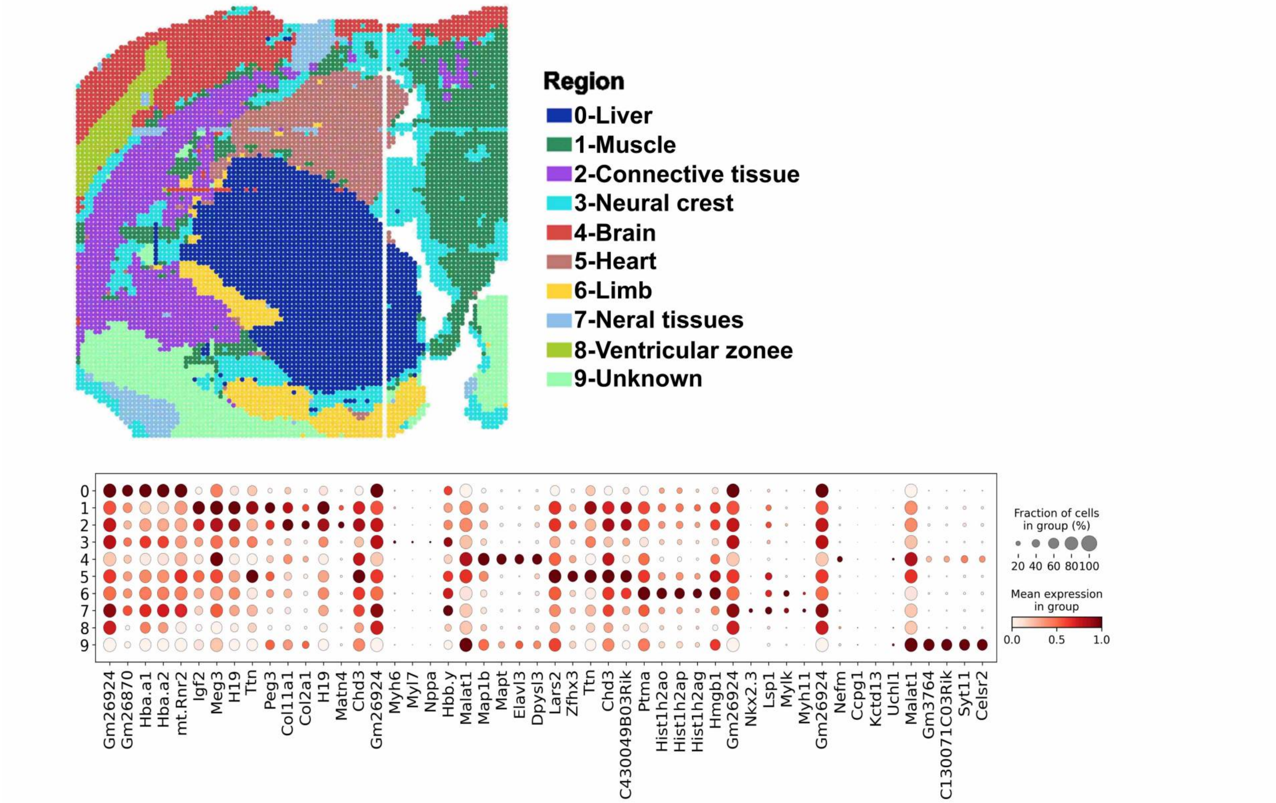

Supplementary Figure S27: **Expression characteristics of domain-specific marker genes in mouse embryonic spatial transcriptomic data.** Upper panel: spatial heatmap of manually annotated regions. Lower panel: dot plot illustrating the expression levels and composition of the top five marker genes across distinct anatomical domains.

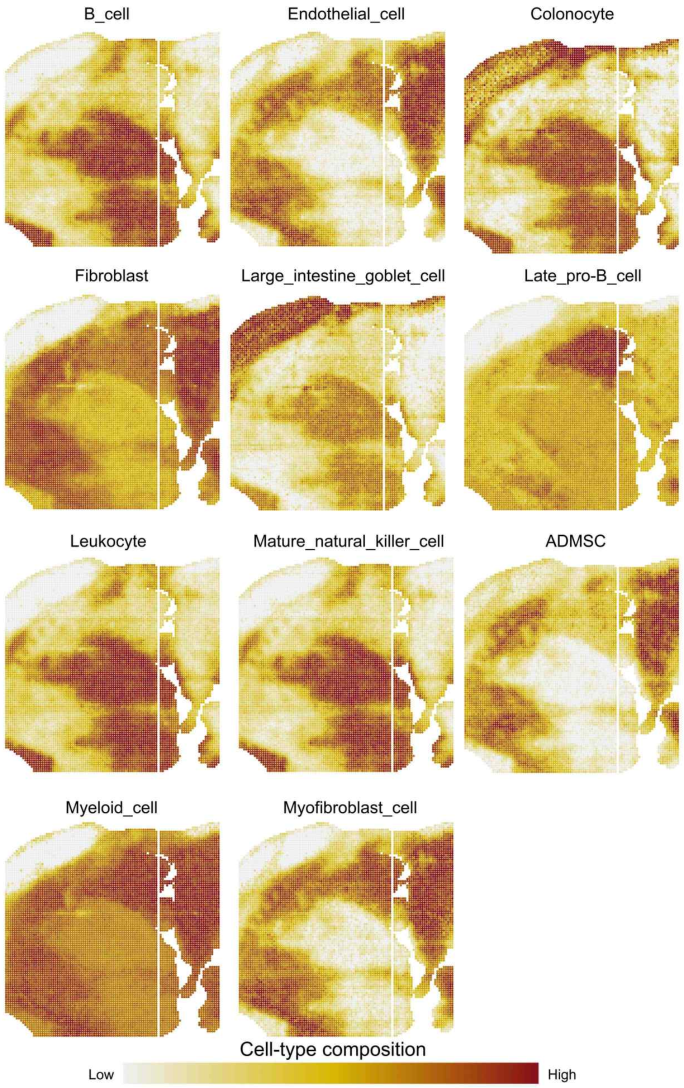

Supplementary Figure S28: **Spatial distributions of cell-type composition inferred by SMODER in mouse embryo data.** Spatial heatmaps show the inferred composition of each cell type obtained from SMODER deconvolution, visualized separately for individual cell types not illustrated in the main text.

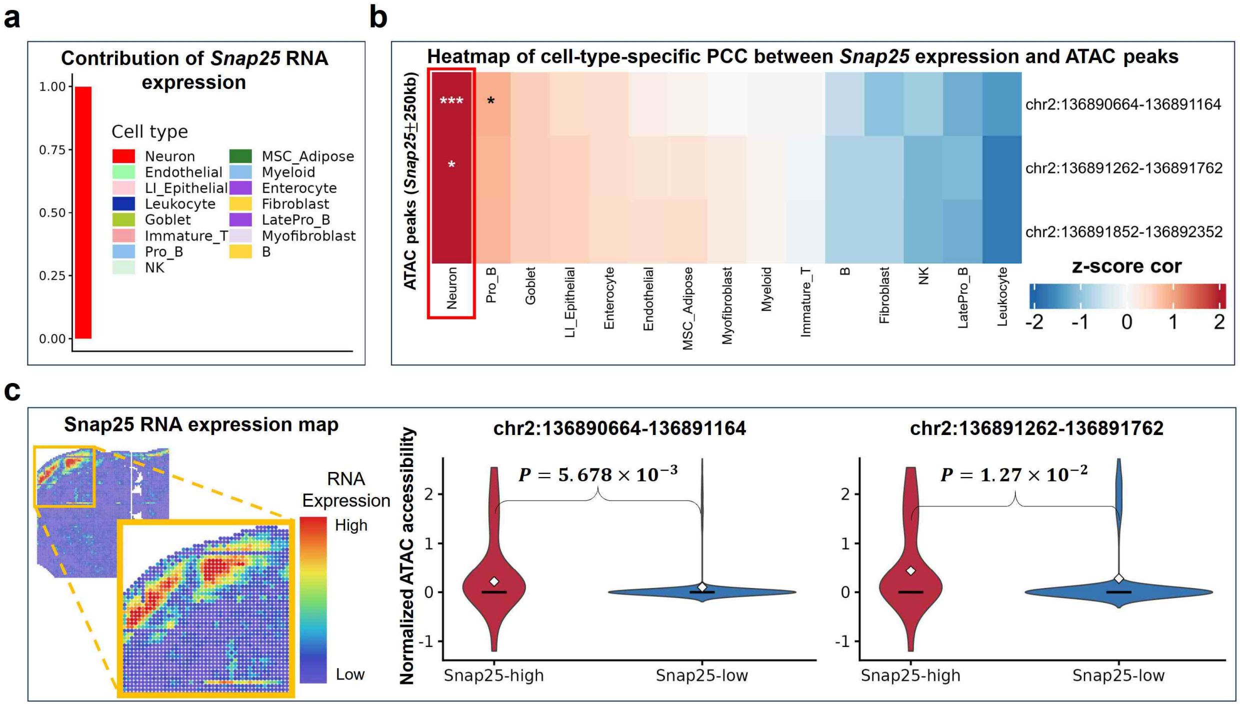

Supplementary Figure S29: **Cell-type-aware integration of spatial RNA–ATAC-seq data reveals regulatory mechanisms underlying E13 embryonic brain development.** **a**, Decomposition of spatial *Snap25* expression into cell-type-specific contributions using SMODER-inferred cell-type composition, showing dominant expression in neural populations. **b**, Proportion-based association analysis linking chromatin accessibility at distal regulatory elements to *Snap25* expression at cell-type resolution. Each row represents an ATAC peak within ±250kb of the *Snap25* transcription start site, and each column corresponds to a major cell type. Colors indicate z-scored Spearman correlation coefficients between peak accessibility and cell-type-specific *Snap25* expression (red, positive; blue, negative). Statistical significance is indicated by * ($0.01 < p \leq 0.05$), ** ($0.001 < p \leq 0.01$) or *** ($p \leq 0.001$). **c**, Comparison of chromatin accessibility at *Snap25*-associated distal regulatory elements between *Snap25*-high and *Snap25*-low spatial regions, revealing selective enhancer accessibility in transcriptionally active domains.

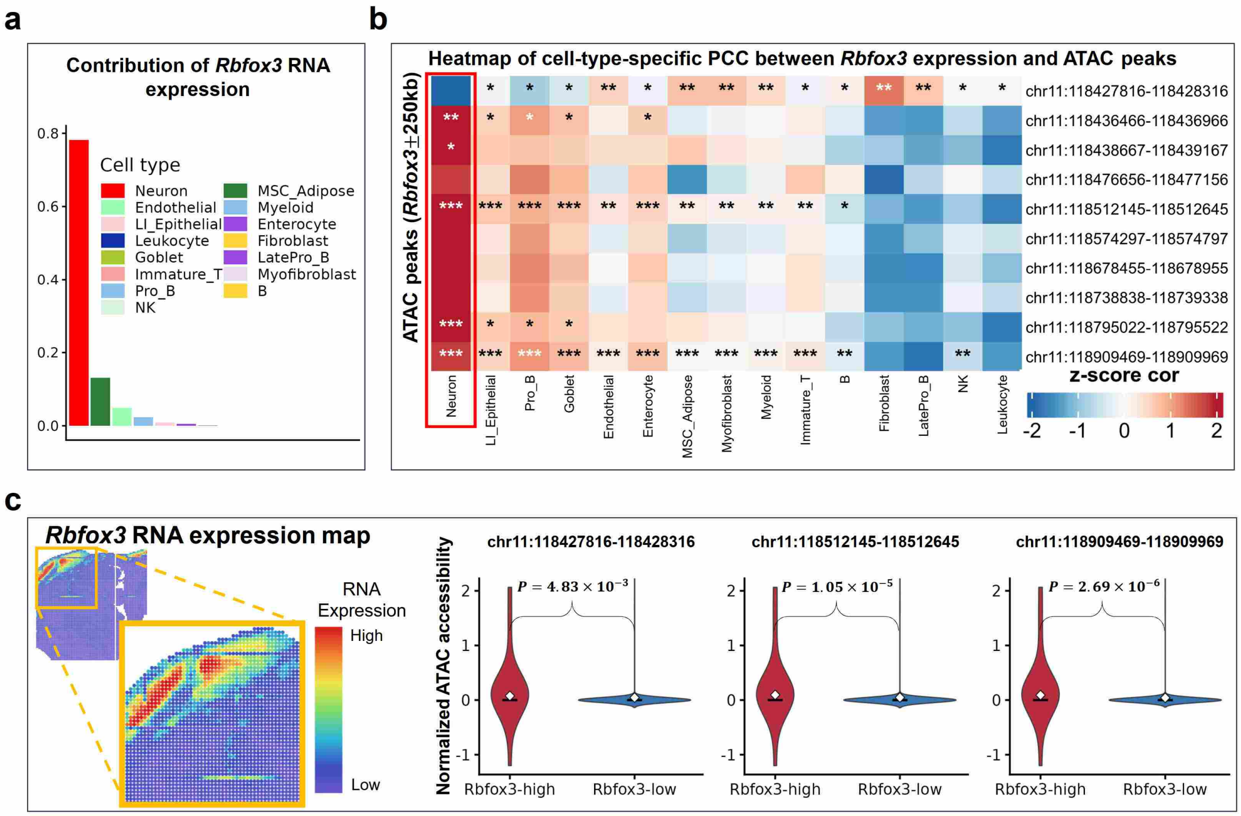

Supplementary Figure S30: **Cell-type-aware integration of spatial RNA–ATAC-seq data reveals regulatory mechanisms underlying E13 embryonic brain development.** **a**, Decomposition of spatial *Rbfox3* expression into cell-type-specific contributions using SMODER-inferred cell-type composition, showing dominant expression in neural populations. **b**, Proportion-based association analysis linking chromatin accessibility at distal regulatory elements to *Rbfox3* expression at cell-type resolution. Each row represents an ATAC peak within ±250kb of the *Rbfox3* transcription start site, and each column corresponds to a major cell type. Colors indicate z-scored Spearman correlation coefficients between peak accessibility and cell-type-specific *Rbfox3* expression (red, positive; blue, negative). Statistical significance is indicated by * ($0.01 < p \leq 0.05$), ** ($0.001 < p \leq 0.01$) or *** ($p \leq 0.001$). **c**, Comparison of chromatin accessibility at *Rbfox3*-associated distal regulatory elements between *Rbfox3*-high and *Rbfox3*-low spatial regions, revealing selective enhancer accessibility in transcriptionally active domains.

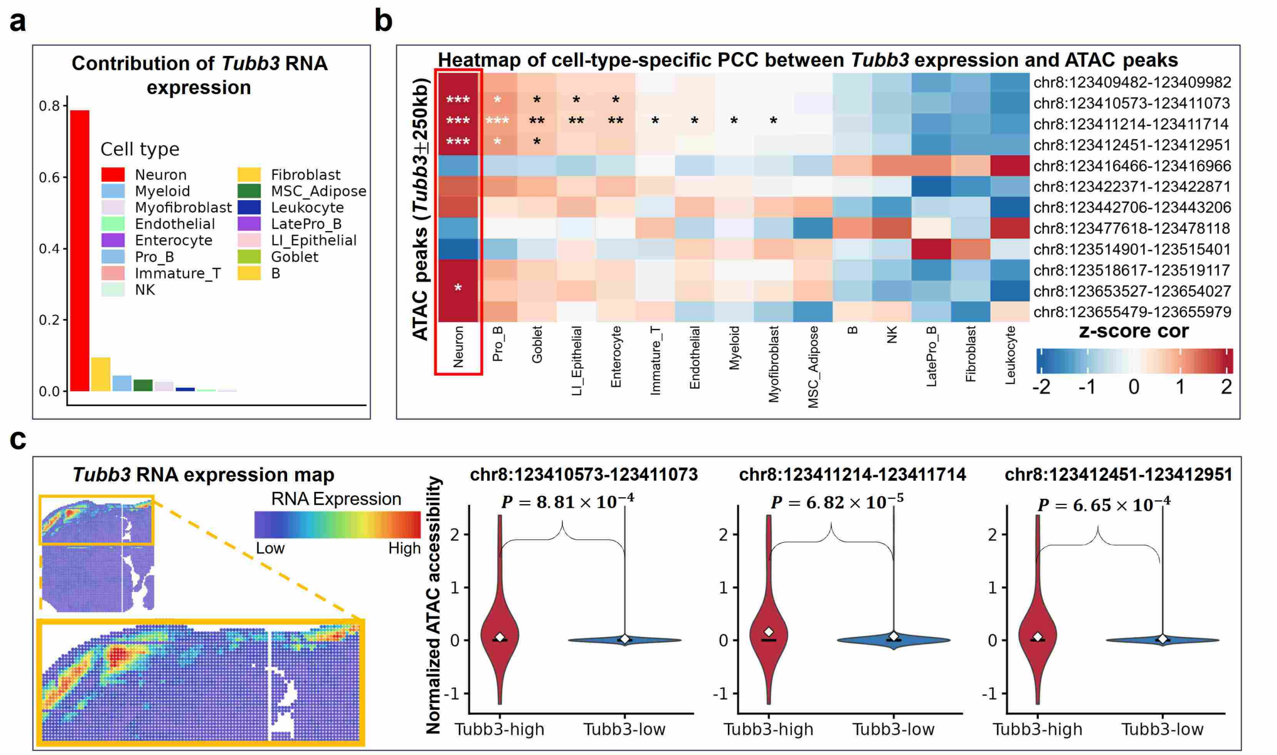

Supplementary Figure S31: **Cell-type-aware integration of spatial RNA–ATAC-seq data reveals regulatory mechanisms underlying E13 embryonic brain development.** **a**, Decomposition of spatial *Tubb3* expression into cell-type-specific contributions using SMODER-inferred cell-type composition, showing dominant expression in neural populations. **b**, Proportion-based association analysis linking chromatin accessibility at distal regulatory elements to *Tubb3* expression at cell-type resolution. Each row represents an ATAC peak within ±250kb of the *Tubb3* transcription start site, and each column corresponds to a major cell type. Colors indicate z-scored Spearman correlation coefficients between peak accessibility and cell-type-specific *Tubb3* expression (red, positive; blue, negative). Statistical significance is indicated by * ($0.01 < p \leq 0.05$), ** ($0.001 < p \leq 0.01$) or *** ($p \leq 0.001$). **c**, Comparison of chromatin accessibility at *Tubb3*-associated distal regulatory elements between *Tubb3*-high and *Tubb3*-low spatial regions, revealing selective enhancer accessibility in transcriptionally active domains.

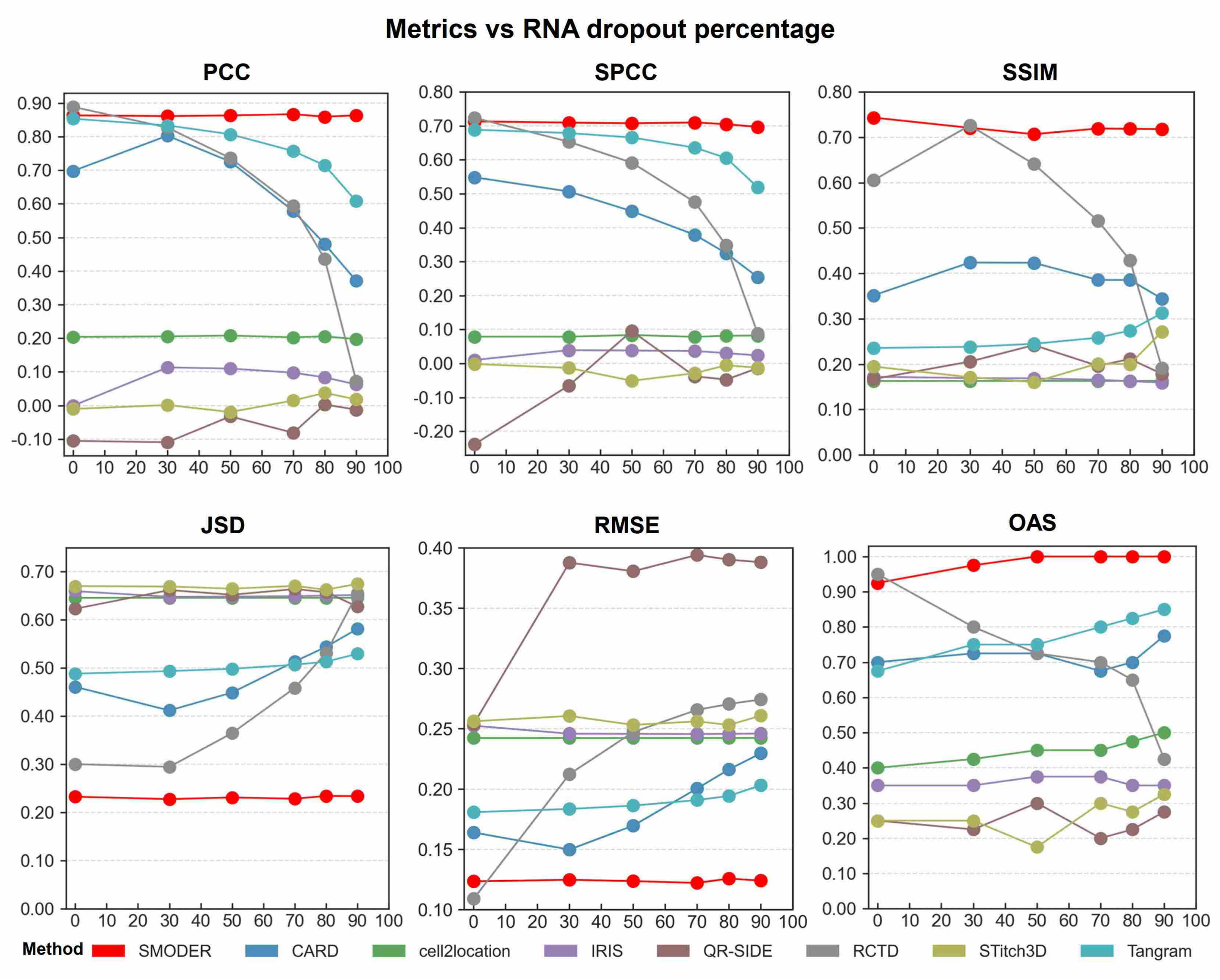

Supplementary Figure S32: **Quantitative analysis in generalization experiments with different dropout rates.** Line charts illustrate five metrics of deconvolution performance, including PCC, SPCC, SSIM, RMSE and JSD, under different dropout rates (from 0% to 90%) on spatial transcriptomic data. Meanwhile, it presents the overall average score (OAS) calculated based on the ranking of each metric.

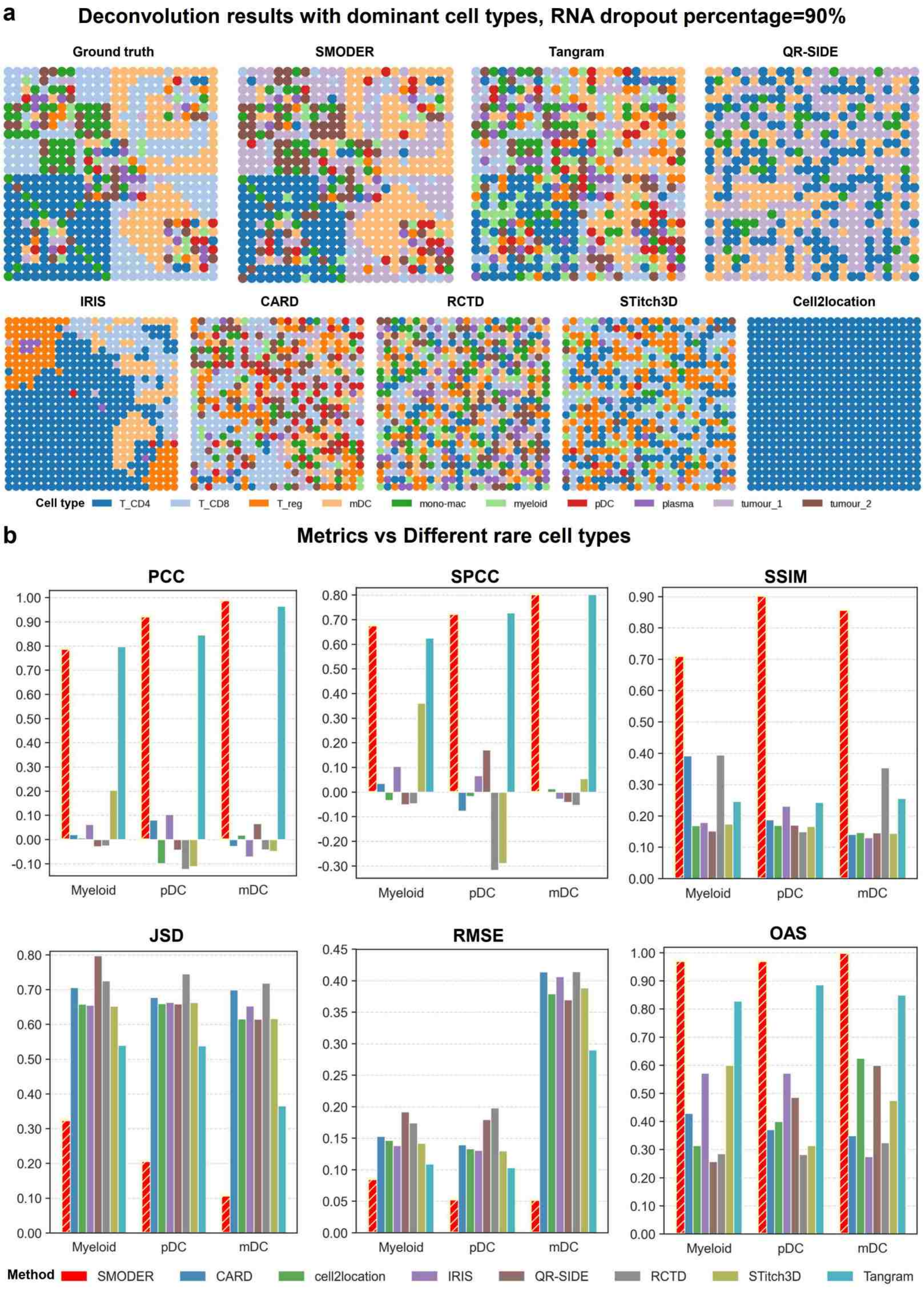

Supplementary Figure S33: **Quantitative analysis in generalization experiments. a**, Spatial heatmaps of dominant cell types inferred by SMODER and competing methods, alongside the ground truth, under a 90% RNA dropout rate. **b**, Bar plots summarizing deconvolution performance across five metrics (PCC, SPCC, SSIM, RMSE and JSD) for three rare cell types (Myeloid, 1.89%; pDC, 1.56%; mDC, 0.32%). Results are compared across SMODER and competing methods. The overall average score (OAS), computed based on metric rankings, is also shown.

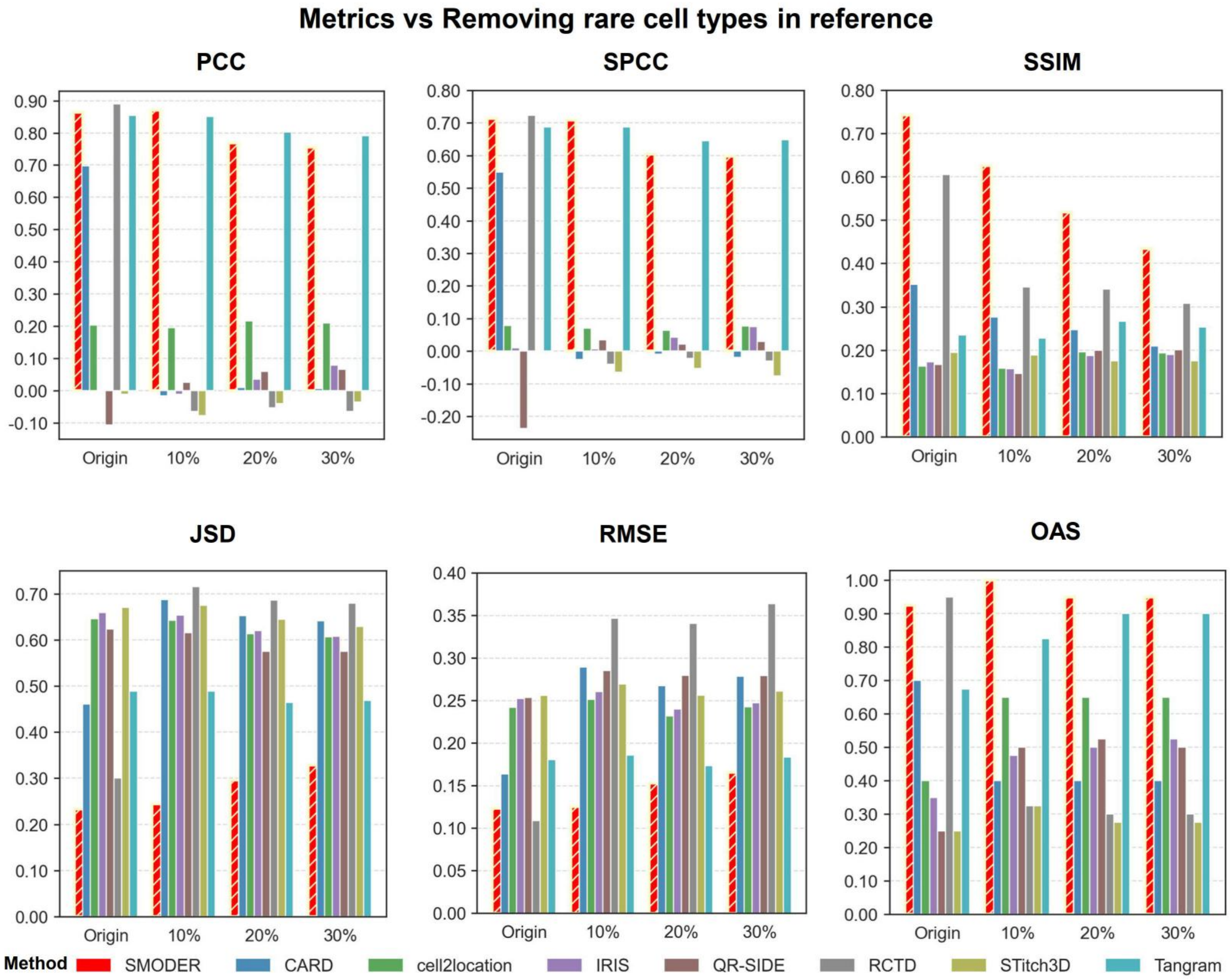

Supplementary Figure S34: **Quantitative analysis of deconvolution performance with various composition of cell types removed from reference data.** Bar plots show five deconvolution performance metrics (PCC, SPCC, SSIM, RMSE, JSD) and the corresponding OAS calculated from metric rankings, under simulated experiments with various cell-type removal ratios in reference data.

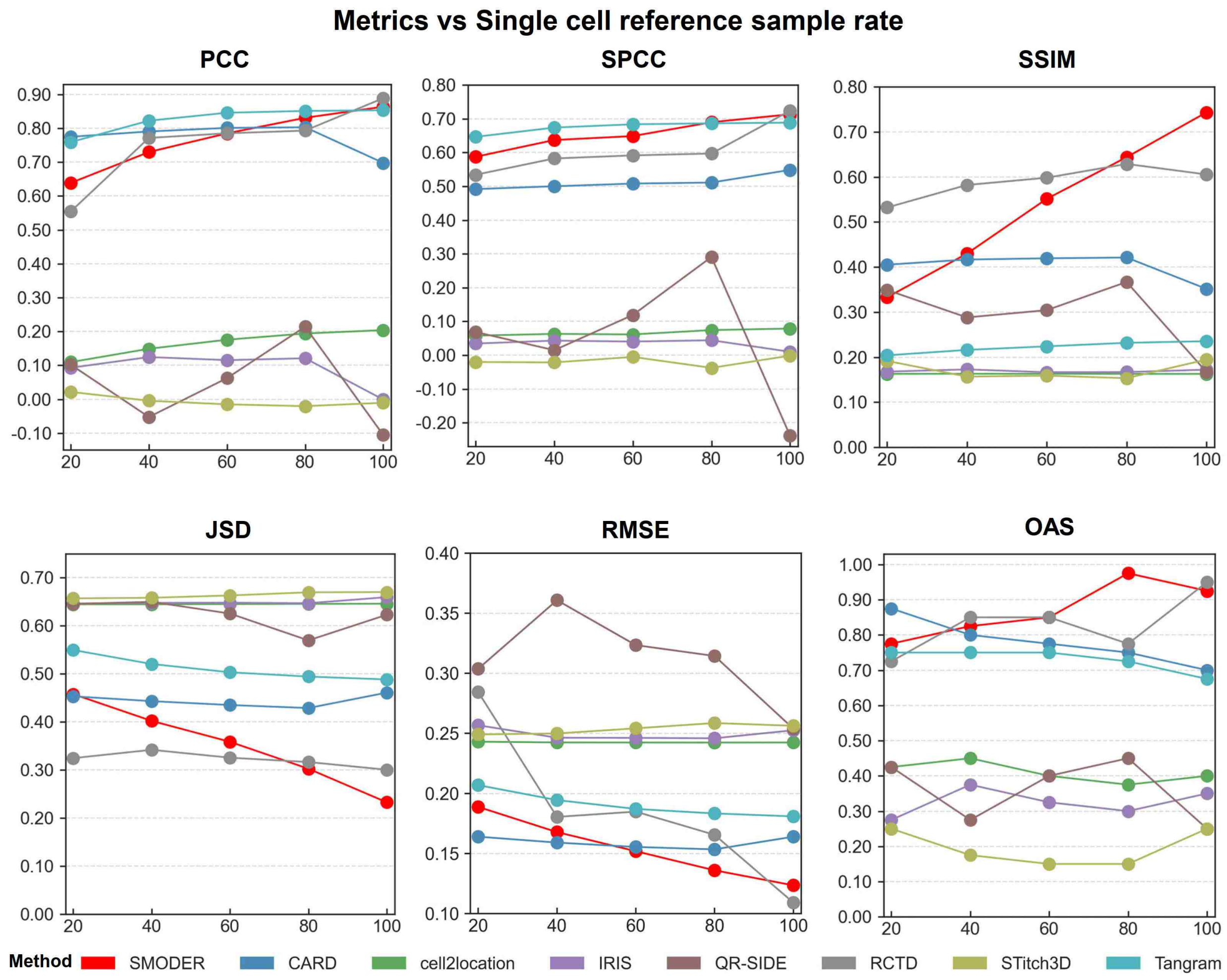

Supplementary Figure S35: **Quantitative analysis under different sampling rates in scRNA-seq reference data.** Line charts show five deconvolution performance metrics (PCC, SPCC, SSIM, RMSE, JSD) and the corresponding OAS calculated from metric rankings, under different sampling rates (from 20% to 100%) in scRNA-seq reference data.

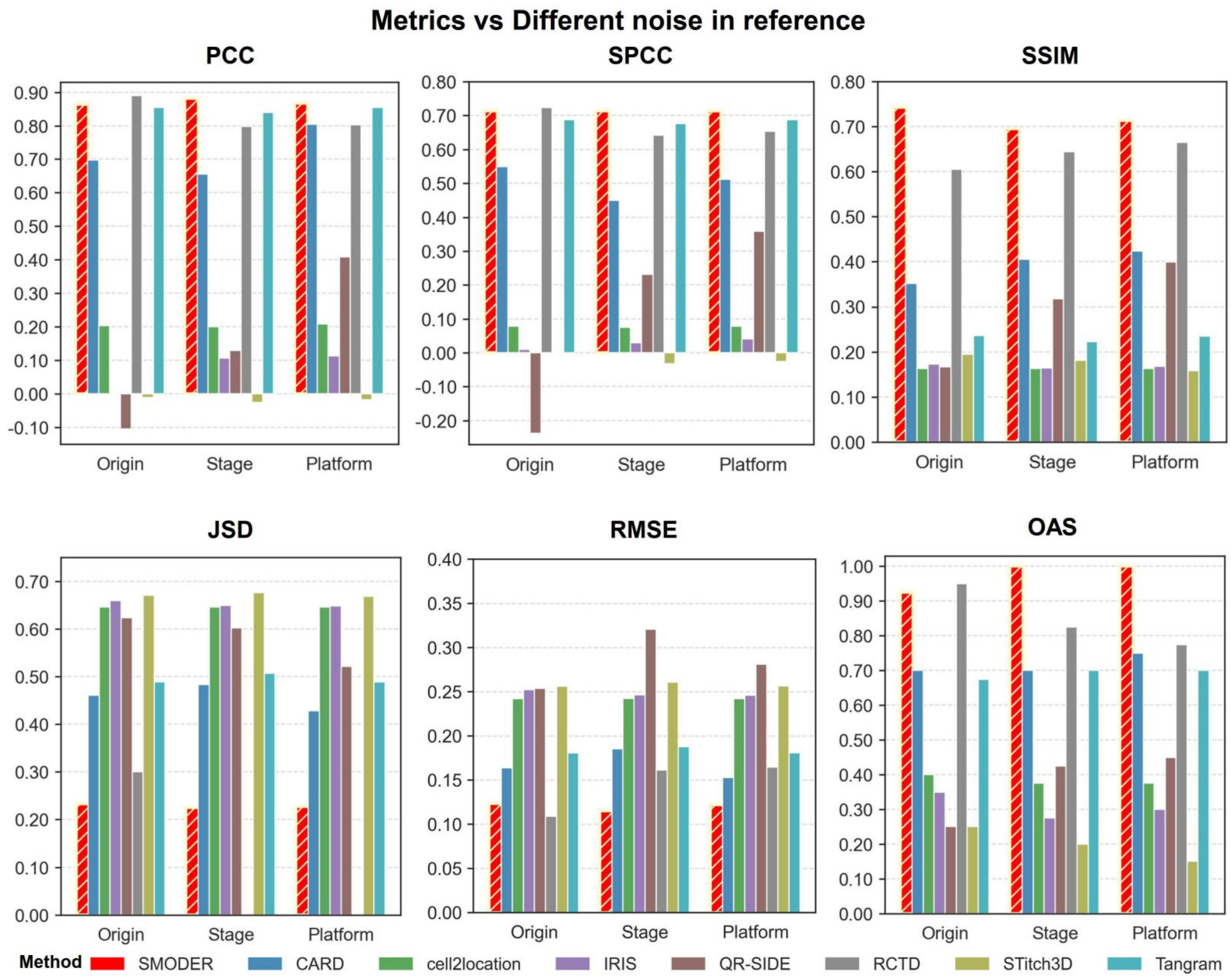

Supplementary Figure S36: **Quantitative analysis in generalization experiments under different types of noise imposed on reference data.** Bar plots illustrate the performance of SMODER and other competing methods under various noise conditions imposed on reference data, including noise from different development stages and different platforms. The performance was evaluated by five metrics (PCC, SPCC, SSIM, RMSE, JSD), and the overall average score (OAS) was calculated based on the ranking of each metric.

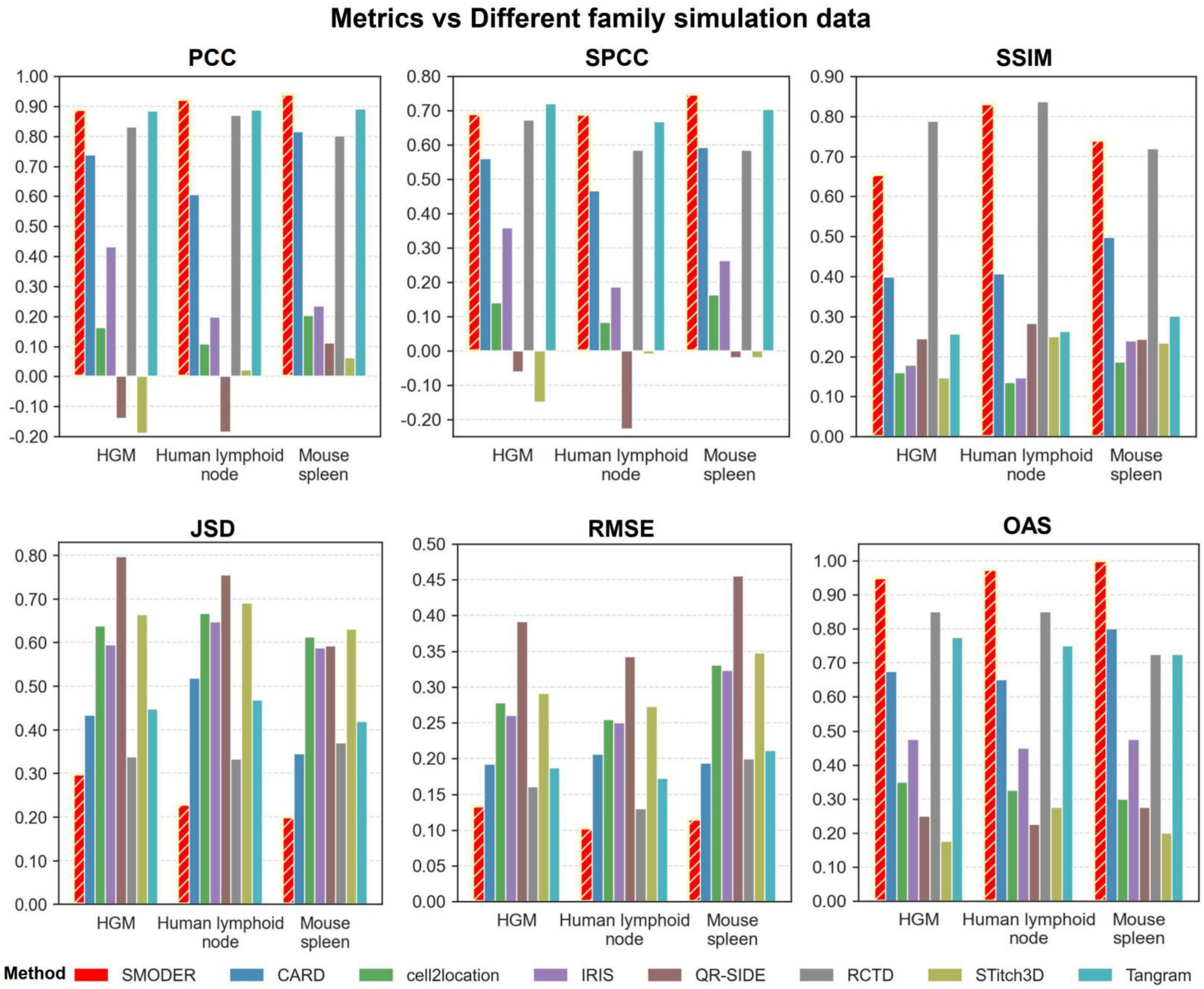

Supplementary Figure S37: **Quantitative analysis of generalization experiments under different datasets.** Bar plots illustrate the performance of SMODER and other competing methods across different datasets, including HGM, human lymphoid node and mouse spleen datasets. The performance was evaluated by five metrics (PCC, SPCC, SSIM, RMSE, JSD), and the OAS was calculated based on the ranking of each metric.

### Supplementary Notes

### 1 Methodological details of SMODER

This section provides a detailed description of the SMODER framework, including the probabilistic generative model, neural network parameterization, and training procedure. While the main text introduces the overall architecture, we present here the full formulation and implementation details to ensure reproducibility.

#### 1.1 Generative model and likelihood

To account for overdispersed count data in spatial transcriptomics, SMODER models RNA expression using a negative binomial (NB) distribution. For each spatial location $s$ and gene $g$,

$$X_{sg}^{\text{rna}} \sim \text{NB}(\mu_{sg}, \theta_g),$$

where $\mu_{sg}$ denotes the mean and $\theta_g$ is a gene-specific dispersion parameter. The mean parameter is linked to the inferred cell-type composition through

$$\mu_{sg} = l_s \exp\left\{\log\left(\sum_{c=1}^{C} P_{s,c} V_{cg}\right) + \alpha_s + \gamma_g\right\},$$

where $l_s$ is the library size, and $\alpha_s$ and $\gamma_g$ capture spot- and gene-specific effects.

The corresponding negative log-likelihood is given by

$$\mathcal{L}_{NB} = -\sum_{s=1}^{S}\sum_{g=1}^{G} \log P(X_{sg}^{\text{rna}} \mid \mu_{sg}, \theta_g),$$

where the negative binomial probability mass function is

$$P(X_{sg}^{\text{rna}} \mid \mu_{sg}, \theta_g) = \frac{\Gamma(X_{sg}^{\text{rna}} + \theta_g)}{\Gamma(\theta_g)\, X_{sg}^{\text{rna}}!} \left(\frac{\theta_g}{\theta_g + \mu_{sg}}\right)^{\theta_g} \left(\frac{\mu_{sg}}{\theta_g + \mu_{sg}}\right)^{X_{sg}^{\text{rna}}},$$

where $\Gamma(\cdot)$ is the gamma function.

#### 1.2 Neural network architecture

SMODER consists of modality-specific graph encoders, a cross-modal attention module, a fusion network, and task-specific decoders.

For each modality, we set $D = 256$ if the number of features in the original normalized expression matrix exceeds 256; otherwise, $D$ is set to the original feature dimension. The resulting input feature matrix $F \in \mathbb{R}^{S \times D}$ is then processed by two parallel graph convolutional encoders, constructed from the feature graph and the spatial graph, respectively. Each encoder follows a single-layer graph convolution

$$H = \sigma(\tilde{A}FW),$$

where $\tilde{A}$ is the normalized adjacency matrix, $W \in \mathbb{R}^{D\times 64}$ is a learnable weight matrix, and $\sigma(\cdot)$ denotes a nonlinear activation function (ReLU). This produces two modality-specific representations, which are then combined via an attention mechanism to yield unified embedding matrices $Z_{\text{rna}}$ and $Z_{\text{other}} \in \mathbb{R}^{S\times 50}$.

To integrate information across modalities, a bidirectional cross-attention mechanism is applied. Specifically, each modality embedding attends to the other through scaled dot-product attention, producing updated representations that capture cross-modal dependencies. The resulting embeddings are concatenated and passed through a fully connected layer to obtain a shared latent representation $Z \in \mathbb{R}^{S\times 50}$.

Cell-type composition are predicted from $Z$ using a two-layer fully connected network:

$$Z \to \text{ReLU}(ZW_1 + b_1) \to ZW_2 + b_2 \to \text{Softmax},$$

where $W_1 \in \mathbb{R}^{50\times 128}$ and $W_2 \in \mathbb{R}^{128\times C}$. The softmax layer ensures that the estimated composition are nonnegative and sum to one.

To preserve modality-specific information, two decoders are introduced to reconstruct the original feature matrices from the shared embedding. Each decoder is implemented as a two-layer fully connected network with hidden dimension 128 and ReLU activation:

$$Z \to \text{ReLU}(ZW_d^{(1)} + b_d^{(1)}) \to ZW_d^{(2)} + b_d^{(2)}.$$

These decoders map the latent representation back to the original feature space, enabling reconstruction-based regularization during training.

##### 1.3 Training objective, optimization, and regularization

The overall objective function combines four components:

$$\mathcal{L} = \mathfrak{L}_{NB} + \omega_{\text{recon}}\mathfrak{L}_{\text{recon}} + \omega_{\text{consist}}\mathfrak{L}_{\text{consist}} + \omega_{\text{spatial}}\mathfrak{L}_{\text{spatial}}.$$

The default weighting coefficients are set to $\omega_{\text{recon}} = 10^{-3}$, $\omega_{\text{consist}} = 1$, and $\omega_{\text{spatial}} = 10^{-5}$. To account for differences in feature distributions across datasets, the coefficients $(\omega_{\text{recon}}, \omega_{\text{consist}}, \omega_{\text{spatial}})$ are adjusted as follows: $(10^{-3}, 0.1, 10^{-5})$ for simulation data, $(10^{-3}, 1, 10^{-5})$ for mouse brain data, and $(10^{-3}, 1, 10^{-4})$ for both human breast cancer and mouse embryo datasets.

Model parameters are optimized using the Adam optimizer with a learning rate of $2 \times 10^{-3}$. Early stopping is employed based on the validation loss, with training terminated if no improvement is observed for 60 consecutive epochs. To enhance generalization and prevent overfitting, dropout with rate 0.1 is applied in the fusion module, and $\ell_2$ weight decay with coefficient $10^{-5}$ is incorporated into the optimization.

#### 2 Description on compared methods

**Benchmarking multi-modal methods on conception validation** To evaluate the impact of treating molecular modalities symmetrically in spatial deconvolution, we implemented two representative symmetric multi-modal baselines. The first baseline performs joint-feature deconvolution by enforcing a shared set of cell-type composition to simultaneously explain both spatial RNA measurements and a secondary molecular modality, using non-negative least squares (NNLS) regression (Algorithm 1). The second baseline comprises two methods, both of which adopt a joint-latent strategy commonly used in multimodal integration. The first employs nonnegative matrix factorization (NMF; Algorithm **??**) to learn a nonnegative shared latent representation, which is directly normalized across latent dimensions to obtain cell-type composition probabilities. The second uses SpatialGlue [1] to learn a shared latent representation, to which a softmax transformation is applied to obtain the corresponding cell-type composition probabilities.

Both baselines treat molecular modalities symmetrically. However, the NNLS-based baseline corresponds to an idealized scenario requiring paired reference profiles across modalities, which are unavailable in our setting. By contrast, the NMF/SpatialGlue-based approach does not explicitly incorporate reference cell-type guidance during factorization, and therefore latent components must subsequently be aligned with cell types in a post hoc manner.

---

**Algorithm 1** NNLS

---

**Input:** Spatial RNA matrix $X^{\text{RNA}} \in \mathbb{R}^{S\times G}$;
Secondary modality matrix $X^{\text{other}} \in \mathbb{R}^{S\times M}$;
Reference signatures $B^{\text{RNA}} \in \mathbb{R}^{G\times C}$ and $B^{\text{other}} \in \mathbb{R}^{M\times C}$;
Modality weight $\lambda = 1$.
**Output:** Estimated cell-type composition $\widehat{\Pi} \in \mathbb{R}^{S\times C}$.
Construct joint feature matrix:

$$X^{\text{joint}} = \left[X^{\text{RNA}}, \sqrt{\lambda}X^{\text{other}}\right]$$

Construct joint reference matrix:

$$B^{joint} = \begin{bmatrix} B^{\text{RNA}} \\ \sqrt{\lambda}B^{\text{other}} \end{bmatrix}$$

**for** $s = 1, \ldots, S$ **do**

  Solve non-negative least square regression:

$$\widehat{\pi}_s = \arg\min_{\pi\geq 0} \|X_s^{\text{joint}} - B^{\text{joint}}\pi\|_2^2$$

  Normalize $\widehat{\pi}_s$ to sum to one.

**return** $\widehat{\Pi}$

---

**Algorithm 2** NMF

**Input:** Spatial RNA matrix $X^{\text{RNA}} \in \mathbb{R}^{S\times G}$;
Secondary modality matrix $X^{\text{other}} \in \mathbb{R}^{S\times M}$;
Latent dimension $C$.
**Output:** Estimated composition $\widehat{\Pi} \in \mathbb{R}^{S\times C}$.
Concatenate modalities:

$$X^{joint} = [X^{\text{RNA}}, X^{\text{other}}]$$

Perform joint nonnegative matrix factorization:

$$X^{\text{joint}} \approx \Pi W^{\top},$$

subject to

$$\Pi \geq 0, \quad W \geq 0,$$

where

$$\Pi \in \mathbb{R}_{+}^{S\times C}, \quad W \in \mathbb{R}_{+}^{(G+M)\times C}.$$

Normalize rows of $\Pi$ to obtain cell-type composition:

$$\widehat{\pi}_s = \frac{\Pi_s}{\mathbf{1}^{\top}\Pi_i}.$$

**return** $\widehat{\Pi}$

**Spatial RNA-only methods** We compared SMODER with seven established spatial RNA-only deconvolution methods: RCTD [2], CARD [3], IRIS [4], QR-SIDE [5], Tangram [6], cell2location [7], STitch3D [8] and TACIT [9]. RCTD models spatial transcriptomics counts using a Poisson mixture framework, where spot-level expression is represented as a weighted combination of reference cell-type profiles, with platform effects explicitly accounted for. CARD adopts a conditional autoregressive (CAR) model to incorporate spatial smoothness, combining a regression-based deconvolution framework with spatial priors to improve robustness. IRIS employs a probabilistic model with iterative refinement, integrating reference and spatial data through likelihood-based inference to enhance cell-type proportion estimation. QR-SIDE is a qualitative-reference-based spatial deconvolution method that estimates cell-type composition by adaptively weighting marker genes. It further integrates spatial information through a Potts model, enabling joint inference of cell-type composition and spatial domains while promoting spatial continuity. Tangram uses a deep learning framework to map single-cell data onto spatial locations by optimizing the alignment between cell-type expression and spatial transcriptomics profiles. Cell2location is based on a hierarchical Bayesian model that leverages negative binomial likelihoods and variational inference to jointly model cell-type abundance and technical effects. STitch3D integrates spatial transcriptomics with single-cell data using a graph-based framework, reconstructing spatial organization by learning cross-modality correspondences in a three-dimensional latent space. TACIT is originally designed for cell-type annotation in single-cell-resolution spatial omics data. To adapt TACIT to spot-level deconvolution, we applied its signature-scoring strategy to each spatial spot. Specifically, for each spot, we calculated its relevance score for every cell type by comparing the spot-level signature-gene expression vector with the corresponding cell-type profile in the basis matrix. The resulting cell-type scores were then normalized across cell types to obtain a compositional vector summing to one, which was interpreted as the estimated cell-type proportions of that spot.

All methods were implemented using their official software packages, and executed following recommended guidelines whenever available. Specifically, Tangram (v1.0.4) was implemented in Python using default parameters without manual tuning. Its preprocessing pipeline was aligned with that of SMODER. STitch3D (v1.0.3) was also run in Python with default settings, and its preprocessing was kept consistent with that of Tangram. Cell2location (v0.1.5) was executed in Python with a learning rate of 0.005 and 1,000 training epochs, balancing computational efficiency and model convergence; its preprocessing was consistent with the aforementioned methods. RCTD was implemented using the `spacexr` R package (v2.2.1) with default parameters. CARD was implemented using the `CARD` R package (v1.1) following the authors' recommended settings. QR-SIDE was implemented using the `QRSIDE` R package (v1.0.0), and IRIS was implemented using the `IRIS` R package (v1.0), both with default parameters. TACIT was excuted in `TACIT` R package (0.0.0.1).

### 3 Details on evaluations in both simulations and real data sets

#### 3.1 Evaluations in simulations

In simulation studies, the ground-truth cell-type composition matrix $P^* = (P^*_{s,c}) \in \mathbb{R}^{S\times C}$ is available, where $S$ denotes the number of spatial spots and $C$ denotes the number of cell types. Let $P = (P_{s,c})$ denote the estimated composition matrix. We evaluated deconvolution accuracy using five complementary metrics: root mean square error (RMSE), Pearson correlation coefficient (PCC), Spearman correlation coefficient (SPCC), Jensen–Shannon divergence (JSD), and structural similarity index (SSIM).

**Root mean square error (RMSE).** RMSE measures the average magnitude of estimation errors between the estimated and true cell-type composition:

$$\mathrm{RMSE} = \sqrt{\frac{1}{SC}\sum_{s=1}^{S}\sum_{c=1}^{C}\left(P_{s,c} - P^*_{s,c}\right)^2}.$$

**Pearson correlation coefficient (PCC).** PCC evaluates the linear association between estimated and true cell-type proportions. Let $\mathrm{vec}(\cdot)$ denote the vectorization operator. PCC is computed as

$$\mathrm{PCC} = \frac{\mathrm{cov}(\mathrm{vec}(P), \mathrm{vec}(P^*))}{\sigma_{\mathrm{vec}(P)}\sigma_{\mathrm{vec}(P^*)}},$$

where $\mathrm{cov}(\cdot,\cdot)$ denotes covariance and $\sigma_{\mathrm{vec}(\cdot)}$ denotes the standard deviation.

**Spearman correlation coefficient (SPCC).** SPCC measures the rank-based correlation between estimated and true proportions, reflecting robustness to scale differences. Let rank$(\cdot)$ denote the rank transformation. SPCC is defined as

$$\text{SPCC} = \text{corr}\left(\text{rank}(\text{vec}(P)), \text{rank}(\text{vec}(P^*))\right),$$

where corr$(\cdot,\cdot)$ represents the Pearson correlation coefficient.

**Jensen–Shannon divergence (JSD).** To measure similarity between the distributions of estimated and true composition, we computed the Jensen–Shannon divergence. Let $p$ and $q$ denote the normalized composition distributions derived from vec$(P)$ and vec$(P^*)$, respectively. JSD is defined as

$$\text{JSD}(p\|q) = \frac{1}{2}D_{\text{KL}}(p\|m) + \frac{1}{2}D_{\text{KL}}(q\|m),$$

where $m = \frac{1}{2}(p+q)$ and $D_{\text{KL}}(\cdot\|\cdot)$ denotes the Kullback–Leibler divergence: $D_{\text{KL}}(p\|q) = \sum_i p_i \log \frac{p_i}{q_i}$. JSD is symmetric and bounded between 0 and log2, with smaller values indicating closer agreement between the two distributions.

**Structural similarity index (SSIM).** To evaluate the preservation of spatial structure in the proportion maps, we employed the structural similarity index. For each cell type $c$, SSIM compares the spatial maps $\mathbf{p}_c = (P_{1,c}, \ldots, P_{S,c})$ and $\mathbf{p}_c^* = (P_{1,c}^*, \ldots, P_{S,c}^*)$. SSIM is defined as

$$\text{SSIM}(\mathbf{p}_c, \mathbf{p}_c^*) = \frac{(2\mu_c\mu_c^* + \epsilon_1)(2\sigma_{cc^*} + \epsilon_2)}{(\mu_c^2 + (\mu_c^*)^2 + \epsilon_1)(\sigma_c^2 + (\sigma_c^*)^2 + \epsilon_2)},$$

where $\mu_c$ and $\mu_c^*$ denote the means of the two maps, $\sigma_c^2$ and $(\sigma_c^*)^2$ denote their variances, and $\sigma_{cc^*}$ denotes the covariance between them. Constants $\epsilon_1$ and $\epsilon_2$ are small positive values used for numerical stability. The final SSIM score is obtained by averaging over all cell types:

$$\text{SSIM} = \frac{1}{C}\sum_{c=1}^{C}\text{SSIM}(\mathbf{p}_c, \mathbf{p}_c^*).$$

Higher SSIM values indicate better recovery of spatial patterns.

#### 3.2 Evaluations in real data sets

In real data applications where the underlying cell-type composition matrix is unavailable, we use eight complementary metrics that characterize local mixing, domain separation, spatial continuity, boundary sharpness, and global concordance with annotated spatial domains, including local inverse Simpson's index, silhouette coefficient, percentage of abnormal spots, and chaos score, spatial map similarity, balanced edge agreement ratio. In addition, adjusted Rand index and normalized mutual information computed between dominated cell-type assignments and manual annotations assess the global consistency of inferred spatial structure.

**Local inverse Simpson's index** The Local Inverse Simpson's Index (LISI) quantifies how well shared cell populations are merged across a tissue section by assigning each spot a diversity score that reflects the effective number of spatial domains present in its local neighborhood. In real data, accurate deconvolution should preserve LISI values close to 1, indicating that each spot's neighborhood is dominated by a single spatial domain, as defined by the inferred proportion matrix. In contrast, an erroneous proportion matrix produces neighborhoods with LISI values greater than 1, with the worst cases approaching the total number of spatial domains, suggesting that neighboring spots belong to entirely different domains.

**Silhouette coefficient score** We evaluated the quality of the inferred composition matrix using the silhouette coefficient (SC) based on manually annotated domain labels. Specifically, each spatial location is represented by its inferred cell-type composition vector, and the silhouette coefficient is computed with respect to the domain assignments in $\mathcal{C}$.

For a given spot $i$, let $a(i)$ denote the average distance between $i$ and all other spots within the same domain, and $b(i)$ the minimum average distance between $i$ and spots in other domains. The silhouette coefficient for spot $i$ is defined as

$$s(i) = \frac{b(i) - a(i)}{\max\{a(i),\, b(i)\}}.$$

The final SC score is obtained by averaging $s(i)$ over all samples. A higher SC score indicates that samples within the same annotated domain are more similar to each other and better separated from other domains, suggesting that the inferred composition matrix more effectively preserves domain-specific biological structure.

**CHAOS score** The CHAOS score [10] quantifies the spatial continuity of detected domains based on the inferred composition matrix. Specifically, each spot is represented by its cell-type composition vector, and neighborhood relationships are defined in this composition space.

For each domain $k \in \{1, \ldots, K\}$, we construct a one-nearest-neighbor (1NN) graph by connecting each spot to its nearest neighbor (within the same domain) according to Euclidean distance in the composition space. Let $w_{ij}$ denote the Euclidean distance between spot $i$ and its nearest neighbor $j$. The CHAOS score is then defined as the average edge length across all such 1NN graphs:

$$\text{CHAOS} = \frac{1}{S} \sum_{k=1}^{K} \sum_{i \in \mathcal{C}_k} w_{i,\text{NN}(i)},$$

where $S$ is the total number of spots, $K$ is the number of domains, $\mathcal{C}_k$ denotes the set of spots in domain $k$, and $\text{NN}(i)$ is the nearest neighbor of spot $i$ within $\mathcal{C}_k$. A smaller CHAOS score indicates higher spatial continuity, as spots within the same domain are more similar in the inferred proportion space.

**Percentage of abnormal spots** The percentage of abnormal spots (PAS) score measures the randomness of the spots that are located outside of the spatial domain where it was clustered to. For each spot, we use the proportion matrix to define its neighbors. The PAS score is calculated as the ratio of spots with a cluster label that is different from at least six of its neighboring ten spots. A small PAS score indicates spot homogeneity within spatial domains.

**Spatial map similarity** Spatial map similarity (SMS) is a metric we define to quantify the agreement between the spatial distributions of inferred cell composition and annotated reference domains. For each predicted and annotated cell type, spatially smoothed maps are constructed by aggregating information from neighboring locations using a Gaussian kernel, thereby capturing regional enrichment patterns and reducing spot-level noise. Similarity between predicted and annotated spatial maps is quantified using cosine similarity after normalization, which assesses the concordance of spatial patterns independently of scale.

Since the identities and numbers of inferred cell types may not directly correspond to annotated labels, SMS employs an optimal one-to-one matching between predicted and annotated types using the Hungarian algorithm [11]. The final SMS score is defined as the average similarity across matched pairs. Higher SMS values indicate better recovery of annotated spatial domains and their spatial organization by the inferred cell-type composition.

**Balanced edge agreement ratio** We define the balanced edge agreement ratio (BEAR) to quantify the consistency between inferred dominant cell types and annotated domain labels along edges of a spatial $k$-nearest-neighbor graph, while correcting for class imbalance among edges. Specifically, edges are divided into two groups: (i) edges connecting samples with the same annotation in $\mathcal{C}$, and (ii) edges connecting samples with different annotations.

BEAR evaluates performance by combining two complementary criteria: the agreement rate on same-label edges (i.e., the proportion of edges where inferred labels in $\mathcal{K}$ are also identical) and the disagreement rate on different-label edges (i.e., the proportion of edges where inferred labels differ). The final score is defined as the average of these two quantities, assigning equal weight to within-domain continuity and between-domain separation.

This formulation corresponds to an edge-level balanced accuracy and mitigates bias caused by the predominance of same-label edges in spatial data [12]. In particular, a degenerate solution that assigns all samples to a single cell type yields a baseline score of 0.5, ensuring that higher values reflect meaningful recovery of spatial boundaries and local domain structure rather than class imbalance.

**Adjusted Rand index** The adjusted Rand index (ARI) quantifies the agreement between ground truth labels and predicted cluster assignments based on pairwise consistency. Let $\mathcal{C}$ denote the annotated spatial domains and $\mathcal{K}$ the inferred dominated cell types. Among all $\binom{S}{2}$ pairs of samples, let $a$ be the number of pairs assigned to the same cluster in both $\mathcal{C}$ and $\mathcal{K}$, and $b$ the number of pairs assigned to different clusters in both $\mathcal{C}$ and $\mathcal{K}$. The (unadjusted) Rand index (RI) is defined as

$$RI = \frac{a+b}{\binom{S}{2}}.$$

However, RI does not account for chance agreement and may yield inflated values under random label assignments. To correct for this effect, the ARI adjusts RI by its expected value under a random permutation model:

$$\text{ARI} = \frac{RI - \mathbb{E}[RI]}{\max(RI) - \mathbb{E}[RI]}.$$

This normalization ensures that ARI is close to 0 for random partitions and equals 1 for perfect agreement.

**Normalized mutual information** The normalized mutual information (NMI) measures the agreement between $\mathcal{C}$ and $\mathcal{K}$ from an information-theoretic perspective. The entropy of $\mathcal{C}$ is defined as

$$H(\mathcal{C}) = -\sum_{i=1}^{|\mathcal{C}|} P(i)\text{log}P(i),$$

where $P(i) = \frac{|\mathcal{C}_i|}{S}$ is the probability that a randomly selected sample belongs to domain $\mathcal{C}_i$. Similarly, the entropy of $\mathcal{K}$ is

$$H(\mathcal{K}) = -\sum_{j=1}^{|\mathcal{K}|} P(j)\text{log}P(j),$$

where $P(j) = \frac{|\mathcal{K}_j|}{S}$.

The mutual information between $\mathcal{C}$ and $\mathcal{K}$ is defined as

$$\text{MI}(\mathcal{C}, \mathcal{K}) = \sum_{i=1}^{|\mathcal{C}|}\sum_{j=1}^{|\mathcal{K}|} P(i,j)\text{log}\frac{P(i,j)}{P(i)P(j)},$$

where $P(i,j) = \frac{|\mathcal{C}_i \cap \mathcal{K}_j|}{S}$ denotes the joint probability.

To facilitate comparison, the mutual information is normalized by the entropies:

$$\text{NMI}(\mathcal{C}, \mathcal{K}) = \frac{\text{MI}(\mathcal{C}, \mathcal{K})}{\sqrt{H(\mathcal{C})H(\mathcal{K})}}.$$

The resulting score ranges from 0 to 1, with higher values indicating stronger agreement between $\mathcal{C}$ and $\mathcal{K}$.

**Boundary-interior ratio** The boundary-interior ratio (BIR) serves as a refined quantitative indicator to measure the relative difference between boundary discrimination clarity and interior regional homogeneity in tissue architecture. For each spot $s_i$, we identify its 8 neighboring spots using the $K$-nearest neighbor (KNN) algorithm to construct its local neighborhood $\mathcal{N}(s_i)$. A deterministic rule is applied: if more than 2 spots within the neighborhood exhibit annotations distinct from that of $s_i$, $s_i$ is classified as a boundary spot; otherwise, it is regarded as an internal spot. Assuming a total of $\mathcal{K}$ cell types, a $\mathcal{K}$-dimensional neighborhood count vector $\boldsymbol{c}_i = [count_{i1}, count_{i2}, \ldots, count_{i\mathcal{K}}]$ is constructed for each spot, where $count_{ij}$ denotes the number of spots in $\mathcal{N}(s_i)$ for which the $j$-th cell type is the dominant one. The Euclidean distance is adopted to quantify the distribution divergence between the count vector of the central spot and its neighbors. The discrepancy score for a single spot $d_i$ is defined as the mean Euclidean distance between the spot and all its neighbors, formulated as: $d_i = \frac{1}{\mathcal{K}} \sum_{j \in \mathcal{N}(i)} \|\boldsymbol{c}_i - \boldsymbol{c}_j\|_2$ where $\mathcal{N}(i)$ denotes the set of $K$-nearest neighbors of spot $i$. A higher $d_i$ indicates stronger spatial heterogeneity in the cellular composition of the spot.

Based on the classified boundary and internal spots, the average heterogeneity scores of all boundary spots $D_{\text{boundary}}$ and internal spots $D_{\text{interior}}$ are calculated as follows:

$$D_{\text{boundary}} = \frac{1}{N_{\text{bound}}} \sum_{i \in \Omega_{\text{bound}}} d_i, \quad D_{\text{interior}} = \frac{1}{N_{\text{inter}}} \sum_{i \in \Omega_{\text{inter}}} d_i,$$

where $\Omega_b$ and $\Omega_i$ represent the sets of boundary and internal spots, respectively, and $N_{bound}$ and $N_{\text{inter}}$ denote the corresponding numbers of spots. Ultimately, the boundary-interior ratio is defined as the ratio between the average heterogeneity of boundary spots and that of internal spots:

$$\text{BIR} = \frac{D_{\text{boundary}}}{D_{\text{interior}}},$$

Consequently, a higher value of this indicator signifies sharper regional differentiation and more distinct demarcation between tissue functional domains.

### 4 Simulation design

We designed a comprehensive suite of simulation experiments to evaluate the performance of SMODER under diverse conditions, including:

(1) concept validation simulations comparing SMODER with symmetric multi-modal deconvolution methods;

(2) reference-matched and reference perturbation simulations including cell-type removal, misannotation, and addition of artificial cell types;

(3) technology-aware simulations mimicking different spatial multi-omics measurement settings;

(4) robustness and generalization simulations under external reference datasets.

#### 4.1 Concept validation simulation

**Reference data sets** In this simulation experiments, we adopted the following two reference data.
Human melanoma RNA and epigenomic sequencing data For proof-of-concept analysis, a subsampled human melanoma dataset with jointly profiled RNA and epigenomic measurements [13] was used as the reference. The dataset comprises 210 cells across five common cell types, including tumor cells 2 (tumour_2; $n = 80$), tumor cells 1 (tumour_1; $n = 60$), CD4$^+$ T cells (T_CD4; $n = 40$), CD8$^+$ T cells (T_CD8; $n = 20$), and plasma cells ($n = 10$).
Human brain RNA-ADT sequencing data For the dataset of human brain (RNA and protein sequencing) [14], we obtained expression measurements for a total of 2,515 cells from 10 common cell types. These cell types include HSPC (n=500), CD14 Mono (n=450), CD8 TEM (n=300), NK (n=200), B intermediate (n=120), CD4 TEM (n=60), cDC2 (n=50), B naive (n=24), Treg (n=18), and CD8 Naive (n=15).

**Data generation process** We generate synthetic spatial multi-omics data using the above single-cell multi-omics reference data. The generated spatial multi-omics dataset consists of a $24 \times 24$ square spot lattice, within which we delineate 8 spatially structured regions. We predefine the cell-type composition for each region, where the probability of sampling a given cell type is determined by its proportional abundance in the original single-cell data. To better mimic real biological data, we set the average number of cells per spot to 10. The gene expression level at each spatial spot is then computed as the summation of expression values across all sampled cells for the corresponding gene. To recapitulate the noise and technical artifacts inherent in real spatial transcriptomic data, we introduce additional noise and attenuate highly variable genes (HVGs) for the RNA modality. First, we introduce Poisson noise with a baseline parameter $\lambda = 15$. To simulate spatial technical variations, we dynamically adjust the noise intensity for each spot based on its relative position to the center of the tissue region. Specifically, the spatial center $(\bar{x}, \bar{y})$ is calculated as the centroid of all spot coordinates, i.e., $\bar{x} = \frac{1}{N}\sum_{s=1}^{N} x_s$ and $\bar{y} = \frac{1}{N}\sum_{s=1}^{N} y_s$. The Euclidean distance from the $s$-th spot to the center is computed as: $d_s = \sqrt{(x_s - \bar{x})^2 + (y_s - \bar{y})^2}$. Let $d_{\max}$ denote the maximum distance from the center to the outermost boundary spots. The spatially weighted noise parameter $\lambda_s$ for each spot $s$ is defined as:

$$\lambda_s = \left(\frac{d_s}{d_{\max}} \times 2 + 1\right) \times \lambda.$$

Subsequently, we perform dilution of highly variable genes: for each cell type, the expression values of the top 200 highly variable genes are downscaled to 20% (0.2×) of their original magnitude. This completes the generation of the simulated spatial multi-omics data.

Non-negative matrix factorization (NMF) and non-negative least squares(NNLS) regression using complete multi-modal single-cell data are conventional methods for symmetric multi-modal deconvolution. In this experiment, we design a systematic simulation to validate the superiority of SMODER over these general symmetric multi-modal deconvolution approaches.

#### 4.2 Reference-matched and perturbation experiments

**Reference data sets** In this set of simulation experiments, we used the same reference data as in the concept validation simulations, except that we subsampled 2,515 cells from the human melanoma dataset with jointly profiled RNA and epigenomic measurements. The resulting dataset comprised 2,515 cells across 10 cell types, including CD8$^{+}$ T cells ($n$ = 1,020), tumor cells (subtype 1; $n = 561$), monocyte–macrophages ($n = 307$), tumor cells (subtype 2; $n = 272$), plasma cells ($n = 166$), regulatory T cells (Treg; $n = 72$), CD4$^{+}$ T cells ($n = 67$), myeloid cells ($n = 24$), myeloid dendritic cells (mDC; $n = 20$), and plasmacytoid dendritic cells (pDC; $n = 6$).

**Data generation process** To simulate spatial heterogeneity and extreme cell-type composition observed in real biological data, we further extend the data generation process of the concept validation simulation. Specifically, we define five non-overlapping $7 \times 7$ spatial regions, in which we disrupt the original cell-type sampling framework. For each targeted spot, the total cell count was randomly set between 10 and 20. A randomly selected dominant cell type occupied 80% of the total cells, while the rest 20% were randomly sampled from other cell types in the single-cell dataset. Spatial spots within these regions were generated accordingly. This procedure completed the construction of the simulated spatial multi-omics dataset. We analyzed the simulated spatial multi-omics data using SMODER alongside RNA-only deconvolution methods. In addition, we used these simulated datasets to assess the robustness of SMODER to hyperparameter choices.

In the reference-matched setting, we used the reference dataset with the same cell types as those in the simulated spatial multi-omics data. Furthermore, we constructed additional scenarios by perturbing the cell-type composition in the reference data. The specific scenarios are as follows:

Scenario 1: One cell type was removed from the original reference data (plasma was removed from the human melanoma RNA and epigenomic sequencing data, and CD4 TEM was removed from the human brain RNA and protein sequencing data), and the processed data were used for deconvolution of simulated data.

Scenario 2: A new synthetic cell type was added to the original reference data (mono-mac and tumour_2 were synthesized for the human melanoma RNA and epigenomic sequencing data, and CD14 Mono and CD8 TEM were synthesized for the human brain RNA and protein sequencing data), and the enriched data were used for deconvolution of simulated data.

Scenario 3: Certain cell types (plasma in the human melanoma RNA and epigenomic sequencing data, and CD4 TEM in the human brain RNA and protein sequencing data) were misannotated as cell_type_1, and the misannotated reference data were used for deconvolution of simulated data.

#### 4.3 Technology-aware simulations

**Reference data used** In this simulation experiments, we used another two reference data.
Mouse brain scRNA-seq data The original dataset contained 11 annotated cell types [7]. In total, 3,215 cells were subsampled across these cell types, including oligodendrocytes/oligodendrocyte precursor cells (Oligo/OPC; $n = 1000$), cortical excitatory neurons (Exc-Ctx; $n = 600$), non-cortical excitatory neurons (Exc-NonCtx; $n = 800$), inhibitory neurons (general subclass; Inh-General; $n = 400$), inhibitory neurons (molecular marker subtype; Inh-Marker; $n = 100$), and inhibitory neurons (Meis2; Inh-Meis2; $n = 50$), astrocytes (Astro; $n = 200$), neuroblasts (Nb; $n = 15$), endothelial cells (Endo; $n = 10$), microglia (Micro; $n = 20$), and a small group of unassigned or low-quality cells (Unknown; $n = 20$).
Human breast cancer scRNA-seq data ScRNA-seq data from human breast cancer [15] were used for analysis. In total, 3,080 subsampled cells were annotated into nine cell types, including T cells ($n = 1000$), cancer epithelial cells ($n = 900$), myeloid cells ($n = 500$), endothelial cells ($n = 400$), cancer-associated fibroblasts (CAFs; $n = 150$), perivascular-like cells (PVL; $n = 70$), normal epithelial cells ($n = 24$), plasmablasts ($n = 20$), and B cells ($n = 16$).

**Data generation process** We first generated the complementary omics modality by projecting single-cell RNA expression into a synthetic secondary feature space. Specifically, we constructed a sparse projection matrix $W \in \mathbb{R}^{n_{\text{genes}} \times p_{\text{epi}}}$, where $p_{\text{epi}}$ denotes the total number of features in the second modality. Each cell type was assigned a distinct subset of modality-specific marker features corresponding to 5% of all features. Within each subset, 10% of features were designated as high-signal core features with large projection weights, whereas the remaining features were assigned weaker background weights. All remaining entries of $W$ were set to zero, yielding a sparse and partially cell-type-specific cross-modal mapping structure. With two-modal single-cell data at hand, we generated the spatial multi-omics data using the same procedure as that in reference-matched experiments.

Using these simulated datasets, we further evaluated deconvolution performance on challenging subsets of the data, including rare cell types comprising less than 5% of cells in the reference and spatial locations with extreme sparsity levels. Specifically, spots were ranked according to their average RNA expression across five independent simulated datasets, and the top and bottom 10% were defined as low-sparsity and high-sparsity spots, respectively.

#### 4.4 Robustness experiments

**Reference data** For the robustness and generalizability analyses, we incorporated three additional scRNA-seq reference datasets alongside the aforementioned human melanoma multi-omics dataset.

Human glioblastoma scRNA-seq data The human glioblastoma (HGM) dataset comprises 28,646 cells across eight cell types, including glioma cells ($n = 14{,}422$), myeloid cells ($n = 11{,}174$), pericytes ($n = 1{,}488$), T cells ($n = 758$), endothelial cells ($n = 629$), oligodendrocytes ($n = 145$), B cells ($n = 24$), and a small group of unassigned cells ($n = 6$).

Human lymph node scRNA-seq data The human lymph node dataset comprises 2,637 cells across ten cell types, including B cells ($n = 1{,}000$), $CD4^+$ T cells ($n = 800$), $CD8^+$ T cells ($n = 400$), regulatory T cells (Treg/Tfr; $n = 200$), natural killer (NK) cells ($n = 100$), other immune cells ($n = 50$), dendritic cells (DCs; $n = 26$), myeloid cells ($n = 25$), endothelial cells ($n = 24$), and a small group of unassigned cells ($n = 12$).

Mouse spleen scRNA-seq data The mouse spleen dataset includes 11,231 cells across six cell types, including B cells ($n = 7{,}493$), memory T cells ($n = 2{,}872$), natural killer (NK) cells ($n = 321$), dendritic cells ($n = 161$), plasmacytoid dendritic cells ($n = 22$), and a small group of unassigned cells ($n = 362$).

**Data generation process** For robustness and generalization analyses, we extended the simulation framework to evaluate SMODER under external reference datasets and systematic distributional shifts. The spatial generative process and base simulation pipeline followed those described in previous sections using human melanoma single-cell RNA and epigenomic sequencing data as the reference.

Building on this base simulation, we generated perturbed datasets to evaluate robustness under controlled experimental biases. To mimic technical sparsity, RNA dropout was introduced by randomly masking nonzero entries at rates of 0.3, 0.5, 0.7, 0.8, and 0.9. To investigate the effect of spatial resolution, we varied spot cellularity by reducing the Conway–Maxwell–Poisson parameter $\lambda_r$ from the default value of 10 to 5 and 1, thereby generating spots with progressively fewer constituent cells.

To assess robustness to imperfect or mismatched references, we perturbed the scRNA-seq reference in three ways. First, rare cell types were removed by excluding the rarest 10%, 20%, and 30% of cell types. Second, reference size was reduced through stratified subsampling within each cell type at fractions of 0.20, 0.40, 0.60, and 0.80, while retaining at least one cell per type. Third, we introduced reference–target discrepancies through developmental-stage and platform perturbations. Developmental-stage shifts were simulated using cell-type-specific additive shifts combined with multiplicative scaling and variance inflation while preserving the original sparsity structure. Platform perturbations were simulated by adding count-dependent Poisson noise: $X' = X + \varepsilon$, where each entry $\varepsilon_{ij} \sim \text{Poisson}(0.04X_{ij})$.

To investigate the robustness of SMODER to spatial resolution, we generated datasets using the base simulation pipeline with two different average spot sizes: one corresponding to one cell per spot, and the other to five cells per spot.

Finally, to evaluate generalizability beyond the human melanoma reference, we incorporated three additional scRNA-seq datasets from human glioblastoma (HGM), human lymph node, and mouse spleen. For each dataset, a complementary modality was generated using the same cross-modal projection strategy described above, followed by spatial simulation using the identical pipeline. For each perturbation setting, $n = 5$ independent stochastic replicates were generated using different random seeds.

### 5 Data preprocessing

Prior to model training, the preprocessing module is employed to process the input data, ensuring data quality and consistency for subsequent model training. The detailed preprocessing procedures for each data modality are described as follows.

#### 5.1 RNA modality preprocessing

As RNA serves as the anchor modality, both spatial transcriptomics and single-cell transcriptomics data require preprocessing. Specifically, *Scanpy* was used to filter the scRNA-seq data by removing genes with zero counts across all cells and cells with zero counts across all genes by following Ma et al. [3]. Following this initial filtering, differentially expressed genes (DEGs), serving as informative features, were selected using one of two approaches based on $\log_2$ fold change ($\log_2$FC):

- selecting the top 200 genes with the highest $\log_2$FC values for each cell type; or
- retaining all genes with $\log_2$FC values greater than 1.25.

The selected DEGs were then intersected with the gene set of the spatial transcriptomics data to ensure consistency between the reference and spatial datasets.

For the scRNA-seq data, the average gene expression within each cell type is computed to construct a basis matrix, which is used in subsequent analyses. For the spatial transcriptomics data, the filtered spot-by-gene matrix $X_{\text{rna}}$ was log-normalized and subsequently used for principal component analysis (PCA) to obtain the feature matrix $F_{\text{rna}}$.

#### 5.2 Protein modality preprocessing

Protein expression data were subjected to sequential quality control and normalization. First, *Scanpy* was used to remove low-quality measurements, excluding proteins detected in fewer than 50 spots and spots with fewer than 20 detected proteins. After quality control, centered log-ratio (CLR) transformation was applied to normalize protein expression levels and mitigate compositional bias inherent to proteomic data.

##### 5.3 ATAC and histone modality preprocessing

We used the R package *ArchR* to call peaks from the ATAC fragments. For epigenomic data, including ATAC-seq and histone mark profiles, normalization and feature selection were performed to improve data quality and reliability. First, term frequency–inverse document frequency (TF–IDF) normalization was applied to account for variability in signal intensity across genomic regions. Subsequently, 20,000 highly variable peaks were selected from the normalized data and used for downstream analyses.

##### 5.4 Preprocessed scRNA-seq/snRNA-seq reference data

The processed mouse brain snRNA-seq reference dataset [7] comprises 40,532 cells and 31,017 genes across 11 major cell types, including oligodendrocytes/oligodendrocyte precursor cells (Oligo/OPC; $n = 12{,}747$), excitatory neurons in non-cortical regions (Exc-NonCtx; $n = 7{,}611$), cortical excitatory neurons (Exc-Ctx; $n = 6{,}942$), astrocytes (Astro; $n = 3{,}013$), inhibitory neurons (general subclass; Inh-General; $n = 4{,}204$), inhibitory neurons (molecular marker subtype; Inh-Marker; $n = 2{,}320$), inhibitory neurons (Meis2; Inh-Meis2; $n = 1{,}947$), microglia (Micro; $n = 1{,}218$), neuroblasts (Nb; $n = 428$), endothelial cells (Endo; $n = 22$) and other cells ($n = 1{,}324$).

The processed human breast cancer scRNA-seq reference dataset [15] comprises 100,064 cells and 29,733 genes across nine cell types, including T cells ($n = 35{,}214$), cancer epithelial cells ($n = 24{,}489$), myeloid cells ($n = 9{,}675$), endothelial cells ($n = 7{,}605$), cancer-associated fibroblasts (CAFs; $n = 6{,}573$), perivascular-like cells (PVL; $n = 5{,}423$), normal epithelial cells ($n = 4{,}355$), plasmablasts ($n = 3{,}524$), and B cells ($n = 3{,}206$).

The mouse embryo E13 scRNA-seq reference dataset [16] comprises 2,688 cells and 35,100 genes across 15 cell types, including epithelial cells of the large intestine ($n = 553$), enterocytes of the large intestine epithelium ($n = 331$), mesenchymal stem cells of adipose tissue ($n = 328$), fibroblasts ($n = 280$), endothelial cells ($n = 231$), late pro-B cells ($n = 177$), pro-B cells ($n = 171$), goblet cells of the large intestine ($n = 170$), myeloid cells ($n = 103$), leukocytes ($n = 93$), B cells ($n = 83$), neurons ($n = 61$), myofibroblast cells ($n = 42$), immature T cells ($n = 34$), and mature natural killer cells ($n = 31$).

#### 6 Details of sensitivity analysis and ablation study

**Sensitivity analysis** The model involves several key hyperparameters, including the reduced feature dimension $D$ (default: 256), the number of neighbors used in graph construction (default: 8), the embedding dimension of the graph encoders (default: 50), and the $\log_2$ fold-change threshold (default: 1.25) for selecting informative genes in the single-cell data.

To evaluate model robustness, we conducted sensitivity analyses in the simulation experiments by varying one hyperparameter at a time. Specifically, we tested the reduced feature dimension of each modality within grid $\{50, 64, 128, 256, 512\}$, the embedding dimension of the graph encoder within $\{30, 40, 50, 60, 70\}$, the number of neighbors in the kNN graph within $\{4, 6, 8, 10, 12\}$, and the $\log_2$ fold-change threshold for gene selection within $\{0.75, 1, 1.25, 1.5, 1.75\}$. By isolating each factor, we systematically quantified the sensitivity of SMODER to individual hyperparameter settings.

**Ablation study** To evaluate the contribution of each component in the SMODER objective, we performed a systematic ablation analysis by selectively removing individual loss terms. The negative log-likelihood term, $\mathfrak{L}_{NB}$, serves as the primary objective for deconvolution and is therefore retained in all configurations as the baseline. We considered three additional components: the spatial regularization loss ($\mathfrak{L}_{\text{spatial}}$), the reconstruction loss ($\mathfrak{L}_{\text{recon}}$), and the cross-modal consistency loss ($\mathfrak{L}_{\text{consist}}$). We evaluated two ablation settings: (i) removing each of the three auxiliary losses individually, and (ii) removing all pairwise combinations of the three auxiliary losses.

The ablation study was conducted on two real spatial multi-omics datasets with high-confidence annotations: (i) a mouse brain spatial CUT&Tag–RNA-seq dataset profiling H3K27ac, and (ii) a human breast cancer CytAssist RNA–protein co-profiling dataset. Performance was evaluated using the eight complementary metrics in real data evaluation, including SMS, BEAR, SC, ARI, NMI, LISI, PAS, and CHAOS. These results aimed to demonstrate that the proposed loss design is well-balanced, with complementary roles played by spatial regularization, reconstruction, and cross-modal consistency in improving deconvolution accuracy and spatial coherence.
